# Suspension of a Point-Mass-Loaded Filament in Non-Uniform Flows: Passive Dynamics of a Ballooning Spider

**DOI:** 10.1101/2020.06.06.137505

**Authors:** Moonsung Cho, Mariano Nicolas Cruz Bournazou, Peter Neubauer, Ingo Rechenberg

## Abstract

Spiders utilize their fine silk fibres for their aerial dispersal, known as ballooning. With this method, spiders can disperse hundreds of kilometres, reaching as high as 4.5 km. However, the passive dynamics of a ballooning model (a highly flexible filament and a spider body at the end of it) are not well understood. The previous study (Rouse model: without taking into account anisotropic drag of a fibre) suggested that the flexible and extendible fibres reduce the settling speed of the ballooning model in homogeneous turbulence. However, the exact cause of the reduction of the settling speed is not explained and the assumed isotropic drag of a fibre is not realistic in the low Reynolds number flow. Here we introduce a bead-spring model that takes into account the anisotropic drag of a fibre to investigate the passive behaviour of the ballooning model in the various non-uniform flows (a shear flow, a periodic vortex flow field and a homogeneous turbulent flow). For the analysis of the wide range of parameters, we defined a dimensionless parameter, which is called ‘a ballooning number.’ The ballooning number means the ratio of Stokes’ fluid-dynamic force on a fibre by the non-uniform flow field to the gravitational force of a body at the end of the fibre. Our simulation shows that the settling speed of the present model in the homogeneous turbulent flows shows the biased characters of slow settling as the influence of the turbulent flow increases. The causes of this slow settling are investigated by simulating it in a wide range of shear flows. We revealed that the cause of this is the drag anisotropy of the filament structure (spider silk). In more detail, the cause of reduced settling speed lies not only in the deformed geometrical shape of the ballooning silk but also in its generation of fluid-dynamic force in a non-uniform flow (shear flow). Additionally, we found that the ballooning structure could become trapped in a vortex flow. This seemed to be the second reason why the ballooning structure settles slowly in the homogeneous turbulent flow. These results can help deepen our understanding of the passive dynamics of spiders ballooning in the atmospheric boundary layer.

## 1 Introduction

The passive flight of spiders, which is known as ‘ballooning,’ is a unique method in animal locomotion. Spiders use their fine fibres like a sail for their aerial dispersal. It is known that some species of spiders can travel hundreds of miles and reach as high as a few kilometres above sea level. Darwin observed spiders landing while he was traveling at sea about 100 km off the Argentinian seacoast (Darwin 1845). The entomology laboratory in the United States Department of Agriculture collected a vast number of insects in the air with a biplane for five years and found a spider at the altitude of 4.5 km (Glick 1939). This passive flight of spiders is interesting not only from an ecological perspective but also considering the physical question of how point-masses (particles) can be spread widely and energy-efficiently through the air. However, the way in which fluid dynamics affect the ballooning structure is not well understood. The ballooning structure is described as a very light and thin single filament with the weight of a body at the end. (Relatively heavy spiders spin multiple silks, but we consider the simplest case first.) We refer to this as the ‘ballooning structure.’

The first fluid-dynamic approach was made by Humphrey (Humphrey 1987). He assumed a single silk ballooner and determined the possible physical dimensions (thickness and length of fibre) of a ballooner from the viewpoints of fluid dynamics, possible constraint by obstacles in nature and the mechanical properties of spider silk. However, the chart he developed could not explain the ballooning of large spiders heavier than 9 mg. Later, Suter suggested that spiders may use their legs to control their drag when they are suspended in the air (Suter 1992) and also provided possible clues regarding the chaotic motion of turbulent air flows (Suter 1999). In 2006, Reynolds showed that the flexible and extendible ballooning structure settles more slowly than that of a ballooning structure having inflexible and rigid silk (a single link ballooner). The study provided rational insight into spiders’ ballooning in nature. However, the simulation needs some improvements. First, the drag anisotropy of a fibre structure should be considered for the simulation of fibre dynamics because various filament-like structures in nature, such as cilium, flagellum and pappus, experience and generate anisotropic drag forces (Purcell 1977, Childress 1981, Happel and Brenner 1986). Second, the rigid silk model in the simulation is described as a dumbbell model. It is difficult to regard such a structure as a rigid silk in turbulent flow. Third, the simulation considers a spider silk to be flexible and extendible. The assumption of flexibility is reasonable; however, the assumption of extendibility should be re-examined for the following reason. From a simple calculation, the extendibility of a spider silk during flight is almost negligible because the 0.5-mg spider weight (either the case of a small spider on a single fibre or the distributed weight of a large spider, which uses tens of fibres) extends 47 mm in length (1.5% elongation; elastic modulus: 10 Gpa; Gosline et al. 1999) in the case of a spider silk that is 3 m long and 200 nm thick (Cho et al. 2018). On the other hand, the study provides ‘economical’ insight regarding methodology. Although Zhao et al. recently implemented a very detailed numerical simulation using an immersed boundary method, the proposed simulation was limited to two-dimensional flow and the lengths of spider silk were limited to less than 20 cm (Zhao et al. 2017). If we introduce the chain-like fibre and the unsteady-random-Fourier-modes-based turbulence model, the simulation time can be enormously shortened and various combinations of physical parameters can be investigated with the simulation.

From the view of physics, the ballooning structure has three unique features. First, two different flow regimes are involved in its dynamics. As a ballooning silk exhibits the feature of extremely high slenderness (16.1 × 10^7^, Cho 2018), the structure has two different length scales (thickness: 200 nm; length: 3.22 m; Cho 2018). Therefore, one unique feature is that a small segment of spider silk is dominated by a low Reynolds number flow, while the behaviour of the entire silk is governed by a high Reynolds number flow, a turbulent flow. The upper limit of the Reynolds number is about 0.04 (*Re* = *ρVd*/*μ*; air density: 1.225 kg m^−3^; maximum possible velocity: 3 m s^−1^; thickness of spider silk: 211 nm; dynamic viscosity of air: 1.837 × 10^−5^ kg m^−1^ s^−1^). Second, the ballooning silk in turbulent wind is exposed to different wind vectors, unlike aerial seeds that are exposed to quasi-homogeneous (uniform) flow fields because of their small sizes. Whether or not the ballooning structure gains energy from this non-uniform flow field is an interesting question. Third, asymmetric fibre characteristics appear because of the weight of the body at the end of the fibre. As the lower end of a fibre is constrained by the weight (body), the upper part of a fibre can be distorted and adapted more easily than the lower part. How an asymmetrically constrained fibre behaves in a non-uniform flow and what influences this dynamic can also be interesting questions concerning the ballooning flight of spiders.

Fibre suspension in low Reynolds number flow has been frequently dealt with in polymer dynamics (Yamamoto and Matsuoka 1993, Ning and Melrose 1999, Schroeder et al. 2005, Delgado-Buscalioni 2006, Gerashchenko and Steinberg 2006, Winkler 2006, Lindström and Uesaka 2007, Das and Sabhapandit 2008, Delmotte et al. 2015) and in the area of DNA stretching (Bustamante et al. 1994, Perkins et al. 1994, 1995, Jian et al. 1997, Larson 1997, Bustamante et al. 2000, Wong et al. 2003, Schroeder et al. 2004, Lee and Thirumalai 2004, Shaqfeh 2005, Wang and Lu 2007, Jo et al. 2009, Dai et al. 2014). However, the applied flows in these studies were not turbulent flow, but rather a simple shear flow or Brownian motion of solution molecules. Recently, the motion of a fibre in turbulent flow has been given attention because of its anisotropic shape and has been studied by experiment (Brouzet and Le Gal 2014, Verhille and Bartoli 2016), while the Reynolds number, which the local segment of a fibre experiences, is in the range of a moderate Reynolds number flow, about *Re* ≈ 4000, rather than a low Reynolds number flow, *Re* < 0. Asymmetric condition of a fibre is mostly studied as a tethered fibre (Doyle et al. 2000, Ladoux and Doyle 2000, Ibáñez-Garcıá and Hanna 2009, Litvinov et al. 2011). Suspension of a flexible fibre under asymmetric conditions was first simulated by Reynolds et al. (2006) for the study of ballooning dynamics. The study points out that the extendibility and flexibility of a filament may be the causes of the settling retardation. However, as discussed above, the model only considered the isotropic drag of a fibre (Rouse model; Rouse 1953, Dhont 2003, Reynolds at al. 2006, Doi and Edwards 2007), not anisotropic drag.

There are two representative aspects to the study of ballooning. One is from the viewpoint of fluid dynamics, which is a primary factor in the passive aerial dispersal of animals and plant seeds (Humphrey 1987, Suter 1991, 1992, 1999, Reynolds at al. 2006, Zhao et al. 2017, Cho et al. 2018). The other is from the viewpoint that tries to find clues from the earth’s electric field (Gorham 2013, Morley and Robert 2018). In the fluid-dynamic approach, the atmospheric turbulent flow has been examined as an influential factor for high aerial dispersal capability (Suter 1999, Reynolds et al. 2006, Zhao et al. 2017) because flexible spider silks are deformed by turbulent flows and fall slowly (Reynolds et al. 2006). However, the question of how the physical mechanism behind this works has not yet been answered. In the earth’s electric field approach, moreover, it has recently been reported that the electric field induces spiders’ pre-ballooning (tiptoe) behaviour (Morley and Robert 2018). Despite this, we are still on the way to understanding the electrical properties of spider silk in many electrical conditions and the background physics of electrical flight (Vollrath and Edmonds 2013, Ortega-Jimenez and Dudley 2013, Kronenberger and Vollrath 2015, Joel and Baumgartner 2017).

In this study, we focus on the understanding of the suspension character of the ballooning structure in non-uniform flows from the viewpoint of fluid dynamics. To describe a flexible fibre, we employ a bead-spring model, taking into consideration the hydrodynamic interaction between beads (Zimm model, which can describe the anisotropic drag of a fibre). There are two major purposes for this study. First, we want to clarify the role of anisotropic drag of a fibre in spiders’ flight. Second, we want to learn whether or not this fibre induces preferential motion of a ballooning structure in homogeneous turbulent flow.

## 2 Methodology

### 2.1 Modelling

Due to the extreme thinness and long length of each fibre, high computational performance and extensive time are required to solve the dynamics of this flexible thin filament. Therefore, we employ a relatively simple method, the bead-spring model, to describe a flexible filament and simulate the motion of a ballooning structure in various non-uniform fluid flows. Owing to its rapid calculation capacity, this model offers a significant advantage for the investigation of various simulation parameters. For the investigation of the effect of an asymmetric mass constraint on a fibre, several things are assumed: (i) a single silk ballooner; (ii) ignorance of the effect of the body size (the diameter of a spider body is equal to the thickness of a silk); (iii) no consideration of the inertia of the spider body. To simulate the dynamics of a highly-slender filament with the bead-spring model, which has a relatively small slenderness of 45-120, a non-dimensional parameter is defined and an artificial fluid model is applied, the motion of which resembles that of homogeneous turbulence of the air but has high viscosity. These treatments enable the incorporation of the two different flow regimes in a single model, and the universal analysis under the condition of different scales of the structure and various properties of fluids is possible.

#### 2.1.1 Bead-Spring Model

The bead-spring model (Zimm model) is frequently used to approach the polymer physics and dynamics of a microswimmer (Doi and Edwards 1986, Yamamoto and Matsuoka 1993, Gauger and Stark 2006, Lindström and Uesaka 2007). Each bead is connected to the other with a tensile and bending spring. By adjusting each stiffness value, we can simulate various extendibility and flexibility values of a filament (see Figure 1). Moreover, each bead experiences hydrodynamic viscous drag in a low Reynolds number flow, which is governed by Stokes’ law (Happel and Brenner 1986). The Zimm model considers hydrodynamic interactions between beads. This consideration describes the anisotropic drag of a thin filament in a low Reynolds number flow, which shows the drag difference between a transverse drag and a longitudinal drag (Purcell 1977, Childress 1981, Happel and Brenner 1986, Gauger and Stark 2006). The introduction of the hydrodynamic interaction is the main difference of this study compared to the previous simulation (Reynolds et al. 2006).

**Figure 1.**
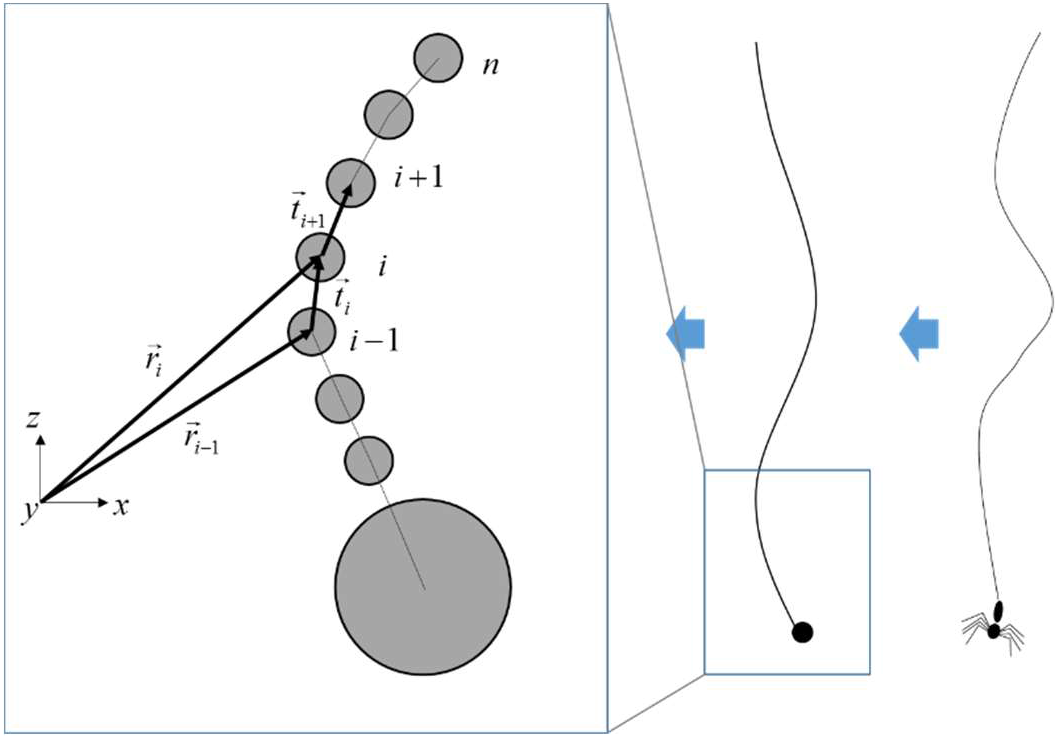
Bead-spring model for a ballooning spider.

The dynamics of each bead is governed by the flow field and forces on the bead; a gravity force 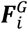, a stretching force of a spring 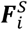, and a bending force of a spring 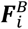 (see Equation (2.1); *i* refers the *i*-th bead). ***μ***_*ij*_ is a mobility tensor that transforms external forces into the mobility components of the *i*-th bead. The details for the modelling are found in S.M. (Supplementary Material) S.1.1 and Gauger and Stark 2006.

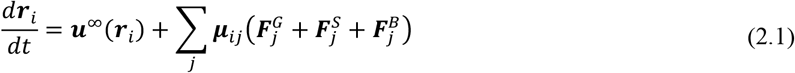

#### 2.1.2 Turbulent Model

The chaotic motion of flows is modelled by Fung’s kinematic simulation, which describes the turbulent flow with multiple Gaussian random velocity fields (see Equation (2.2) and Figure 2; Kraichnan 1970, Drummond et al. 1984, Turfus 1985, Fung et al. 1992). ***a***_*n*_ and ***b***_*n*_ refer amplitude vectors of the *n*-mode. ***k***_*n*_ is the wavenumber vector of the *n*-mode. ω_*n*_ is the frequency of the *n*-mode. ***u***, ***r***, *t*, and *N* refer velocity vector, position vector, time, the maximum number of modes, respectively. Determination of the values is explained in the S.M. S.1.2. This model is used to simulate turbulent flows for various purposes: for example, the study of turbulent structures (Fung 1990, Vassilicos and Fung 1995, Fung and Perkins 2008, Suzuki and Sakai 2013, Lafitte et al. 2014), investigation of particle dynamics (Fung 1993, Fung 1998, Fung and Vassilicos 1998, Fung and Vassilicos 2003, Gustavsson et al. 2012) and so forth. The turbulence model is intended to be homogeneous and to show the von Karman energy spectrum, which can describe grid-generated turbulence and the spectrum in natural wind (Harris 1971, Bearman 1972, Hunt 1973).

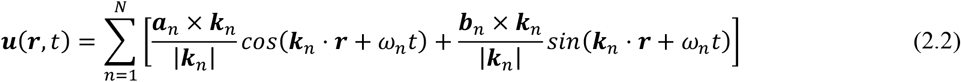

**Figure 2.**
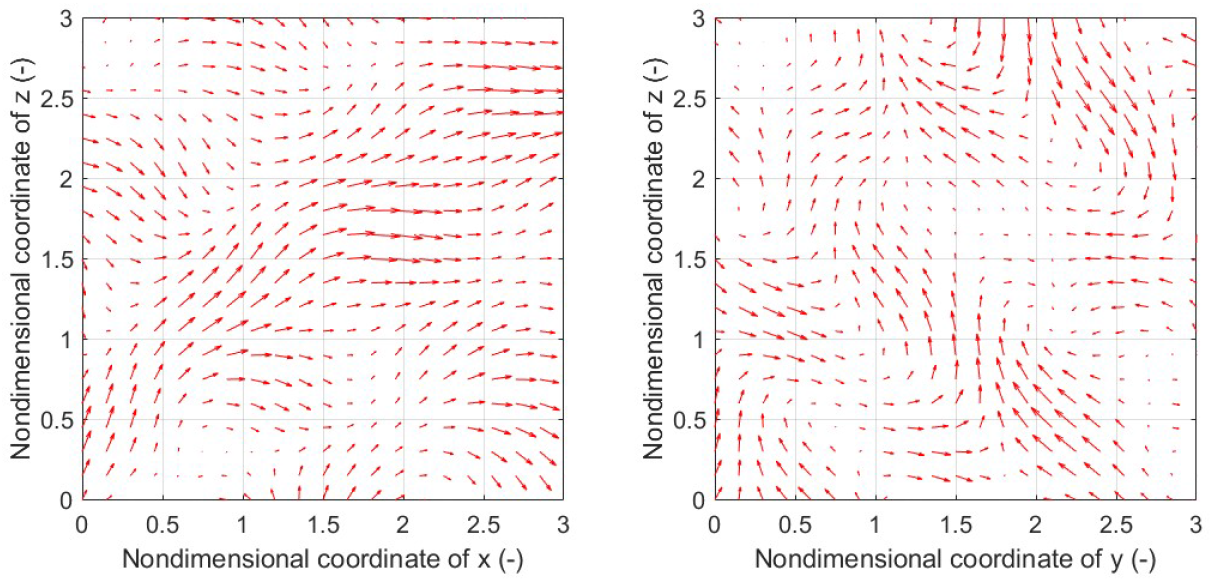
Cross-sectional velocity fields of the 3-dimensional homogeneous turbulence.

#### 2.1.3 Shear Flow Model

In order to investigate the sediment characteristic in a non-uniform flow, a shear flow model is used. Only horizontal components exist in a shear flow. Therefore, it is a good model to investigate the sediment characteristic (vertical motion) of a ballooning structure in a non-uniform flow. Shear flow can be described as in Equation (2.3) (*γ* refers to shear rate, which has units of *s*^−1^, while z is the vertical coordinate; see Figure 3):

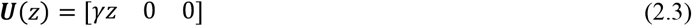

**Figure 3.**
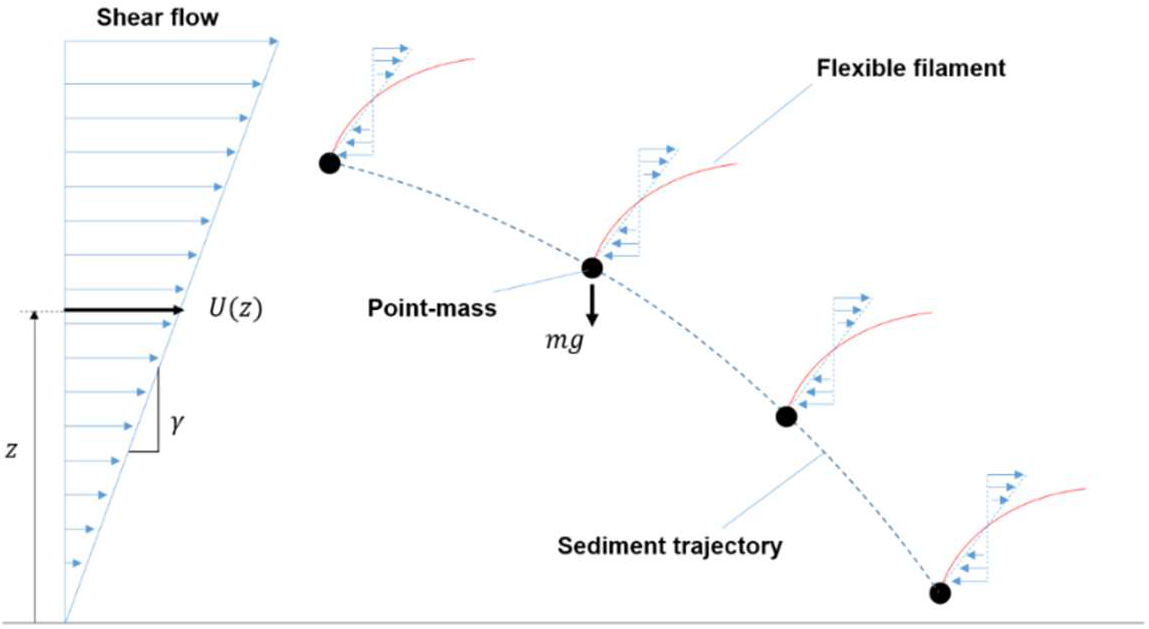
Schematic of the simulation in a shear flow.

To figure out the physical cause of settling speed reduction of a ballooning structure in a shear flow, the decomposition of the relative flow velocities ***U***′_*j*_ on beads is executed into the vertical and horizontal components of the relative flow velocities (see Equation (2.5) and Figure 4). The vertical resistance force due to the pure geometric shape of a filament 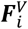 is calculated from the vertical components of the relative velocities on beads 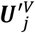 (see Equation (2.6)). Moreover, the vertical resistance force that is caused by the pure horizontal shear flow 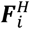 is derived from the horizontal components of the relative flow velocities 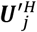 (see Equation (2.6)).

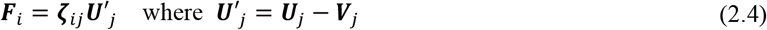

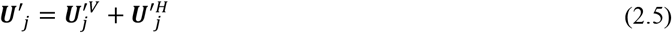

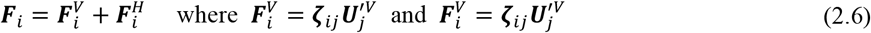

**Figure 4.**
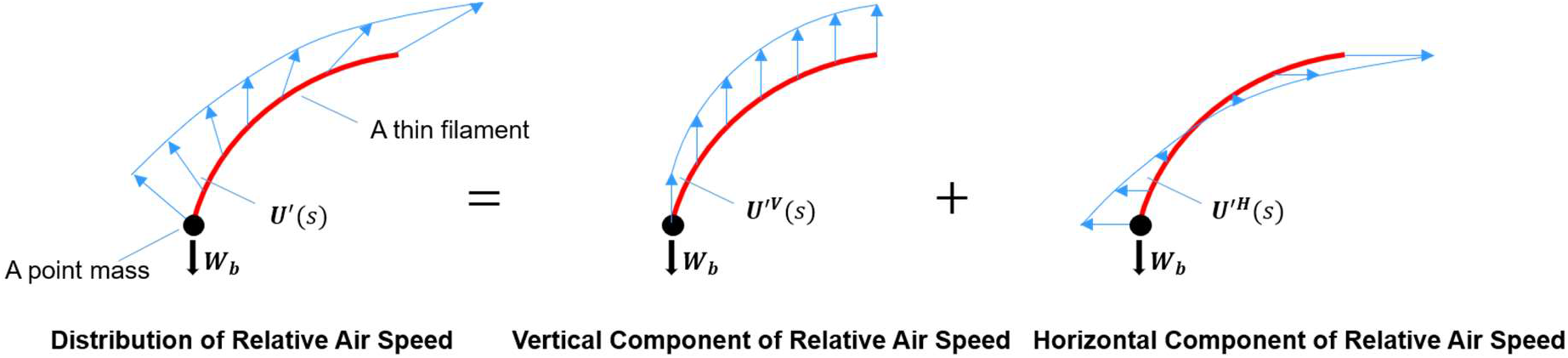
Schematic diagrams showing the decomposition of relative flow velocities on the ballooning structure in a shear flow.

The vertical net forces on the bead spring model are calculated by summating the drag forces on each bead (see Equations (2.7) and (2.8)).

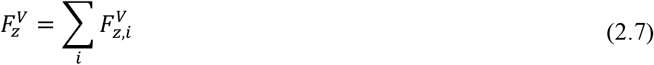

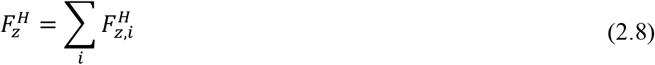

Once the forces have been calculated, the non-dimensional vertical resistance coefficients caused by pure geometric shape and the horizontal shear flow can then be calculated by dividing the vertical components of forces by the viscosity, settling speed and silk length (see Equations (2.9) and (2.10) and Figure 4):

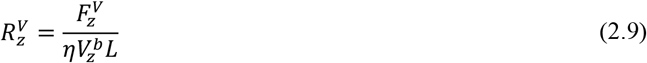

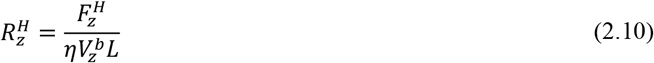

#### 2.1.4 Periodic Cellular Flow Model

The periodic cellular flow model is used to study the settling behaviour of a particle (Maxey and Corrsin 1986, Maxey 1987a, Fung 1997, Bergougnoux et al. 2014). The cellular flow field was first used by Stommel to investigate plankton suspension in Langmuir circulation and has since been frequently used for the study of particle sediment in turbulent flow (Stommel 1949, Maxey and Corrsin 1986, Maxey 1987a, Fung 1997, Bergougnoux et al. 2014). The flow field can be described by means of two-dimensional convection cells (see Figure 5). The flow is two-dimensional, incompressible and steady and specified by a stream function ψ (see Equation (5.22)). *U*_*O*_ and *L*_*c*_ are maximum velocity in the flow field and size of a convection cell, respectively. The velocity field of the cellular flow can be expressed by spatially differentiating the given stream function ψ (see Equations (5.23) and (5.24)).

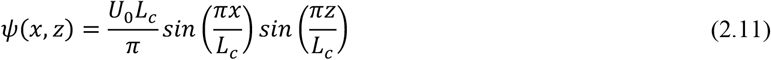

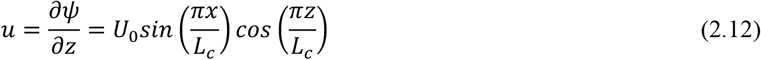

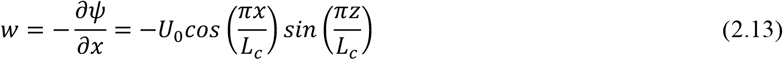

**Figure 5.**
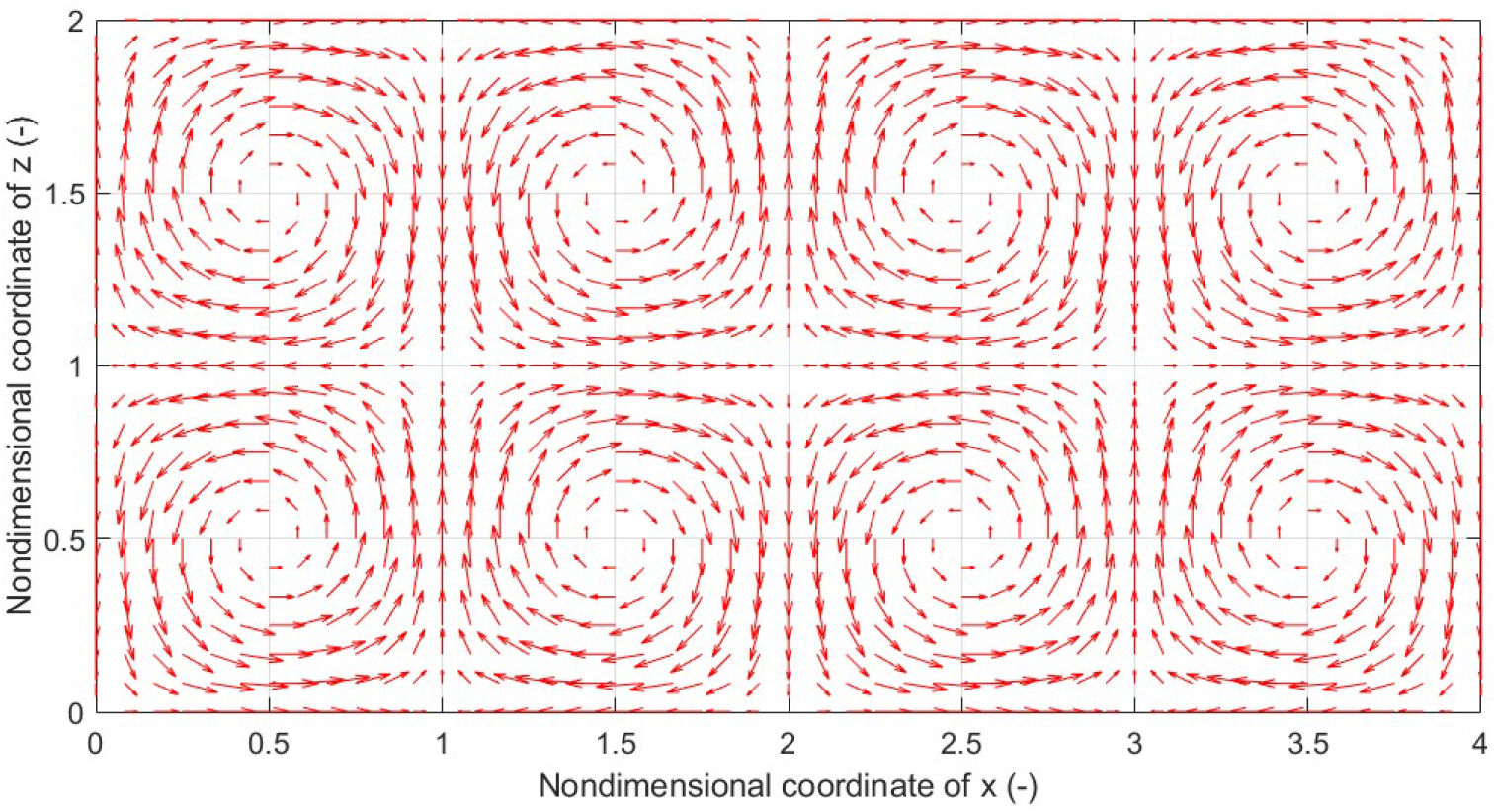
Velocity field of a periodic cellular flow model at *U*_*0*_ = 1 and *L*_*c*_ = 1

### 2.2 Non-Dimensional Parameter

In order to consider a wide range of physical scales, we define a non-dimensional parameter, referred to as the ‘ballooning number,’ by comparing the Stokes force on a filament to a gravity force on the weight, that is, a spider’s body at the end of a slender and flexible filament (see Equations (2.14) and (2.15)). Here, we do not consider the thickness of the filament because the influence of filament thickness on the fluid-dynamic force in a low Reynolds number flow is relatively much smaller than that of filament length. The Stokes force on a filament is described by multiplying the viscosity, the characteristic velocity of fluid-flow and the length of a filament. The gravity force on a weight is defined as the mass of the weight multiplied by a gravity acceleration, 9.8 *m*⁄*s*^2^. The characteristic velocity can be defined in various ways according to the flow conditions. For a shear flow, it is defined as the value of the shear rate multiplied by the length of the filament, *U* = *γL* (see Equation (2.16)). In the case of a cellular flow, moreover, the characteristic velocity is the maximum velocity in the flow *U*_*0*_ (see Equation (2.17)). For homogeneous turbulence, the characteristic velocity is the root-mean-square fluctuation velocity σ (see Equation (2.18)):

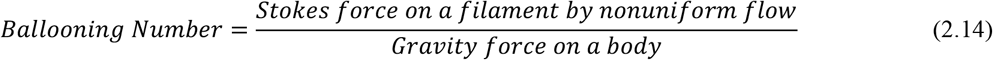

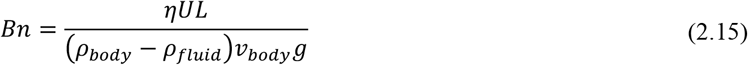

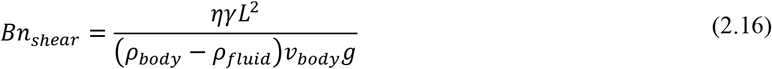

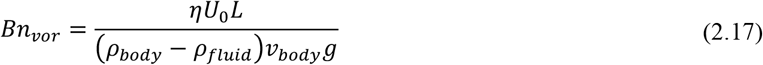

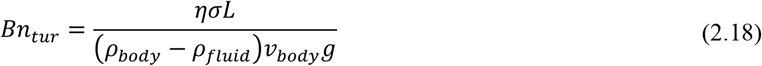

### 2.3 Execution

The motion of the bead-spring model is numerically integrated using the Euler method. The integration of the simulation in a shear flow is performed until the simulation reaches the steady state. The execution time for the simulation in a periodic cellular flow and in homogeneous turbulence are defined by comparing the settling speed of the model in the still fluid medium and the strength of a cellular flow *U*_*0*_ and the root-mean-square fluctuation velocity of the homogeneous turbulence σ, respectively. Twenty times longer than the period of the one characteristic time-revolution is used as simulation time. The detailed definitions of the one characteristic time-revolution are found in S.M. S.1.3.

## 3 Results

### 3.1.1 Homogeneous Turbulence

Figure 6 shows an example of the simulation and the dimensionless mean settling velocities for different ballooning numbers. The horizontal axis represents the ballooning number for homogeneous turbulence. The vertical axis is referred to as non-dimensionalised settling speeds that are divided by the settling speed of the structure in still medium. Therefore, the dimensionless settling speeds that are greater than 1 indicate fast falling (faster than the settling speed in still medium), while the dimensionless settling speeds that are between 0 and 1 indicate slow settling events caused by external flows. Negative dimensionless settling speeds mean rising motion of the structure. As the strength of flow fluctuation becomes more dominant, the ballooning structure exhibits a slow settling character (see Figure 6B).

**Figure 6.**
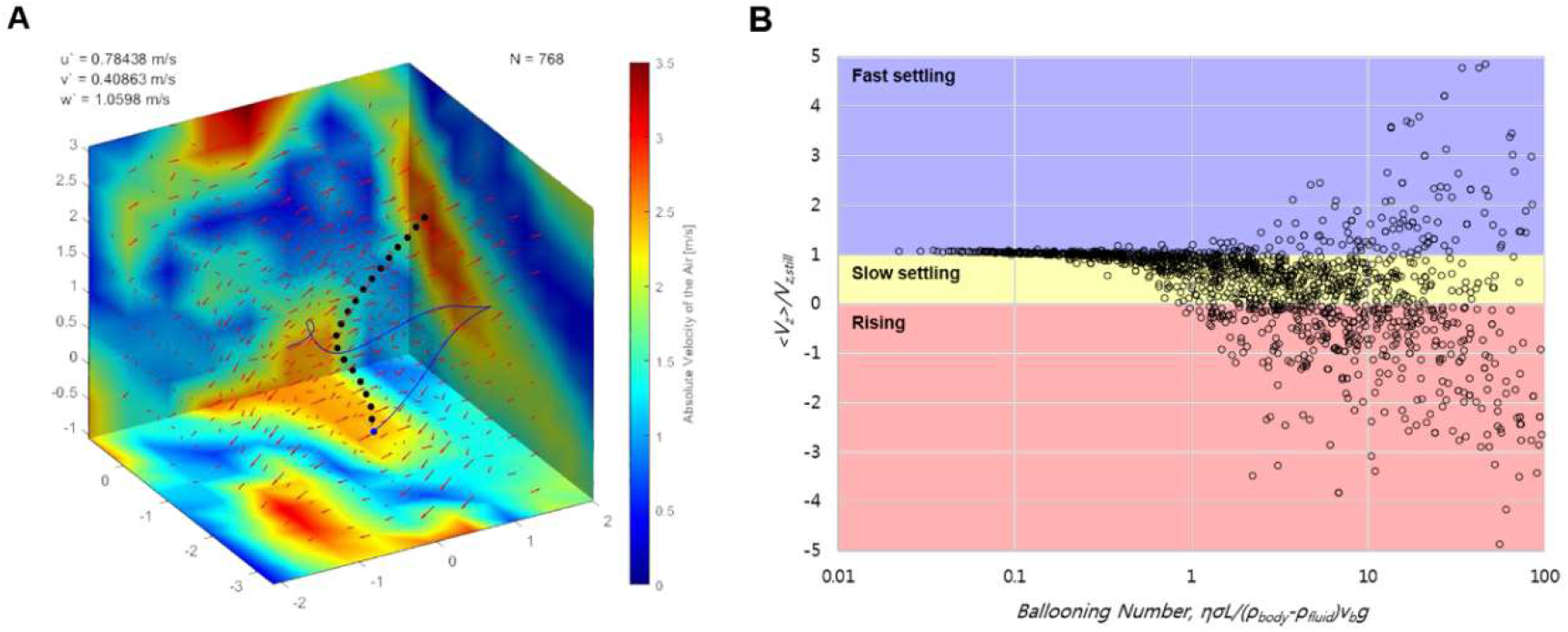
(A) Simulation of the filament-like structure in the homogeneous turbulence. The bold points describe the bead-spring model. The blue point is the bead on which the point mass is applied. The blue solid line is the trajectory of the point mass. The colour bar represents absolute velocities of the flow. *σ*_*u*_, *σ*_*v*_ and *σ*_*w*_ denote the standard deviations of the *x*, *y*, and *z* components of velocity until the nth iteration. (B) Dimensionless settling speed distribution of the ballooning structure in the homogeneous turbulence model according to the ballooning number. A total of 1400 cases of combination of the parameters are simulated and the averaged settling speeds are plotted. Fast settling: the structure falls faster than it would in still air. Slow settling: the structure falls more slowly than it would in still air. Rising: the structure does not fall but rises upward.

An end-to-end vector, 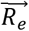, represents the statistical shape of the ballooning structures during their suspension in turbulence (see Figure 7E). The averaged z component of the end-to-end vector approximates 1 at the ballooning number (see Figure 7AB and S.M. Figure S5 (f)). The value decreases as the ballooning number increases (see Figure 7A to 7D).

**Figure 7.**
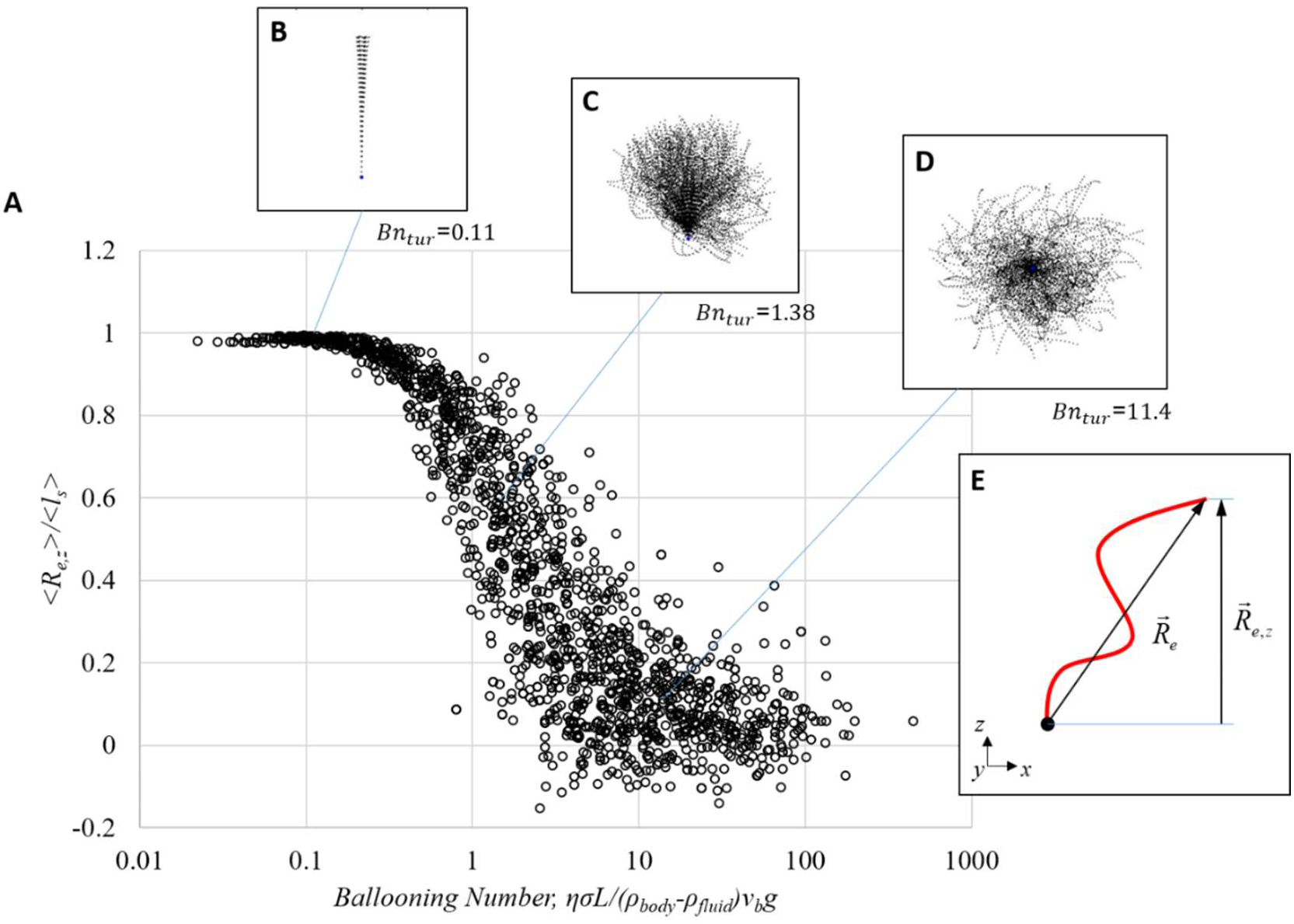
(A) Distribution of the mean vertical (z-direction) components of end-to-end vectors according to the ballooning number for homogeneous turbulence. A total of 1400 cases of combination of the parameters are simulated. (B)-(D) Superposed shapes of ballooning silks during the flight for different ballooning numbers. The reference point (blue dot) is the position of its body. (E) Schematic diagram of an end-to-end vector and the z component of the end-to-end vector.

### 3.1.2 Shear Flow

The vertical velocity difference in a shear flow deforms the filament structure in its sedimentation. As the shear rate increases, the deformation becomes larger, and the deformed geometry in the shear flow affects the sediment velocity of the structure (see Figure 8).

**Figure 8.**
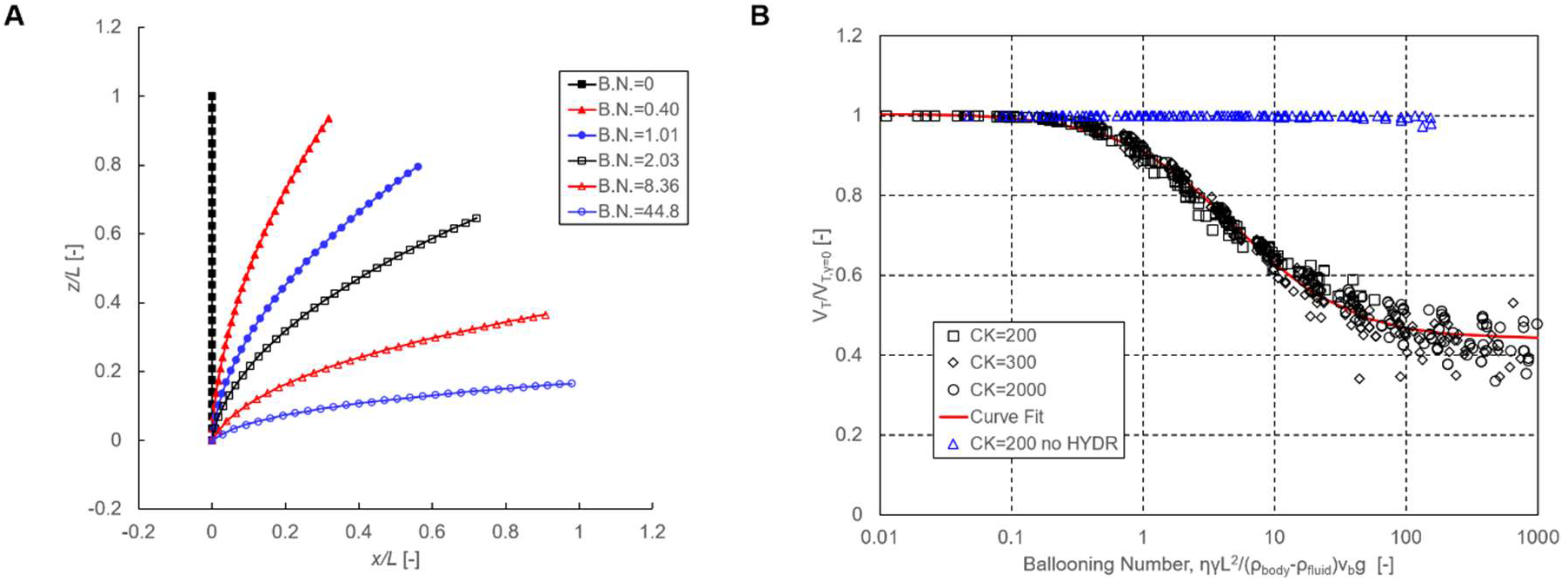
(A) Shapes of a filament in various strengths of a shear flow (various ballooning numbers). (B) Variation of dimensionless settling velocity of the ballooning structure in a shear flow with variation of the ballooning number. Black marks (720 cases) indicate the cases with hydrodynamic interaction (considering the anisotropic drag of the filament), while blue marks (138 cases) are the cases without hydrodynamic interaction (considering the isotropic drag of the filament). CK refers to the spring constants between beads.

The influence of the anisotropic drag is studied by comparing two results, namely, those with and without hydrodynamic interaction. As the dimensionless shear rate (ballooning number for shear flow *Bn*_*she*_) increases, the ballooning structure stretches horizontally (see Figure 8A), and the settling speed is reduced (Zimm model). These results are not seen in cases in which the hydrodynamic interaction is ignored (considering the isotropic drag of a fibre, Rouse model; see Figure 8B). The settling speed is reduced by 40% at the high ballooning number (*Bn*_*shear*_=1000).

The parameter most associated with settling speed is the resistance coefficient of the object. The resistance coefficient of a filament-like object is expressed by the vertical drag force divided by the settling speed and the length of the filament.

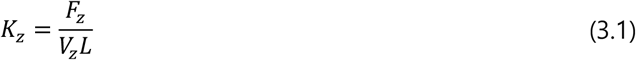

To elucidate the causes of the reduction of the settling speed, the non-dimensional resistance coefficient is decomposed into geometric (see Equation (2.9)) and shear-induced (see Equation (2.10)) non-dimensional resistance coefficients.

Figure 9 shows the variation of the dimensionless resistance coefficient. As the ballooning number increases, so too does the total non-dimensional resistance coefficient. This coincides with the fact that the non-dimensional settling speed decreases as the ballooning number increases (see 8B). The increase of the non-dimensional resistance coefficient at the beginning, 0.1 < *Bn*_*shear*_ < 3, is mostly caused by the geometric shape of a filament, which is stretched horizontally and falls in a transverse attitude (see red triangular marks in Figure 9). At ballooning numbers over 3, the increase of the non-dimensional resistance coefficient is additionally induced by a shear flow (see blue circular marks in Figure 9).

**Figure 9.**
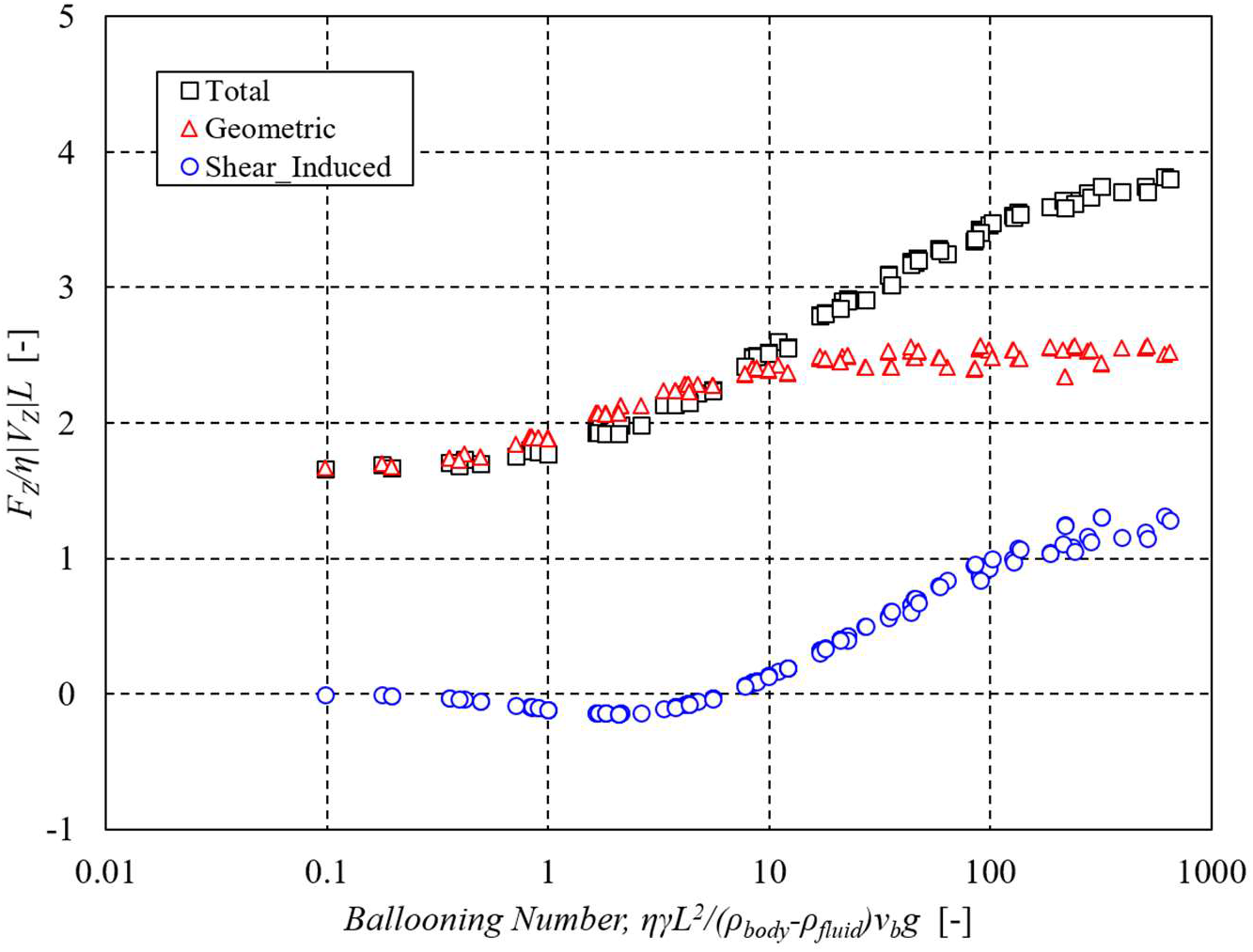
Variation of dimensionless resistance coefficients with the ballooning number for a shear flow. The dimensionless resistance coefficient is decomposed into geometric coefficient and shear-induced coefficient. A total of 128 cases for different ballooning numbers are simulated.

### 3.1.3 Vortex Field

The ballooning structure exhibits various behaviours in a cellular flow field. According to the size and strength of the vortices, the ballooning structure exhibits the following motions: (i) ‘vortex crossing’; (ii) ‘fast settling’; (iii) ‘slow settling’; and (iv) ‘vortex trapping.’ At low ballooning numbers (*Bn*_*vor*_ < 2), the ballooning structure falls crossing the vortices. The influence of flow eddies on their settling speed are small; therefore, their dimensionless settling speeds are around 1 (0.9 ≤ 〈*V*_*z,last*_〉/*V*_*z,still*_ < 1.1, Figure 11 (a)(d)(f), and S.M. Figure S11, Figure S16). As the ballooning number increases, the motions split into three different behaviours. The split depends on the size of the vortices and the initial positions. If the sizes of the eddies are under 1.2 times the silk length, the ballooning structure falls faster than it would in still air (see Figure 10, Figure 11 (e)(h)(i) and S.M. Figure S17, Figure S20, Figure S23, Figure S26); if the eddies are larger than 1.2 times the silk length, however, the ballooning structure falls more slowly than it would in still air (see Figure 10 and Figure 11 (b)). At higher ballooning numbers, the ballooning structure is trapped by an eddy and the mean settling speed approaches zero (see Figure 10 and Figure 11 (c)(f); see S.M. Figure S21, Figure S22, Figure S27 and Figure S32).

**Figure 10.**
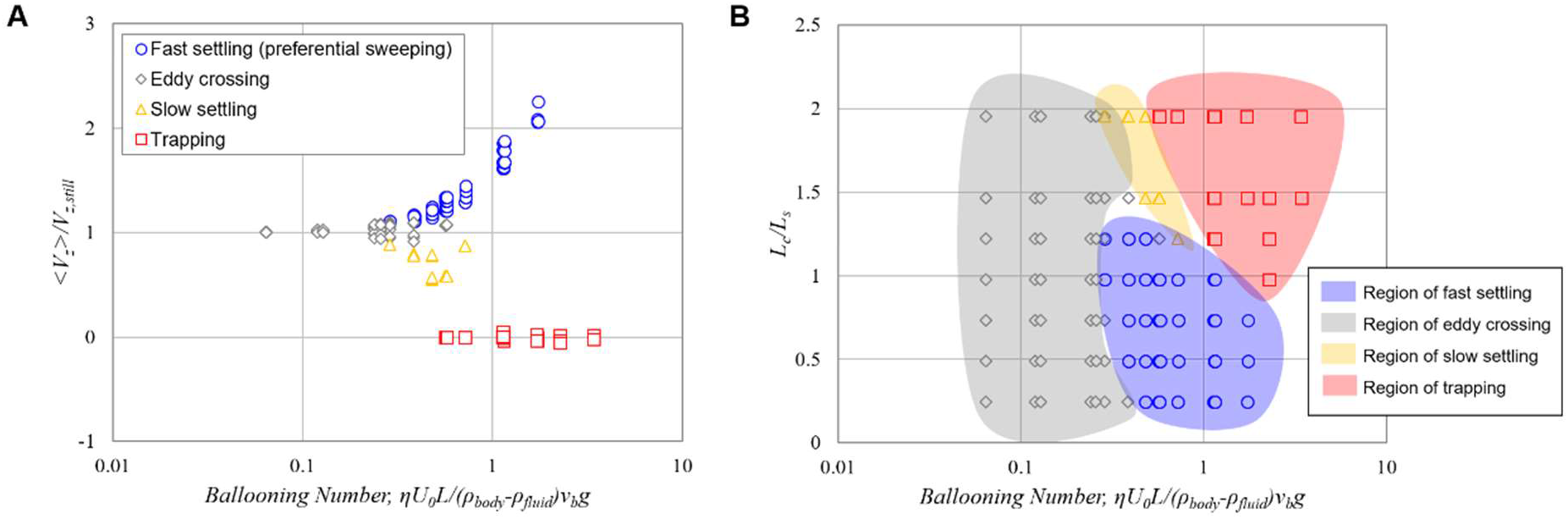
(A) Dimensionless settling speed of the ballooning structure in a cellular flow field. (B) Different behavioural modes according to eddy size and ballooning number. The behavioural modes are categorized by the motion and settling speed of the structure. ‘Eddy crossing’ is the mode in which their dimensionless settling speeds are between 0.9 and 1.1; ‘slow settling’ is where speeds are slower than 0.9; ‘fast settling’ is the mode that falls faster than 1.1. The ‘trapping motion’ is recognised by detecting the averaged settling speed of 0 (or near 0). The legend of (A) is valid in the figure of (B).

**Figure 11.**
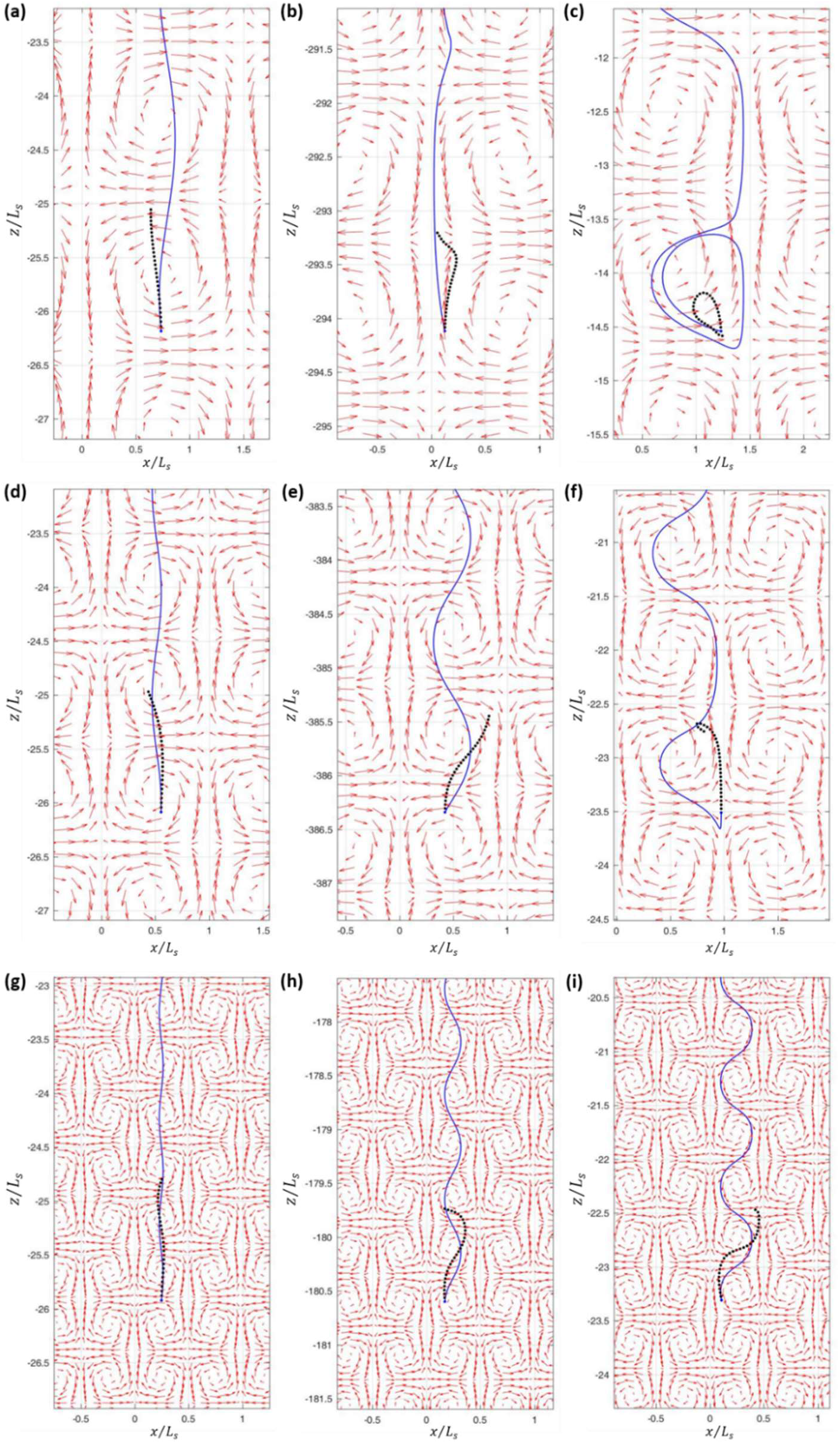
Trajectory of the ballooning structure in a cellular vortex field. (a)(d)(g) Eddy crossing: the ballooning structure falls crossing the eddies; (b) Slow settling: the ballooning structure stays between two vortex columns and falls more slowly than the structure in still air; (c)(f) Trapping: the ballooning structure is trapped by a vortex after a few or many periodic sediment motions; (h)(i) Fast settling (preferential sweeping): the ballooning structure deforms and passes through a curved path between the vortices, and the structure falls faster than it would in still air.

Some interesting features emerging from these results include the fact that the ballooning structure is preferentially trapped by a vortex in certain flow conditions (high ballooning number and in large vortex cells). The structures travel downwards through the high-strain region between vortices; then, after a few oscillations in sediment progress, the structure becomes trapped in a vortex (see S.M.: Figure S27, Figure S29, Figure S31). The distances travelled by sediment before the trapping events depend on its initial positions, where those are released (x direction). However, at the end, all structures were trapped in a vortex. This suggests that a biased character exists in the suspension of the ballooning structure in vortex fields: namely, the ballooning structure is preferentially trapped by a vortex at high ballooning numbers. Moreover, two trapping motions can be observed in the events: cycling trapping, which shows a closed loop trajectory of the body (see Figure 12A), and stationary trapping, which converges to the equilibrium states in the position and shape of the filament (see Figure 12B).

**Figure 12.**
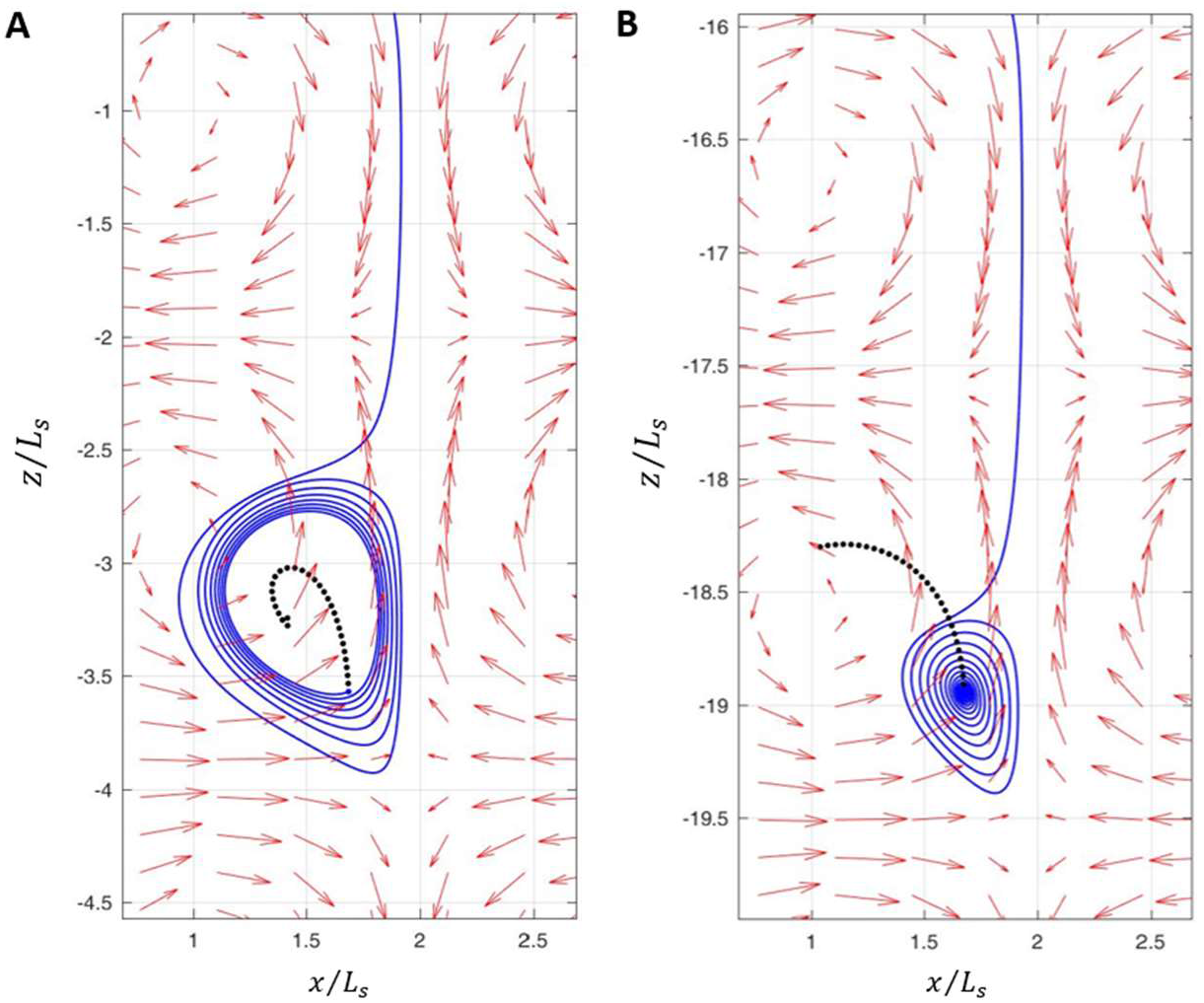
Two different trapping modes in a cellular vortex field. (A) Cycling trapping building a closed loop; (B) Stationary trapping remaining in an equilibrium state.

### 3.1.4 Energy Extraction with a Filament-like Structure

The sediment behaviour of a ballooning spider is compared with a point-like parachuter, which represents dandelion and thistle seeds. The size of the point-like parachuter is adjusted to give it the same settling speed as that of the ballooning spider in still medium (see Figure 13); therefore, both structures have the same weights and settling speeds in still air. Both are released from identical locations in the identical turbulence model.

**Figure 13.**
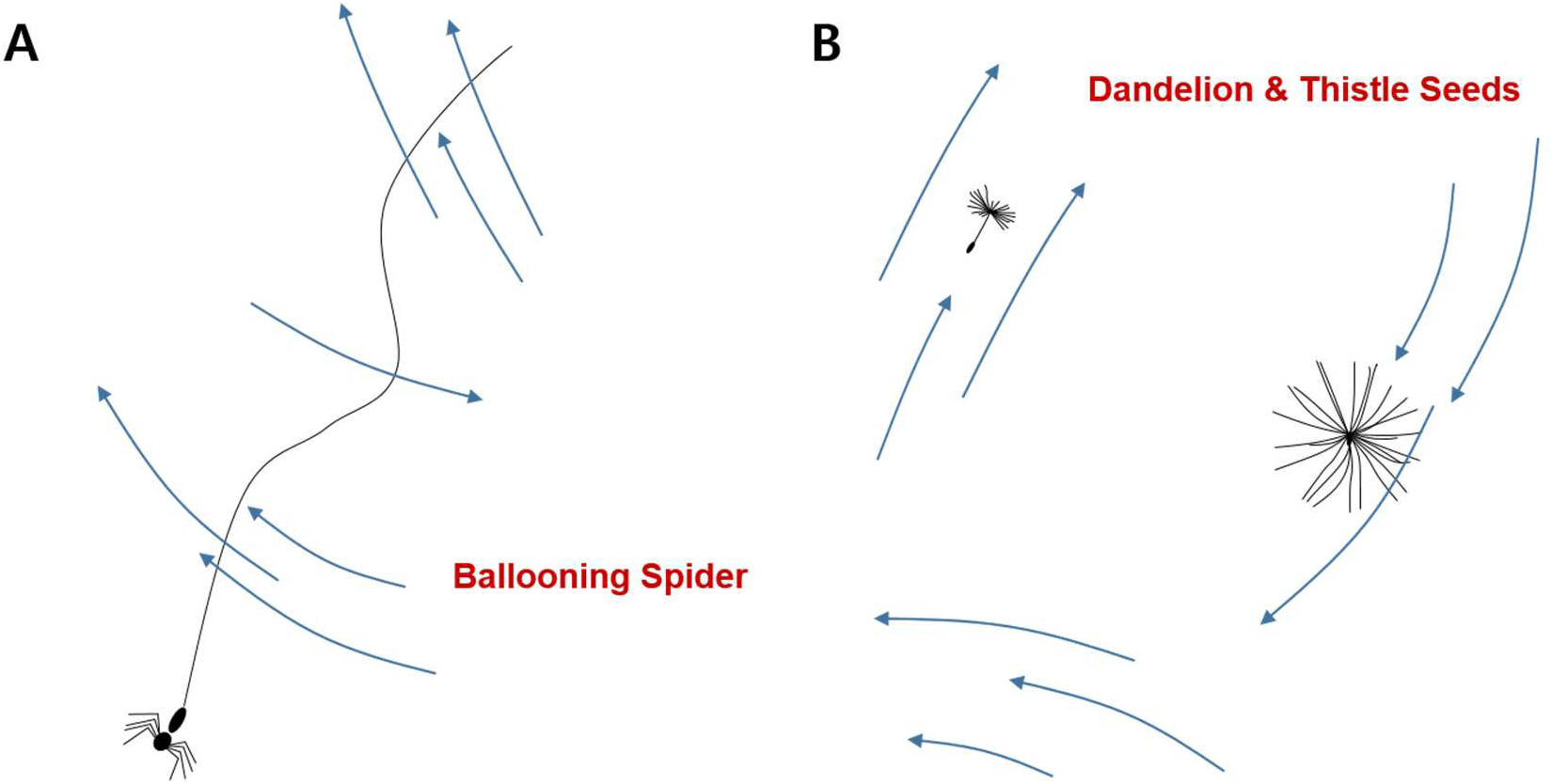
Comparison between the flight of a ballooning spider and the flight of a dandelion (or thistle) seed. (A) Sketch of a spider’s ballooning; (B) Sketch of the flight of dandelion and thistle seeds.

While the point-like parachuter showed a mirror-symmetric distribution about 〈*V*_*z*_〉/*V*_*z,still*_ = 1 (see Figure 14B), the filament-like parachuter showed a biased distribution about 〈*V*_*z*_〉/*V*_*z,still*_ = 1 (see Figure 14A). Figure 15 clearly shows the biased motion of the filament-like parachuter in comparison with the point-like parachuter.

**Figure 14.**
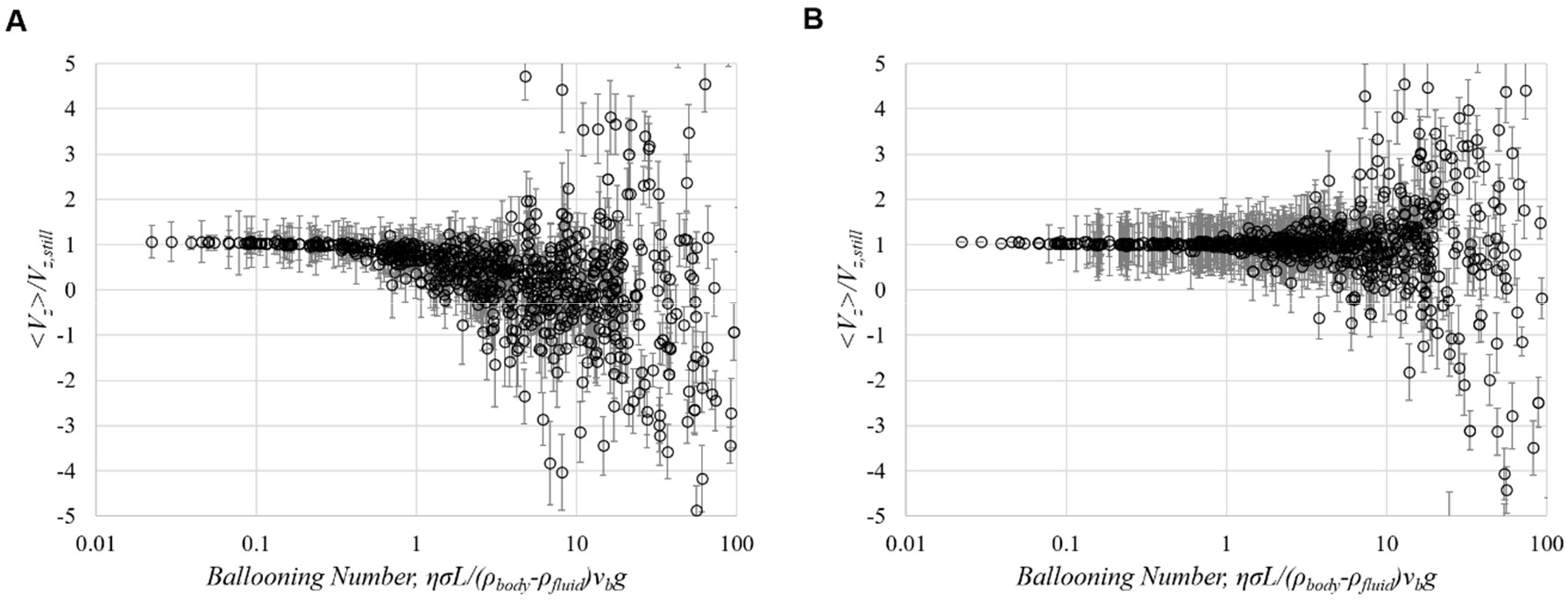
Comparison between a filament-like parachuter and a point-like parachuter under dimensionless settling speed. A total of 750 cases are simulated and the averaged settling speeds with their standard deviations are plotted. (A) Dimensionless settling speed (divided by the settling speed in the still medium) of a filament-like parachuter in homogeneous turbulence; (B) Dimensionless settling speed (divided by the settling speed in the still medium) of a point-like parachuter in homogeneous turbulence.

**Figure 15.**
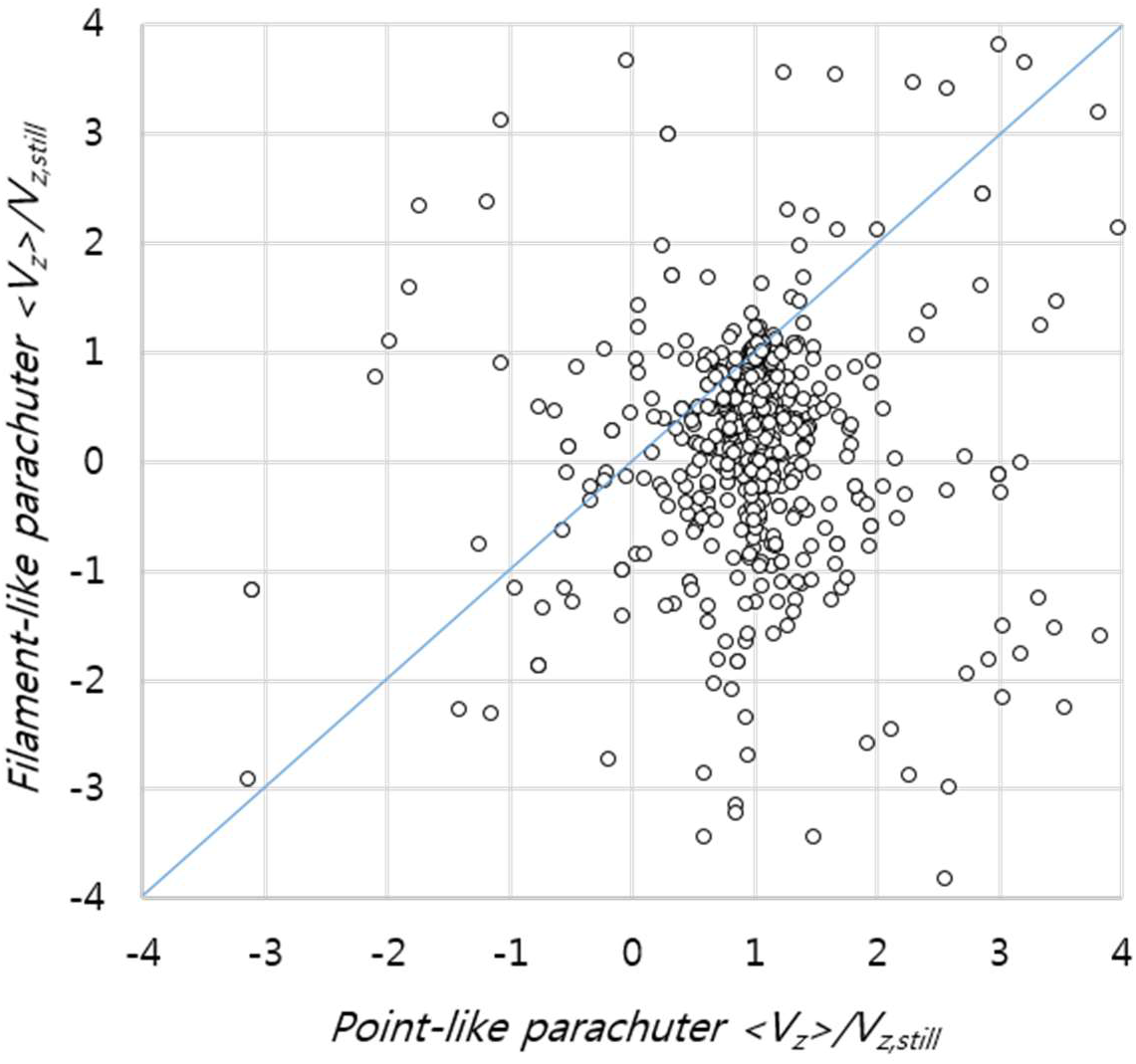
Simultaneous plot of the dimensionless settling speeds for a filament-like parachuter and a point-like parachuter. Both cases are simultaneously simulated in an identical turbulence model.

## 4 Discussion

### 4.1.1 Role of Anisotropic Drag of a Filament

The anisotropic drag of a thin filament structure is a crucial aspect of the dynamics of a microswimmer. The cilia and flagella of a microswimmer enable organisms and cells to move forward in a viscous medium (Purcell 1977, Brennen and Winet 1977, Fung 1990, Happel and Brenner 1991, Lauga and Powers 2009). In the low Reynolds number flow, the perpendicular drag on a filament is about two times larger than its tangential drag. Because of this drag anisotropy, a flagellum can slip in a certain direction in a viscous fluid and can also regulate the fluid-dynamic force on it by deforming its shape for the locomotion of a microswimmer. This character of the anisotropic drag is valid not only in the active dynamics of a swimming microorganism, but also in the passive dynamics of an artificial microswimmer, such as a flexible magnetic filament in an oscillating external magnetic field (Dreyfus et al. 2005, Gauger and Stark 2006). From this, we may infer that the anisotropic drag is also an important factor in the passive dynamics of a flexible filament moved by an external fluid flow.

In the simulation, the influence of the anisotropic drag is checked by releasing the ballooning structure in various shear flow strengths. In the bead-spring model, the drag anisotropy is described by the Rotne-Prager hydrodynamic interaction between beads (Rotne and Prager 1969, Dhont 2003, Gauger and Stark 2006). The drag anisotropy modelled by the hydrodynamic interaction was shown in S.M. Figure S1 and compared with the experimental data. The velocity reduction of the ballooning structure in a shear flow shows that the reduction is due to the drag anisotropy; this is because the model without hydrodynamic interaction did not exhibit any velocity reduction effect during sedimentation in a shear flow (see Figure 8B).

The main cause of reduction of the settling speed in a shear flow is the deformed geometrical shape of a filament and the resulting anisotropic drag. However, the geometric contribution reaches a maximum at the ballooning number of 10. After that the reduction of the settling speed is caused by the shear-induced resistance coefficient (see Figure 9).

### 4.1.2 Trapping Phenomenon by a Vortical Flow

The simulation in a cellular vortical flow field reveals various motions according to the flow conditions (see Figure 12). In the simulation, two flow condition parameters are considered: the size *L* of a vortex and the strength (maximum flow velocity *U*_*0*_) of the vortex. In a previous study of particle sedimentation, Stout et al. (1995) categorised three regimes of heavy particle motion in turbulent flow (see Figure 16): (i) suspension for σ/*W*_*T*_ ≫ 1 (Stommel 1949, Manton 1974); (ii) preferential sweeping for σ/*W*_*T*_ ≈ 1 (Maxey and Corrsin 1986, Maxey 1987b, Wang and Maxey 1993, Fung 1997, 1998, Toschi and Bodenschatz 2009, Balachandar and Eaton 2010); (iii) eddy crossing for σ/*W*_*T*_ ≈ 1 (Stout et al. 1995; see Figure 16). The ballooning structure in a cellular flow field exhibited similar behaviours to particle motions in the turbulent flow (Maxey and Corrsin 1986, Maxey 1987a, Stout et al. 1995, Fung 1997, Bergougnoux et al. 2014). The eddy crossing motion of the ballooning structure in Figure 11(a)(d)(g) is similar to the eddy crossing motion of the heavy particle (see Figure 16(a)). If the point mass of the ballooning structure is heavy enough compared with the resistance force applied to a filament by the eddies, the structure crosses eddies (vortices) in a cellular flow field, and there is no retardation of the settling speed. However, if the mass is comparable to the resistance force on a filament, then the fast settling motion of the ballooning structure in Figure 11(e)(h)(i) is very similar to the preferential sweeping motion in Stout’s theory of heavy particle motion (see Figure 16(b)). The ballooning structure is preferentially swept into the regions of the downdraft path (Maxey 1987b, Stout et al. 1995). This preferential sweeping of the heavy particle is caused by its inertia. However, as the present simulation does not consider the inertia of the point mass, the preferential sweeping of the ballooning structure may thus be caused by the anisotropic drag of a filament structure. If the vortex velocity is dominant and the size of a vortex is large, the ballooning structure is trapped by the vortex (see Figure 11(c)(f)). This is very interesting because the heavy particle does not settle down with the help of a filament. Although the ballooning structure produces negative buoyancy (toward a ground) because of its weight, the ballooning structure either stays in the upward current region of a vortex (see Figure 12B) or gains energy by cycling with a passive motion (see Figure 12A). This trapping motion was also observed by Zhao et al. (2017), who simulated the structure in a cavity flow.

**Figure 16.**
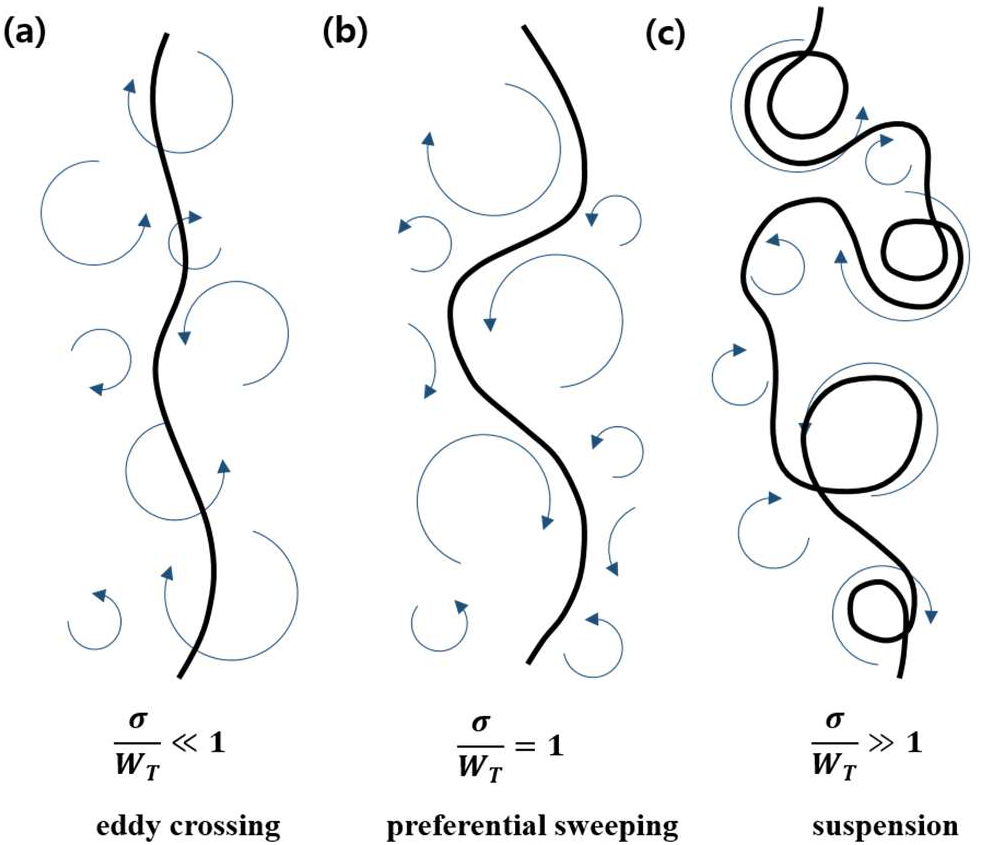
Three different regimes of heavy particle motion (redrawn from Stout el al. 1995)

Abundant turbulent eddies exist in the atmospheric boundary layer. These are mostly produced (i) by mechanical terrains on the earth’s surface, (ii) by rising thermal air parcels, and (iii) by the shear-winds on the surface. These eddies mostly develop upwards as they draft in the downwind direction because of the blockage of the ground (Adrian et al. 2000, Hunt and Morrison 2000, Hommema and Adrian 2003, Adrian 2007, Finnigan et. al 2009). These upward developing eddies, which have certain sizes and strengths, may be helpful to the ballooning spiders by trapping them and allowing them to stay in the air for longer periods.

### 4.1.3 Biased Behaviour in Homogeneous Turbulence

As previously discussed (see Sections 4.1.1 and 4.1.2), a thin filament structure (spider silk) shows a nonlinear drag character on a point-mass (spider) in a non-uniform flow. In a simple shear flow, the ballooning structure settles more slowly than it would in a still medium. In a cellular flow field, the same structure settles faster in small-size and relatively strong vortex cells, while either settling more slowly or becoming trapped in large-size and relatively strong vortex cells. This nonlinear character in homogeneous turbulence also results in a biased settling behaviour (biased toward slow settling) of the structure (see Figure 6B). Reynolds et al. (2006) first proposed that turbulent wind may reduce the rate of fall of the spider by deforming and stretching the flexible fibres. Their study points out that the extensibility and flexibility of a filament may be the causes of the settling retardation. However, as previously discussed in Sections 4.1.1 and 4.1.2, the cause of the settling retardation seems to be the anisotropic drag character of a thin filament and flexible deformation in vortex eddies in turbulence. As the model (extendible and flexible filament) proposed by Reynolds et al. did not consider the hydrodynamic interaction between beads (Rouse model; Rouse 1953, Dhont 2003, Reynolds et al. 2006, Doi and Edwards 2007), the cause of the retardation in Reynolds’ simulation seems to be flexibility (not extensibility), which helps the filament to become deformed and trapped in a quasi-vortical flow in the turbulence model.

Ballooning silks have a longer length-scale (i.e., a few metres) compared to an aerial seed (a few centimetres) (Sudo et al. 2008, Cho et al. 2018). Therefore, each segment of the ballooning silk is exposed to different wind flow fields in direction and magnitude. By contrast, an aerial seed is exposed to quasi-uniform airflow, although it still drifts in the fluctuating winds (see Figure 13). Accordingly, the question is whether the exertion of these non-uniform flows on the ballooning silk generates extra buoyancy capability during a ballooning flight. As shown in Figure 14, the filament-like parachuter evidently shows biased behaviour toward either slow settling (in comparison with that of the point-like parachuter) or rising (see also Figure 15). As discussed in Sections 4.1.1 and 4.1.2, this biased behaviour seems to be induced by both the anisotropic drag of a filament and the vortical flows in homogeneous turbulence.

### 4.1.4 Simulation and Ballooning Flight in Nature

Spiders’ ballooning can be understood as a physical phenomenon in which two different physical scales are connected: the macroscopic weight of the spider, although insects of such size normally utilise a high Reynolds number flow for their flight, is sustained by the micro-fluid-dynamic forces that are harvested through the use of long thin fibres. The physical properties of these two different scales are non-dimensionalised and expressed as the ballooning number (see Equations (2.15)-(2.18)). An increase in the ballooning number means that the fluid-dynamic forces exerted on a silk by the non-uniform flow become dominant compared with the force of gravity that acts on the body. Moreover, a decrease in the ballooning number means that the gravity force on the body becomes more dominant than the fluid-dynamic forces on a silk, meaning that the spider will fall like a heavy particle, cutting through the eddies of the air flow.

If we assume that a spider flies in a 1.96 m s^−1^ mean wind speed (measured data 0.95 m above the ground), the root-mean-square fluctuation velocity is 0.41 m s^−1^ on a sunny day with a light breeze. Thus, the ballooning number of a 20-mg crab spider for turbulence in the field may be about 7.42 (see Equation (4.1); dynamic viscosity of air: 1.837 × 10^−5^ kg m^−1^ s^−1^; length of a spider silk: 3.22 m; number of silks: 60). This number belongs to the order in which the behaviour of the ballooning structure is biased (towards slow settling in the turbulence; see Figure 6B). The observed shapes of the ballooning silks in the field are also believed to be events in the range of the ballooning number 1 < *Bn*_*tUr*_ < 10 (Cho et al. 2018). The superposed shapes of these ballooning silks in the field are compared with the results of the bead-spring model simulation (see Figure 17). As Figure 17 shows the geometric analogy, the introduction of the ballooning number in the spiders’ flight seems to be reasonable.

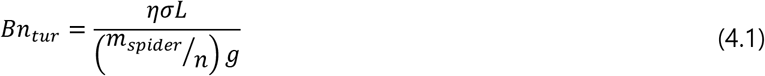

**Figure 17.**
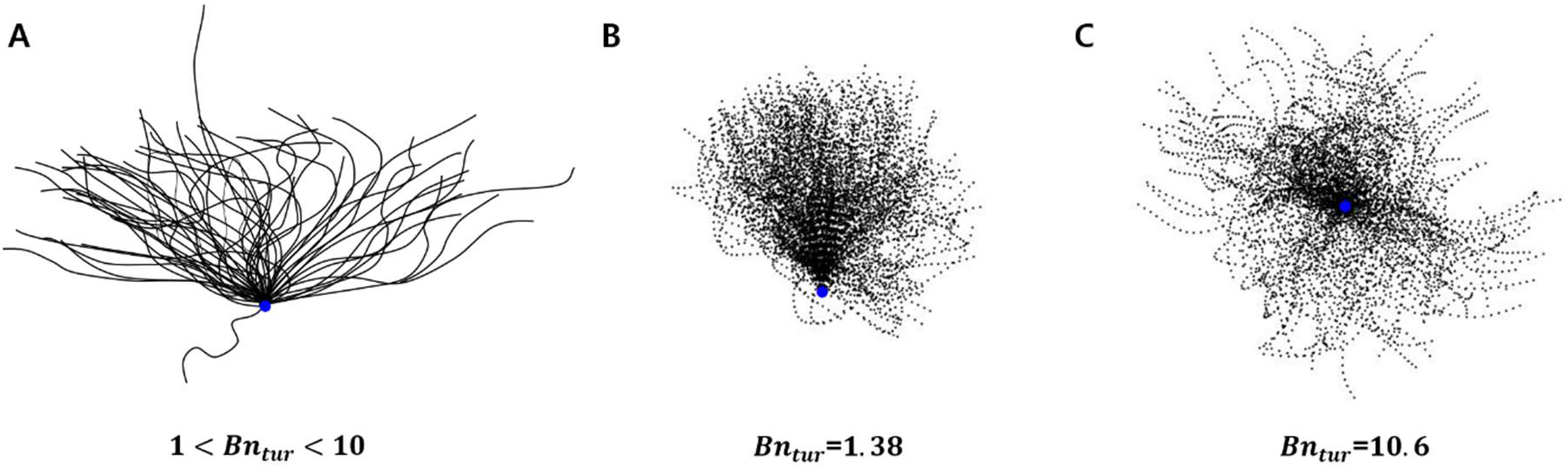
Superposed shapes of ballooning silks during the flight. The reference point is the position of its body. (A) Superposed shapes of the observed ballooning events in the nature. (1 < *Bn*_*tUr*_ < 10; Cho et al. 2018) (B) Superposed shapes of the ballooning silks in the simulation (*Bn*_*tUr*_ = 1.38). The blue point means the position of a spider body (a point-mass). (C) Superposed shapes of the ballooning silks in the simulation (*Bn*_*tUr*_ = 10.6). The blue point means the position of a spider body (a point-mass).

## 5 Conclusion

While the dynamics of a particle and a filament in a fluid flow have been independently and widely studied, the combined asymmetric structure of the two has been investigated far less extensively. This ‘combined asymmetric structure’ refers to a filament at the end of which a point-mass is attached. This structure is also a physical model of a ballooning spider in the air. Here, the suspension dynamics of the structure were studied using numerical simulation. For the generalisation of the parameters, the ballooning number is newly defined as the ratio of the Stokes force on a filament to the gravity force on the weight. The simulation results revealed interesting characteristics of the ballooning structure in non-uniform flows. First, the structure in a shear flow settles more slowly than would be the case in a still medium. This slow settling is caused by the anisotropic drag of a filament in a low Reynolds number flow. The slow settling occurs primarily due to the geometrically inclined attitude of a filament; secondarily, it occurs due to the force induced by and from a shear flow. Moreover, in a cellular flow field, the structure showed various settling and suspension modes. These motions mostly depend on the size and the strength of a cellular flow, and are thereby categorised according to these parameters. At the low ballooning number *Bn*_*vor*_ < 1, the structure behaved like a heavy particle; there is neither a retardation effect nor an acceleration effect 〈*V*_*z*_〉/*V*_*z,still*_ ≈ 1. At the high ballooning number *Bn*_*vor*_ > 1 and small vortex cell size *L*_*c*_/*L*_*s*_ < 1.2, the structure is swept up by the vortical flows and settles faster than would be the case in a still medium. Furthermore, if the size of a vortex cell is larger than 1.2 *L*_*s*_ and *Bn*_*vor*_ is larger than 1, the ballooning structure settles slowly or is trapped by a vortex and suspends with 〈*V*_*z*_〉/*V*_*z,still*_ ≈ 0. Different initial conditions of *U*_*0*_ were also applied. Their motions converged to trapping motion by vortex cells at the end. While the point-mass (spider’s body) alone cannot be trapped by a vortex, this heavy particle (spider) is trapped with the help of the filament (spider silk). The ballooning structure also shows biased behaviour toward slow settling under homogeneous turbulence, while that of a point-like aerial seed was not similarly biased under identical turbulence conditions. As the nonlinearity of anisotropic drag of a flexible filament, which is deformed by non-uniform flows, was recognised as the cause of retardation and trapping of the ballooning structure in a shear flow and a cellular flow field, the drag anisotropy of a spider silk and the vortex trapping motion seem to be the main reasons they can suspend in the convective air current for a long time. Due to the abundance of turbulent eddies in the atmosphere, these retarding and trapping phenomena are plausible in spiders’ ballooning flight and can explain how they acquire energies from the non-uniform air current for their long aerial journey.

## 6 Acknowledgments

The author deeply thanks Prof. Dr. Ing. Klaus Affeld for scientific discussions and support concerning this project.

## Notes

### Competing Interest Statement

The authors have declared no competing interest.

### Summary of Updates

A few parts in modeling are moved into the supplementary material.

## References

Adrian, R. J. (2007): Hairpin vortex organization in wall turbulence. In Physics of Fluids 19 (4), p. 41301. DOI: 10.1063/1.2717527.

Adrian, R. J.; Meinhart, C. D.; Tomkins, C. D. (2000): Vortex organization in the outer region of the turbulent boundary layer. In J. Fluid Mech. 422, pp. 1–54. DOI: 10.1017/S0022112000001580.

Balachandar, S.; Eaton, John K. (2010): Turbulent Dispersed Multiphase Flow. In Annu. Rev. Fluid Mech. 42 (1), pp. 111–133. DOI: 10.1146/annurev.fluid.010908.165243.

Bearman, P. W. (1972): Some measurements of the distortion of turbulence approaching a two-dimensional bluff body. In J. Fluid Mech. 53 (03), p. 451. DOI: 10.1017/S0022112072000254.

Bergougnoux, Laurence; Bouchet, Gilles; Lopez, Diego; Guazzelli, Élisabeth (2014): The motion of solid spherical particles falling in a cellular flow field at low Stokes number. In Physics of Fluids 26 (9), p. 93302. DOI: 10.1063/1.4895736.

Bonino, Mark J. (2003). Material Properties of Spider Silk. Degree of Master of Science. University of Rochester, Rochester, NY.

Brennen, C.; Winet, H. (1977): Fluid Mechanics of Propulsion by Cilia and Flagella. In Annu. Rev. Fluid Mech. 9 (1), pp. 339–398. DOI: 10.1146/annurev.fl.09.010177.002011.

Brouzet, C.; Verhille, G.; Le Gal, P. (2014): Flexible fiber in a turbulent flow: a macroscopic polymer. In Physical review letters 112 (7), p. 74501. DOI: 10.1103/PhysRevLett.112.074501.

Bustamante, C.; Marko, J.; Siggia, E.; Smith, S. (1994): Entropic elasticity of lambda-phage DNA. In Science 265 (5178), pp. 1599–1600. DOI: 10.1126/science.8079175.

Bustamante, C.; Smith, S. B.; Liphardt, J.; Smith, D. (2000): Single-molecule studies of DNA mechanics. In Current opinion in structural biology 10 (3), pp. 279–285.

Childress, Stephen (1981): Mechanics of swimming and flying. Cambridge: Cambridge University Press (Cambridge studies in mathematical biology, 2). Available online at https://doi.org/10.1017/CBO9780511569593.

Cho, Moonsung; Neubauer, Peter; Fahrenson, Christoph; Rechenberg, Ingo (2018): An observational study of ballooning in large spiders: Nanoscale multifibers enable large spiders’ soaring flight. In PLoS biology 16 (6), e2004405. DOI: 10.1371/journal.pbio.2004405.

Cho, Moonsung (2020): Suspension of a Point-Mass-Loaded Filament in Non-Uniform Flows: The Ballooning Flight of Spiders. PhD Thesis. Technical University of Berlin Repository

Dai, Liang; van der Maarel, Johan; Doyle, Patrick S. (2014): Extended de Gennes Regime of DNA Confined in a Nanochannel. In Macromolecules 47 (7), pp. 2445–2450. DOI: 10.1021/ma500326w.

Darwin, Charles (1845): Journal of researches into the natural history and geology of the countries visited during the voyage of H.M.S. Beagle round the world,, under the Command of Capt. Fitz Roy, R.N. Second Edition. New York: John Murray, Albemarle Street.

Das, Dibyendu; Sabhapandit, Sanjib (2008): Accurate statistics of a flexible polymer chain in shear flow. In Physical review letters 101 (18), p. 188301. DOI: 10.1103/PhysRevLett.101.188301.

Delgado-Buscalioni, Rafael (2006): Cyclic motion of a grafted polymer under shear flow. In Physical review letters 96 (8), p. 88303. DOI: 10.1103/PhysRevLett.96.088303.

Delmotte, Blaise; Climent, Eric; Plouraboué, Franck (2015): A general formulation of Bead Models applied to flexible fibers and active filaments at low Reynolds number. In Journal of Computational Physics 286, pp. 14–37. DOI: 10.1016/j.jcp.2015.01.026.

Dhont, Jan K. G. (2003): An introduction to dynamics of colloids. 2. impr. Amsterdam: Elsevier (Studies in interface science, 2).

Doi, Masao; Edwards, Samuel F. (2007): The theory of polymer dynamics. Oxford: Clarendon Press (International series of monographs on physics, 73).

Doyle, P. S.; Ladoux, B.; Viovy, J. L. (2000): Dynamics of a tethered polymer in shear flow. In Physical review letters 84 (20), pp. 4769–4772. DOI: 10.1103/PhysRevLett.84.4769.

Dreyfus, Rémi; Baudry, Jean; Roper, Marcus L.; Fermigier, Marc; Stone, Howard A.; Bibette, Jérôme (2005): Microscopic artificial swimmers. In Nature 437 (7060), pp. 862–865. DOI: 10.1038/nature04090.

Drummond, I. T.; Duane, S.; Horgan, R. R. (1984): Scalar diffusion in simulated helical turbulence with molecular diffusivity. In J. Fluid Mech. 138 (−1), p. 75. DOI: 10.1017/S0022112084000045.

Finnigan, J. J.; Shaw, Roger H.; Patton, Edward G. (2009): Turbulence structure above a vegetation canopy. In J. Fluid Mech. 637, p. 387. DOI: 10.1017/S0022112009990589.

Fung, J.; Vassilicos, J. (1998): Two-particle dispersion in turbulent like flows. In Phys. Rev. E 57 (2), pp. 1677–1690. DOI: 10.1103/PhysRevE.57.1677.

Fung, J. C. H. (1993): Gravitational settling of particles and bubbles in homogeneous turbulence. In J. Geophys. Res. 98 (C11), p. 20287. DOI: 10.1029/93JC01845.

Fung, J. C. H. (1998): Effect of nonlinear drag on the settling velocity of particles in homogeneous isotropic turbulence. In J. Geophys. Res. 103 (C12), pp. 27905–27917. DOI: 10.1029/98JC02822.

Fung, J. C. H.; Hunt, J. C. R.; Malik, N. A.; Perkins, R. J. (1992): Kinematic simulation of homogeneous turbulence by unsteady random Fourier modes. In J. Fluid Mech. 236 (−1), p. 281. DOI: 10.1017/S0022112092001423.

Fung, J. C. H.; Perkins, R. J. (2008): Dispersion modeling by kinematic simulation: Cloud dispersion model. In Fluid Dyn. Res. 40 (4), pp. 273–309. DOI: 10.1016/j.fluiddyn.2007.06.005.

Fung, J. C. H.; Vassilicos, J. C. (2003): Inertial particle segregation by turbulence. In Physical review. E, Statistical, nonlinear, and soft matter physics 68 (4 Pt 2), p. 46309. DOI: 10.1103/PhysRevE.68.046309.

Fung, J.C.H. (1997): Gravitational settling of small spherical particles in unsteady cellular flow fields. In Journal of Aerosol Science 28 (5), pp. 753–787. DOI: 10.1016/S0021-8502(96)00478-8.

Fung, Jimmy Chi Hung (1990): Kinematic simulation of turbulent flow and particle motions. Apollo - University of Cambridge Repository.

Fung, Y. C. (1990): Biomechanics. Motion, Flow, Stress, and Growth. New York, NY: Springer. Available online at http://dx.doi.org/10.1007/978-1-4419-6856-2.

Gauger, Erik; Stark, Holger (2006): Numerical study of a microscopic artificial swimmer. In Physical review. E, Statistical, nonlinear, and soft matter physics 74 (2 Pt 1), p. 21907. DOI: 10.1103/PhysRevE.74.021907.

Gerashchenko, Sergiy; Steinberg, Victor (2006): Statistics of tumbling of a single polymer molecule in shear flow. In Physical review letters 96 (3), p. 38304. DOI: 10.1103/PhysRevLett.96.038304.

Glick, P. A. (1939): The distribution of insects, spiders, and mites in the air. In USDA Tech Bull (673), pp. 1–150.

Gorham, Peter W. (2013): Ballooning Spiders: The Case for Electrostatic Flight. Available online at https://arxiv.org/abs/1309.4731v2.

Gosline, J. M.; Guerette, P. A.; Ortlepp, C. S.; Savage, K. N. (1999): The mechanical design of spider silks: from fibroin sequence to mechanical function. In The Journal of experimental biology 202 (Pt 23), pp. 3295–3303.

Gustavsson, K.; Meneguz, E.; Reeks, M.; Mehlig, B. (2012): Inertial-particle dynamics in turbulent flows: caustics, concentration fluctuations and random uncorrelated motion. In New J. Phys. 14 (11), p. 115017. DOI: 10.1088/1367-2630/14/11/115017.

Happel, John; Brenner, Howard (1991): Low Reynolds number hydrodynamics. With special applications to particulate media. 5. printing. Dordrecht: Kluwer Acad. Publ (Mechanics of fluids and transport processes).

Harris, I. (Ed.) (1971): The nature of the wind. Proc. Seminar at Inst. Civil Engrs, June 1970.

Hommema, Scott E.; Adrian, Ronald J. (2003): Packet structure of surface eddies in the atmospheric boundary layer. In Boundary-Layer Meteorology 106 (1), pp. 147–170. DOI: 10.1023/A:1020868132429.

Humphrey, J. A. C. (1987): Fluid mechanic constraints on spider ballooning. In Oecologia 73 (3), pp. 469–477. DOI: 10.1007/BF00385267.

Hunt, J. C. R. (1973): A theory of turbulent flow round two-dimensional bluff bodies. In J. Fluid Mech. 61 (04), p. 625. DOI: 10.1017/S0022112073000893.

Hunt, Julian C.R.; Morrison, Jonathan F. (2000): Eddy structure in turbulent boundary layers. In European Journal of Mechanics - B/Fluids 19 (5), pp. 673–694. DOI: 10.1016/S0997-7546(00)00129-1.

Ibáñez-García, Gabriel O.; Hanna, Simon (2009): Relaxation of an initially-stretched, tethered polymer under shear flow: a Brownian dynamics simulation. In Soft Matter 5 (22), p. 4464. DOI: 10.1039/b916087f.

Jian, Hongmei; Vologodskii, Alexander V.; Schlick, Tamar (1997): A Combined Wormlike-Chain and Bead Model for Dynamic Simulations of Long Linear DNA. In Journal of Computational Physics 136 (1), pp. 168–179. DOI: 10.1006/jcph.1997.5765.

Jo, Kyubong; Chen, Yeng-Long; Pablo, Juan J. de; Schwartz, David C. (2009): Elongation and migration of single DNA molecules in microchannels using oscillatory shear flows. In Lab on a chip 9 (16), pp. 2348–2355. DOI: 10.1039/b902292a.

Joel, Anna-Christin; Baumgartner, Werner (2017): Nanofibre production in spiders without electric charge. In The Journal of experimental biology 220 (Pt 12), pp. 2243–2249. DOI: 10.1242/jeb.157594.

Kraichnan, Robert H. (1970): Diffusion by a Random Velocity Field. In Phys. Fluids 13 (1), p. 22. DOI: 10.1063/1.1692799.

Kronenberger, Katrin; Vollrath, Fritz (2015): Spiders spinning electrically charged nano-fibres. In Biology letters 11 (1), p. 20140813. DOI: 10.1098/rsbl.2014.0813.

Ladoux, B.; Doyle, P. S. (2000): Stretching tethered DNA chains in shear flow. In Europhys. Lett. 52 (5), pp. 511–517. DOI: 10.1209/epl/i2000-00467-y.

Lafitte, Anthony; Le Garrec, Thomas; Bailly, Christophe; Laurendeau, Estelle (2014): Turbulence Generation from a Sweeping-Based Stochastic Model. In AIAA Journal 52 (2), pp. 281–292. DOI: 10.2514/1.J052368.

Larson, R. G.; Perkins, T. T.; Smith, D. E.; Chu, S. (1997): Hydrodynamics of a DNA molecule in a flow field. In Phys. Rev. E 55 (2), pp. 1794–1797. DOI: 10.1103/PhysRevE.55.1794.

Lauga, Eric; Powers, Thomas R. (2009): The hydrodynamics of swimming microorganisms. In Rep. Prog. Phys. 72 (9), p. 96601. DOI: 10.1088/0034-4885/72/9/096601.

Lee, Nam-Kyung; d. Thirumalai (2004): Pulling-Speed-Dependent Force-Extension Profiles for Semiflexible Chains. In Biophysical Journal 86 (5), pp. 2641–2649. DOI: 10.1016/S0006-3495(04)74320-9.

Lindström, Stefan B.; Uesaka, Tetsu (2007): Simulation of the motion of flexible fibers in viscous fluid flow. In Physics of Fluids 19 (11), p. 113307. DOI: 10.1063/1.2778937.

Litvinov, S.; Hu, X. Y.; Adams, N. A. (2011): Numerical simulation of tethered DNA in shear flow. In Journal of physics. Condensed matter : an Institute of Physics journal 23 (18), p. 184118. DOI: 10.1088/0953-8984/23/18/184118.

Manton, M. J. (1974): On the motion of a small particle in the atmosphere. In Boundary-Layer Meteorol 6 (3-4), pp. 487–504. DOI: 10.1007/BF02137681.

Maxey, M. R. (1987a): The gravitational settling of aerosol particles in homogeneous turbulence and random flow fields. In J. Fluid Mech. 174 (1), p. 441. DOI: 10.1017/S0022112087000193.

Maxey, M. R. (1987b): The motion of small spherical particles in a cellular flow field. In Phys. Fluids 30 (7), p. 1915. DOI: 10.1063/1.866206.

Maxey, M. R.; Corrsin, S. (1986): Gravitational Settling of Aerosol Particles in Randomly Oriented Cellular Flow Fields. In J. Atmos. Sci. 43 (11), pp. 1112–1134. DOI: 10.1175/1520-0469(1986)0431112:GSOAPitalic2.0.CO;2.

Morley, Erica L.; Robert, Daniel (2018): Electric Fields Elicit Ballooning in Spiders. In Current biology : CB 28 (14), 2324–2330.e2. DOI: 10.1016/j.cub.2018.05.057.

Ning, Zemin; Melrose, John R. (1999): A numerical model for simulating mechanical behavior of flexible fibers. In The Journal of Chemical Physics 111 (23), pp. 10717–10726. DOI: 10.1063/1.480426.

Ortega-Jimenez, Victor Manuel; Dudley, Robert (2013): Spiderweb deformation induced by electrostatically charged insects. In Scientific reports 3, p. 2108. DOI: 10.1038/srep02108.

Perkins, T.; Quake; Smith, D.; Chu, S. (1994): Relaxation of a single DNA molecule observed by optical microscopy. In Science 264 (5160), pp. 822–826. DOI: 10.1126/science.8171336.

Perkins, T.; Smith, D.; Larson, R.; Chu, S. (1995): Stretching of a single tethered polymer in a uniform flow. In Science 268 (5207), pp. 83–87. DOI: 10.1126/science.7701345.

Purcell, E. M. (1977): Life at low Reynolds number. In American Journal of Physics 45 (1), pp. 3–11. DOI: 10.1119/1.10903.

Reynolds, A. M.; Bohan, D. A.; Bell, J. R. (2006): Ballooning dispersal in arthropod taxa with convergent behaviours: dynamic properties of ballooning silk in turbulent flows. In Biology letters 2 (3), pp. 371–373. DOI: 10.1098/rsbl.2006.0486.

Rotne, Jens; Prager, Stephen (1969): Variational Treatment of Hydrodynamic Interaction in Polymers. In The Journal of Chemical Physics 50 (11), pp. 4831–4837. DOI: 10.1063/1.1670977.

Rouse, Prince E. (1953): A Theory of the Linear Viscoelastic Properties of Dilute Solutions of Coiling Polymers. In The Journal of Chemical Physics 21 (7), pp. 1272–1280. DOI: 10.1063/1.1699180.

Schroeder, Charles M.; Shaqfeh, Eric S. G.; Chu, Steven (2004): Effect of Hydrodynamic Interactions on DNA Dynamics in Extensional Flow: Simulation and Single Molecule Experiment. In Macromolecules 37 (24), pp. 9242–9256. DOI: 10.1021/ma049461l.

Schroeder, Charles M.; Teixeira, Rodrigo E.; Shaqfeh, Eric S. G.; Chu, Steven (2005): Characteristic periodic motion of polymers in shear flow. In Physical review letters 95 (1), p. 18301. DOI: 10.1103/PhysRevLett.95.018301.

Shaqfeh, Eric S.G. (2005): The dynamics of single-molecule DNA in flow. In Journal of Non-Newtonian Fluid Mechanics 130 (1), pp. 1–28. DOI: 10.1016/j.jnnfm.2005.05.011.

Stommel, H. (1949): Trajectories of small bodies sinking slowly through convection cells. In J. Mar. Res. 8, pp. 24–29.

Stout, J. E.; Arya, S. P.; Genikhovich, E. L. (1995): The Effect of Nonlinear Drag on the Motion and Settling Velocity of Heavy Particles. In J. Atmos. Sci. 52 (22), pp. 3836–3848. DOI: 10.1175/1520-0469(1995)052<3836:TEONDO>2.0.CO;2.

Sudo, S., Matsui, N., Tsuyuki, K., & Yano, T. (2008). Morphological Design of Dandelion. Proceedings of the XIth International Congress and Exposition, June 2-5, 2008 Orlando, Florida USA, Society for Experimental Mechanics.

Suter, Robert B. (1991): Ballooning in spiders. Results of wind tunnel experiments. In Ethology Ecology & Evolution 3 (1), pp. 13–25. DOI: 10.1080/08927014.1991.9525385.

Suter, Robert B. (1992): Ballooning: Data from Spiders in Freefall Indicate the Importance of Posture. In The Journal of Arachnology 2 (20), pp. 107–113. Available online at http://www.jstor.org/stable/3705774.

Suter, Robert B. (1999): An Aerial Lottery: The Physics of Ballooning in a Chaotic Atmosphere 1 (27), pp. 281–293. Available online at http://www.jstor.org/stable/3705999.

Suzuki, Takeshi; Sakai, Yasuhiko (2013): Numerical Simulation of Reactive Turbulent Scalar Mixing Layer by the Random Fourier Modes Method and Lagrangian Molecular Mixing Model. In Transactions of the Japan Society of Mechanical Engeneers, Series B 79 (798), pp. 104–114. DOI: 10.1299/kikaib.79.104.

Toschi, Federico; Bodenschatz, Eberhard (2009): Lagrangian Properties of Particles in Turbulence. In Annu. Rev. Fluid Mech. 41 (1), pp. 375–404. DOI: 10.1146/annurev.fluid.010908.165210.

Turfus C. (1985): Stochastic Modelling of Turbulent Dispersion Near Surfaces: University of Cambridge.

Vassilicos, J. C.; Fung, J. C. H. (1995): The self-similar topology of passive interfaces advected by two-dimensional turbulent-like flows. In Physics of Fluids 7 (8), pp. 1970–1998. DOI: 10.1063/1.868510.

Verhille, Gautier; Bartoli, Adrien (2016): 3D conformation of a flexible fiber in a turbulent flow. In Exp Fluids 57 (7), p. 74501. DOI: 10.1007/s00348-016-2201-1.

Vollrath, Fritz; Edmonds, Donald (2013): Consequences of electrical conductivity in an orb spider’s capture web. In Die Naturwissenschaften 100 (12), pp. 1163–1169. DOI: 10.1007/s00114-013-1120-8.

Wang, Jun; Lu, Chang (2007): Single molecule λ-DNA stretching studied by microfluidics and single particle tracking. In Journal of Applied Physics 102 (7), p. 74703. DOI: 10.1063/1.2786896.

Wang, Lian-Ping; Maxey, Martin R. (1993): Settling velocity and concentration distribution of heavy particles in homogeneous isotropic turbulence. In J. Fluid Mech. 256 (-1), p. 27. DOI: 10.1017/S0022112093002708.

Winkler, Roland G. (2006): Semiflexible polymers in shear flow. In Physical review letters 97 (12), p. 128301. DOI: 10.1103/PhysRevLett.97.128301.

Wong, P. Kina. K.; Lee, Yi-Kuen; Ho, Chih-Ming (2003): Deformation of DNA molecules by hydrodynamic focusing. In J. Fluid Mech. 497, pp. 55–65. DOI: 10.1017/S002211200300658X.

Yamamoto, Satoru; Matsuoka, Takaaki (1993): A method for dynamic simulation of rigid and flexible fibers in a flow field. In The Journal of Chemical Physics 98 (1), pp. 644–650. DOI: 10.1063/1.464607.

Zhao, Longhua; Panayotova, Lordanka N.; Chuang, Angela; Sheldon, Kimberly S.; Bourouiba, Lydia; Miller, Laura A. (2017): Flying Spiders: Simulating and Modeling the Dynamics of Ballooning. In Anita T. Layton, Laura A. Miller (Eds.): Women in Mathematical Biology, vol. 8. Cham: Springer International Publishing (Association for Women in Mathematics Series), pp. 179–210.

