## Supplementary material for "Suspension of a Point-Mass-Loaded Filament in Non-Uniform Flows: Passive Dynamics of a Ballooning Spider": S.M.

### S.1 Modelling

#### S.1.1 Bead Spring Model

Each bead in the bead spring model experiences hydrodynamic viscous drag in a low Reynolds number flow, which is governed by Stokes' law (see Equation (S.1); Happel and Brenner 1986). $\eta$ indicates dynamic viscosity of fluid medium. $a$ is the radius of a bead and $U$ is the relative flow velocity on the bead.

$$F = 6\pi\eta a U \tag{S.1}$$

In a low Reynolds number flow, the influence of inertia is almost negligible. This means that at every moment, the net force on the structure is zero (see Equation (S.2) ; Doyle and Underhill 2005). The net force $\boldsymbol{F}_i^{total}$ on each bead includes a fluid-dynamic force $\boldsymbol{F}_i^F$, a gravity force $\boldsymbol{F}_i^G$, a stretching force $\boldsymbol{F}_i^S$ and a bending force $\boldsymbol{F}_i^B$:

$$\boldsymbol{F}_i^{total} = \boldsymbol{F}_i^F + \boldsymbol{F}_i^G + \boldsymbol{F}_i^S + \boldsymbol{F}_i^B \approx 0 \tag{S.2}$$

The fluid-dynamic force is proportional to the fluid velocity. Therefore, resistances are defined as a reciprocal of bead mobilities $\boldsymbol{\mu}_{ij}$:

$$\boldsymbol{F}_j^F = \frac{1}{\boldsymbol{\mu}_{ij}}\left(\boldsymbol{u}^{\infty}(\boldsymbol{r}_i) - \frac{d\boldsymbol{r}_i}{dt}\right) \tag{S.3}$$

The bead mobility is derived through the consideration of the hydrodynamic interactions between beads. Here, the Rotne-Prager approximation, applicable to different sizes of beads, is used (see Equation (S.5)). Equation (S.6) shows the self-induction elements of the mobility tensor $\boldsymbol{\mu}_{ij}$ (Jeffrey and Onishi 1984, Gauger and Stark 2006).

$$\frac{d\boldsymbol{r}_i}{dt} = \boldsymbol{u}^\infty(\boldsymbol{r}_i) + \sum_j \boldsymbol{\mu}_{ij}\left(\boldsymbol{F}_j^G + \boldsymbol{F}_j^S + \boldsymbol{F}_j^B\right) \quad \text{(S.4)}$$

$$\boldsymbol{\mu}_{ij} = \frac{1}{6\pi\eta\boldsymbol{r}_{ij}}\left[\frac{3}{4}\left(\boldsymbol{1} + \hat{\boldsymbol{r}}_{ij} \otimes \hat{\boldsymbol{r}}_{ij}\right) + \frac{1}{4}\frac{a_i^2 + a_j^2}{\boldsymbol{r}_{ij}^2}\left(\boldsymbol{1} - 3\hat{\boldsymbol{r}}_{ij} \otimes \hat{\boldsymbol{r}}_{ij}\right)\right] \quad \text{(S.5)}$$

$$\boldsymbol{\mu}_{ii} = \mu_0\boldsymbol{1} \quad \text{with} \quad \mu_0 = \frac{1}{6\pi\eta a_i} \quad \text{(S.6)}$$

Equation (S.7)) describes the gravity force on each bead.

$$\boldsymbol{F}_i^G = [0 \quad 0 \quad -v_i(\rho_i - \rho_{air})g]^{\mathrm{T}} \quad \text{(S.7)}$$

Here, we consider a linearly varying spring. Therefore, the total stretching energy can be described as in Equation (S.8), while the stretching force, Equation (S.9)), on a bead can be obtained by differentiating Equation (S.8):

$$E^S = \frac{1}{2}k_0\sum_{i=2}^{N}(l_i - l_0)^2 \quad \text{(S.8)}$$

$$\boldsymbol{F}_i^S = -\boldsymbol{\nabla}_{r_i}E^S = -k_0(l_i - l_0)\hat{\boldsymbol{t}}_i + k_0(l_{i+1} - l_0)\hat{\boldsymbol{t}}_{i+1} \quad \text{(S.9)}$$

The bending force can also be obtained by differentiating the bending energy on a bead (see Equation (S.10)) and (S.11)):

$$E^B = \frac{A}{l_0}\sum_{i=1}^{N} f_i(1 - \hat{\boldsymbol{t}}_{i+1} \cdot \hat{\boldsymbol{t}}_i) \quad \text{where} \quad f_i = \begin{cases} 1 \; for \; 2 \le i \le N-1 \\ 0 \; for \; i = 1, N \end{cases} \quad \text{(S.10)}$$

$$\begin{aligned}\boldsymbol{F}_i^B = -\boldsymbol{\nabla}_{r_i}E^B = \frac{A}{l_0}\Bigg\{&\frac{f_{i-1}}{l_i}\hat{\boldsymbol{t}}_{i-1} - \left[\frac{f_{i-1}}{l_i}\hat{\boldsymbol{t}}_{i-1} \cdot \hat{\boldsymbol{t}}_i + \frac{f_i}{l_{i+1}} + \frac{f_i}{l_i}\hat{\boldsymbol{t}}_i \cdot \hat{\boldsymbol{t}}_{i+1}\right]\hat{\boldsymbol{t}}_i \\ &+ \left[\frac{f_i}{l_{i+1}}\hat{\boldsymbol{t}}_i \cdot \hat{\boldsymbol{t}}_{i+1} + \frac{f_i}{l_i} + \frac{f_{i+1}}{l_{i+1}}\hat{\boldsymbol{t}}_{i+1} \cdot \hat{\boldsymbol{t}}_{i+2}\right]\hat{\boldsymbol{t}}_{i+1} - \frac{f_{i+1}}{l_{i+1}}\hat{\boldsymbol{t}}_{i+2}\Bigg\}\end{aligned} \quad \text{(S.11)}$$

#### S.1.2 Homogeneous Turbulence

The homogeneous turbulence is modelled with multiple Gaussian random velocity fields, that are proposed by Fung (see Equation (S.12); Kraichnan 1970, Drummond et al. 1984, Turfus 1985, Fung et al. 1992).

$$\boldsymbol{u}(\boldsymbol{r},t) = \sum_{n=1}^{N} \left[ \frac{\boldsymbol{a}_n \times \boldsymbol{k}_n}{|\boldsymbol{k}_n|} cos(\boldsymbol{k}_n \cdot \boldsymbol{r} + \omega_n t) + \frac{\boldsymbol{b}_n \times \boldsymbol{k}_n}{|\boldsymbol{k}_n|} sin(\boldsymbol{k}_n \cdot \boldsymbol{r} + \omega_n t) \right] \quad \text{(S.12)}$$

The subscript $n$ refers to the quantity of the nth Fourier mode. The wavenumber vectors $\boldsymbol{k}_n$ are randomly chosen on each spherical shell of radius $k_n$. The coefficients $\boldsymbol{a}_n$ and $\boldsymbol{b}_n$ are determined by picking independently random vectors from a three-dimensional isotropic Gaussian distribution, with zero mean vector and variance $3\,\gamma \int_{k_n-\delta k/}^{k_n+\delta k/2} E(k)dk$ , where $\gamma = \int_0^{\infty} E(k)dk / \int_0^{k_\eta} E(k)dk$ and $\delta k = (k_\eta - k_c)/(N_k - 1)$ (Fung et al. 1992). Wavenumbers range from $k_c = 1$ to $k_\eta = 10$. For 10 Fourier modes $N_k$, $k_n = k_c + (k_\eta - k_c)(n-1)/(N_k - 1)$, where $N_k = 10$ (Fung et al. 1992). Therefore, the sizes of eddies range from 0.628 to 6.28. Equation (S.13) expresses the von Karman energy spectrum. The numerical constants of $g_1 = 0.558$ and $g_2 = 1.196$ are used for the model (Hunt 1973, Hinze 1987, Fung et al. 1992).

$$E(k) = \frac{g_2 k^4}{(g_1 + k^2)^{17/6}} \quad \text{(S.13)}$$

#### S.1.3 Execution Time

The total number of integration time-steps is calculated by Equations (S.14), (S.15), (S.16)) and (S.17)). If the sedimentation is more dominant than the background velocity fields, the longest period for one cycle of motion is calculated with the characteristic length $L$, dividing it by the smallest possible sediment velocities $|V_{z,still} + U_0|$ and $|V_{z,still} + \sigma|$ (see Equations (S.14) and (S.16))). If the strength of the background velocity field is more dominant than the sedimentation, the perimeters of a vortex in a periodic cellular flow and the largest eddy in the turbulence, $\pi L_c$ and $\pi l_{large\ eddy}$, respectively, are used to calculate the longest period for one cycle of motion, dividing those characteristic lengths by the possible smallest drift velocities $|V_{z,still} + U_0|$ and $|V_{z,still} + \sigma|$ (see Equations (S.15) and (S.17))). Twenty times longer than the period of the one time-revolution is used as simulation time ($f_{vor} = 20$, $f_{tur} = 20$). The total number of integration time-steps is shown in Equations (S.14))-(S.17)).

$$N_{vor} = f_{vor} \frac{L}{|V_{z,still} + U_0|\Delta t} \qquad \text{if } |V_{z,still}| > U_0 \quad \text{(S.14)}$$

$$N_{vor} = f_{vor}\frac{\pi L_c}{|V_{z,still} + U_0|\Delta t} \qquad \text{if } |V_{z,still}| < U_0 \tag{S.15}$$

$$N_{tur} = f_{tur}\frac{L}{|V_{z,still} + \sigma|\Delta t} \qquad \text{if } |V_{z,still}| > \sigma \tag{S.16}$$

$$N_{tur} = f_{tur}\frac{\pi l_{large\ eddy}}{|V_{z,still} + \sigma|\Delta t} \qquad \text{if } |V_{z,still}| < \sigma \tag{S.17}$$

### S.2 Validation

To validate the anisotropic character of the bead model, we compared the results of hydrodynamic simulation with the results of the approximate method proposed by Burgers (1938, Happel and Brenner 1991) and experimental data. The experiment is implemented using straight and cylindrical steel wires with thicknesses ranging from 0.3-0.8 mm. The drag forces of these steel wires are measured in both directions (transverse, in the direction perpendicular to the axis, and longitudinally, in the direction of the axis) by dropping the wires into corn syrup with viscosities ranging from 2-3 kg/ms. The simulation values coincide very well with those of the experiments and the theoretical values. The experiment shows the 1.3-2.2 factors of the ratio between a transverse drag and a longitudinal drag. The values from the approximated method indicate the factors of 1.5-1.6. The simulation shows the factors of 1.3-1.5. This ratio of the drag on a wire moving perpendicular to the axis to the drag on a wire moving parallel to its length is supposed to be a factor of 2, however Purcell proposed that it may be around 1.5 from his experiment (Purcell 1977). Although there are large differences in the slenderness of filaments between real ballooning phenomena and our model, a simulation with slenderness of 45 to 120 can describe the anisotropic character of a filament in a low Reynolds flow and can also be used for the study of fibre motion in various fluid flows (Lauga and Powers 2009).

The non-dimensional resistance coefficients, which are based on Burgers' approximated methods (Eqns. (S.18)) and (S.19))), are calculated by means of Eqns. (S.20)) and (S.21)). The non-dimensional resistance coefficients differ from the translation tensor, which has length scale (Happel and Brenner 1991).

$$F_N = \frac{4\pi\eta UL}{\left(ln\left(2\frac{L}{d}\right) + 0.5\right)} \tag{S.18}$$

$$F_T = \frac{2\pi\eta UL}{\left(ln\left(2\frac{L}{d}\right) - 0.72\right)} \tag{S.19}$$

$$R_N = \frac{F_N}{\eta UL} \tag{S.20}$$

$$R_T = \frac{F_T}{\eta U L} \tag{S.21}$$

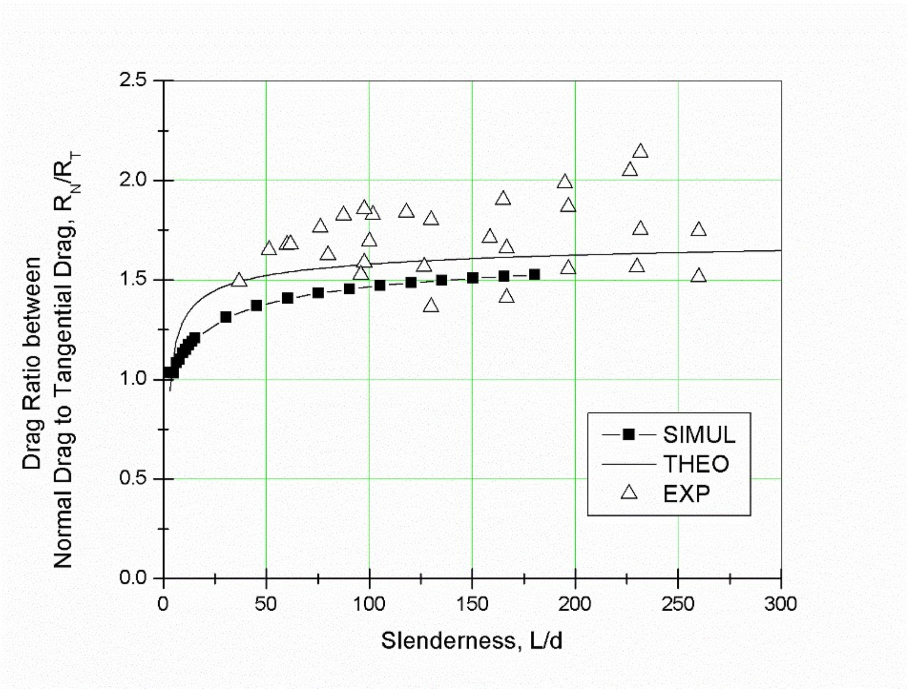


Figure S1. Drag ratio of the transverse resistance force to the longitudinal resistance force of a thin cylindrical rod. (Cho 2020)

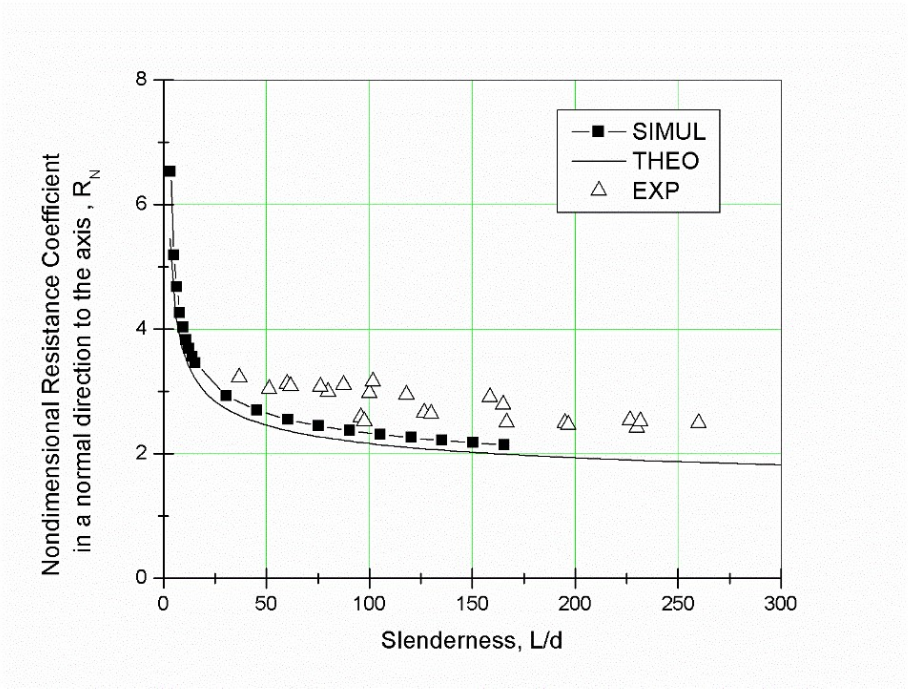


Figure S2. Dimensionless transverse (normal) resistance coefficients of a thin cylinder as a function of the slenderness of the cylinder. (Cho 2020)

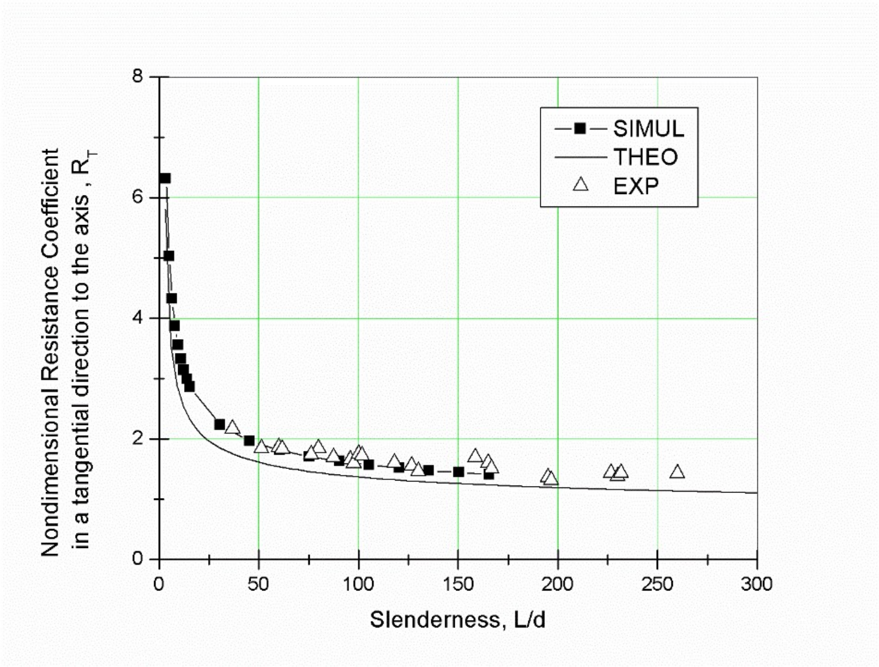


Figure S3. Dimensionless longitudinal (tangential) resistance coefficients of a thin cylinder as a function of the slenderness of the cylinder. (Cho 2020)

### S.3 Input Parameters

#### S.3.1 Homogeneous Turbulence

The ballooning number is varied by composing different parameters, e.g. the length of the structure, the viscosity of the medium, the turbulent kinetic energy and its weight. Table S1 shows the input parameters which are used in the simulation. The elastic constant of the tensile spring is increased, because the elongation of a spider silk is limited to the 30% of the initial length (Ko et al. 2001, Bonino 2003). The divergence problem of the bead-spring model in the turbulent flow field is corrected by adjusting the bending stiffness of the bending spring to 0.1 Nm.

Table S1. Input parameters for a homogeneous turbulence.

| | Values |
|---|---|
| $Bn_{tur}$ | 0.022 - 445 |
| $n$ | 20, 30, 40 |
| $TKE$ $[m^2/s^2]$ | 0.23 - 2.45 |
| $\sigma$ | 0.68 - 2.22 |
| $\eta$ $[kg/(m \cdot s)]$ | 0.1, 0.2, 0.5, 1, 2, 5 |
| $a$ $[m]$ | 0.1 |
| $a_1$ | $a$ |
| $a_2, \cdots, a_n$ | $a$ |
| $g$ $[kg\ m/s^2]$ | 9.81 |
| $l_0$ | $3a$ |
| $w_1$ $[kg]$ | 0.01, 0.02, 0.05, 0.1, 0.2 0. 5, 1 |
| $\rho_2, \cdots, \rho_n$ $[kg/m^3]$ | 1.225 |
| $\rho_f$ $[kg/m^3]$ | 1.225 |
| $k$ $[N/m]$ | 300 |
| A $[Nm]$ | 0.1 |
| $\Delta t$ $[sec]$ | 0.001 |

#### S.3.2 Shear Flow

Table S2 shows input parameter sets for a shear flow. Different sets and combinations of input parameters are used, in order to cover the wide range of the ballooning number. The sets of 1-3 in Table S2 are used for the model with the hydrodynamic interaction between beads. The set 4 is used for the model without with the hydrodynamic interaction between beads. The variation of the spring constants is applied in order not to exceed elongation of 1.2 of the chain, because the drag anisotropy diminishes as the distance between beads increases.

Table S2. Input parameters for a shear flow.

| | **Set 1** | **Set 2** | **Set 3** | **Set 4 (No Hydro.)** |
|---|---|---|---|---|
| $Bn_{shear}$ | 0.011 - 57 | 0.042 - 1133 | 1.9 - 4539 | 0.047 - 155 |
| $n$ | 20, 30, 40 | 40, 60 | 40, 80 | 30, 40, 50 |
| $\gamma$ [/s] | 0.1, 0.2, 0.5, 1, 2, 5 | 0.1, 0.2, 0.5, 1, 2, 5 | 1, 2, 5, 10 | 0.1, 0.2, 0.5, 1, 2, 5 |
| $\eta$ $[kg/(m \cdot s)]$ | 0.1, 0.2, 0.5, 1, 2, 5 | 0.1, 0.2, 0.5, 1, 2, 5 | 0.5, 1, 2, 5, 10 | 0.1, 0.2, 0.5, 1, 2, 5 |
| $a$ $[m]$ | 0.1 | 0.1 | 0.1 | 0.1 |
| $a_1$ | $a$ | $a$ | $a$ | $a$ |
| $a_2, \cdots, a_n$ | $a$ | $a$ | $a$ | $a$ |
| $g$ $[kg\, m/s^2]$ | 9.81 | 9.81 | 9.81 | 9.81 |
| $l_0$ | $3a$ | $3a$ | $3a$ | $3a$ |
| $w_1$ $[kg]$ | $0.1, 0.2, 0.5, 1$ | $0.1, 0.2, 0.5, 1$ | $0.1, 0.2, 0.5, 1$ | $0.1, 0.2, 0.5, 1$ |
| $\rho_2, \cdots, \rho_n$ $[kg/m^3]$ | 1.225 | 1.225 | 1.225 | 1.225 |
| $\rho_f$ $[kg/m^3]$ | 1.225 | 1.225 | 1.225 | 1.225 |
| $k$ $[N/m]$ | 200 | 300 | 1000 | 200 |
| A $[Nm]$ | $1.0 \times 10^{-4}$ | $1.0 \times 10^{-4}$ | $1.0 \times 10^{-4}$ | $1.0 \times 10^{-4}$ |
| $\Delta t$ $[sec]$ | 0.001 | 0.001 | 0.0005 | 0.001 |

#### S.3.3 Periodic Cellular Flow

The wide range of the ballooning number is achieved by varying the characteristic velocity of the periodic cellular flow and the weight of the ballooning structure. The input parameters are summarised in Table S3. The ratio of the filament length to the diameter of a vortex cell is additionally considered by simulating in various sizes of cellular vortex fields. The three different x-axis positions are employed to observe the dependency of the initial condition at the release.

Table S3. Input parameters for a periodic cellular flow.

| | **Values** |
|---|---|
| $Bn_{vor}$ | 0.064 - 3.42 |
| $n$ | 30 |
| $U_0$ [m/s] | 0.25, 0.5, 1, 1.5 |
| $L_c$ | 0.25, 0.5, 0.75, 1, 1.25, 1.5, 2 $\times L$ |
| $x_0$ | 0.25, 0.5, 0.75 $\times L$ |
| $\eta$ $[kg/(m \cdot s)]$ | 0.5 |
| $a$ $[m]$ | 0.1 |
| $a_1$ | $a$ |
| $a_2, \cdots, a_n$ | $a$ |
| $g$ $[kg\, m/s^2]$ | 9.81 |
| $l_0$ | $3a$ |
| $w_1$ $[kg]$ | $0.1, 0.2, 0.5, 1$ |
| $\rho_2, \cdots, \rho_n$ $[kg/m^3]$ | 1.225 |
| $\rho_f$ $[kg/m^3]$ | 1.225 |
| $k$ $[N/m]$ | 200 |
| A $[Nm]$ | $1.0 \times 10^{-4}$ |
| $\Delta t$ $[sec]$ | 0.002 |

### S.4 Dynamic Motions

#### S.4.1 Dynamic Motions in Homogeneous Turbulence

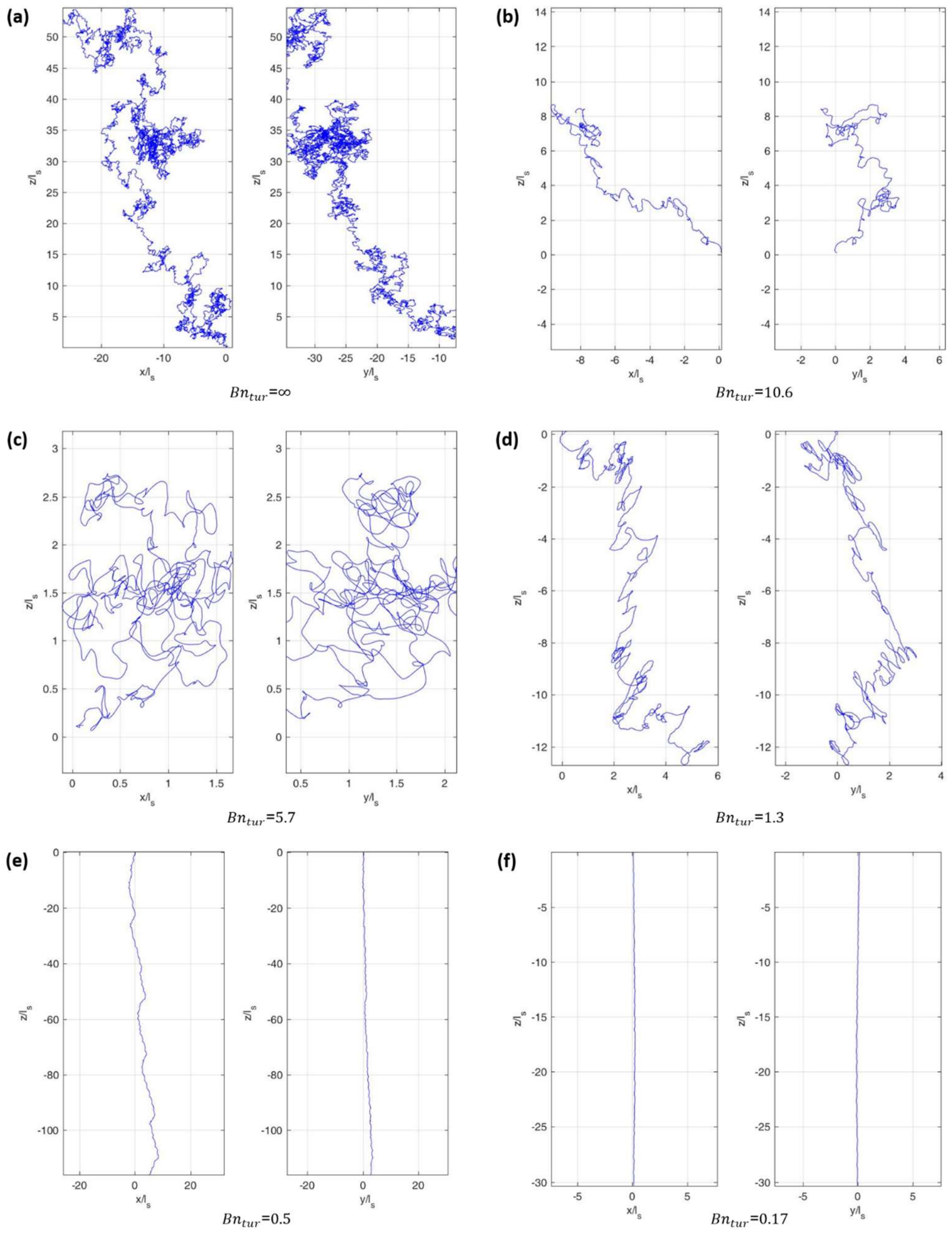


Figure S4. Trace plots in homogeneous turbulence. (the number of beads: 30, length: 4.45 m, viscosity: 0.2 kg/ms) (a) $Bn_{tur} = \infty$ (mass: $0.61 \times 10^{-4}\ kg$, $\sigma = 0.578$, $u' = 0.381$, $v' = 0.799$, $w' = 0.471$), (b) $Bn_{tur} = 10.6$ (mass: $0.05\ kg$, $\sigma = 0.713$, $u' = 0.633$, $v' = 0.772$, $w' = 0.725$), (c) $Bn_{tur} = 5.7$ (mass: $0.02\ kg$, $\sigma = 0.714$, $u' = 0.510$, $v' = 0.860$, $w' = 0.728$), (d) $Bn_{tur} = 1.3$ (mass: $0.1\ kg$, $\sigma = 0.831$, $u' = 0.735$, $v' = 0.686$, $w' = 1.030$), (e) $Bn_{tur} = 0.5$ (mass: $0.2\ kg$, $\sigma = 0.647$, $u' = 0.712$, $v' = 0.477$, $w' = 0.722$), (f) $Bn_{tur} = 0.17$ (mass: $1\ kg$, $\sigma = 1.012$, $u' = 1.118$, $v' = 1.251$, $w' = 0.509$) ; Unit of $\sigma, u', v', w'$: $[m/s]$

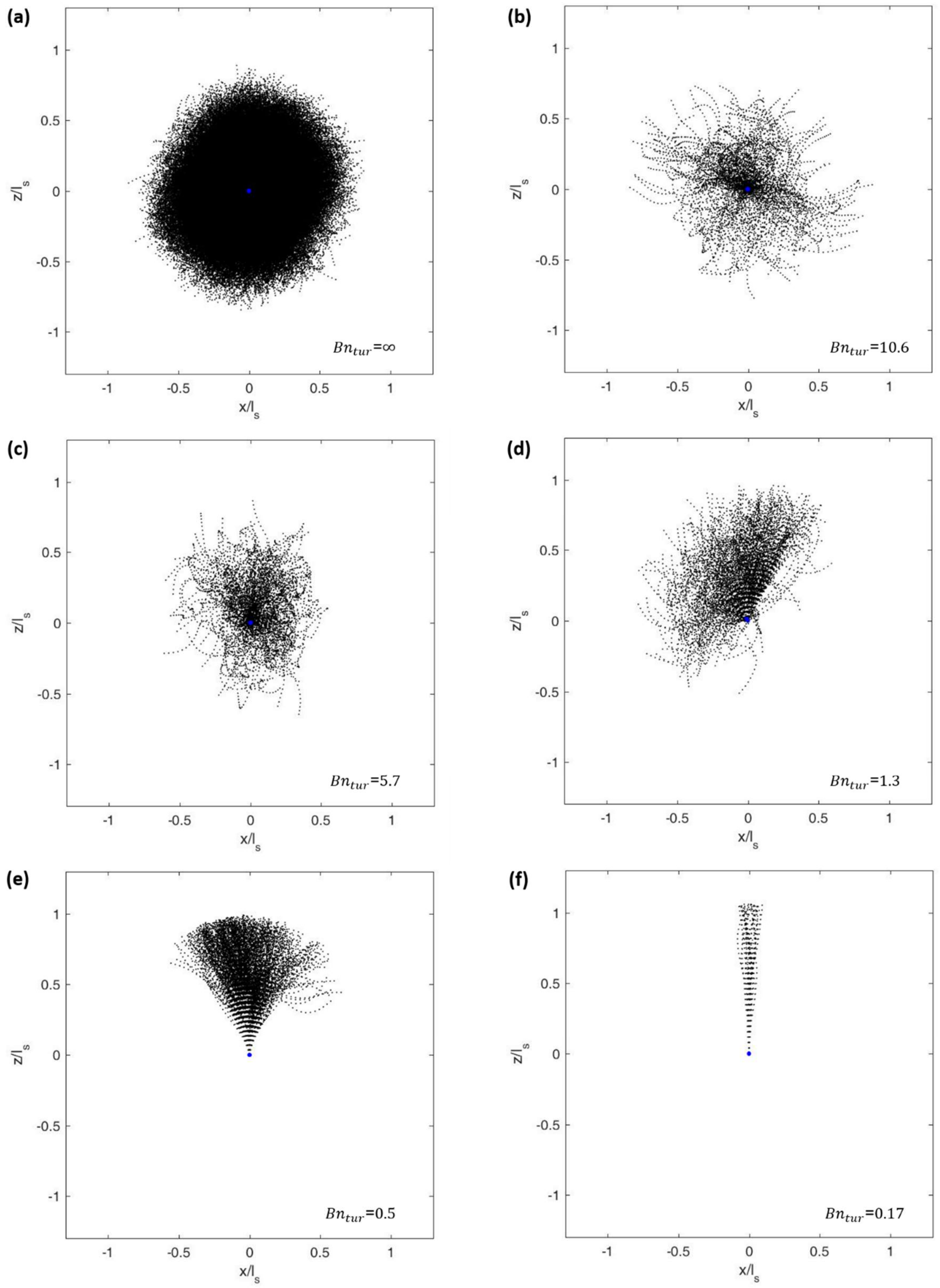


Figure S5. Projection plots of the filament structure in homogeneous turbulence in the relative coordinate of the lower end. (the number of beads: 30, length: 4.45 m, viscosity: 0.2 kg/ms) (a) $Bn_{tur} = \infty$ (mass: $0.61 \times 10^{-4}\,kg$, $\sigma = 0.578$, $u' = 0.381$, $v' = 0.799$, $w' = 0.471$), (b) $Bn_{tur} = 10.6$ (mass: $0.05\,kg$, $\sigma = 0.713$, $u' = 0.633$, $v' = 0.772$, $w' = 0.725$), (c) $Bn_{tur} = 5.7$ (mass: $0.02\,kg$, $\sigma = 0.714$, $u' = 0.510$, $v' = 0.860$, $w' = 0.728$), (d) $Bn_{tur} = 1.3$ (mass: $0.1\,kg$, $\sigma = 0.831$, $u' = 0.735$, $v' = 0.686$, $w' = 1.030$), (e) $Bn_{tur} = 0.5$ (mass: $0.2\,kg$, $\sigma = 0.647$, $u' = 0.712$, $v' = 0.477$, $w' = 0.722$), (f) $Bn_{tur} = 0.17$ (mass: $1\,kg$, $\sigma = 1.012$, $u' = 1.118$, $v' = 1.251$, $w' = 0.509$) ; Unit of $\sigma, u', v', w'$: $[m/s]$

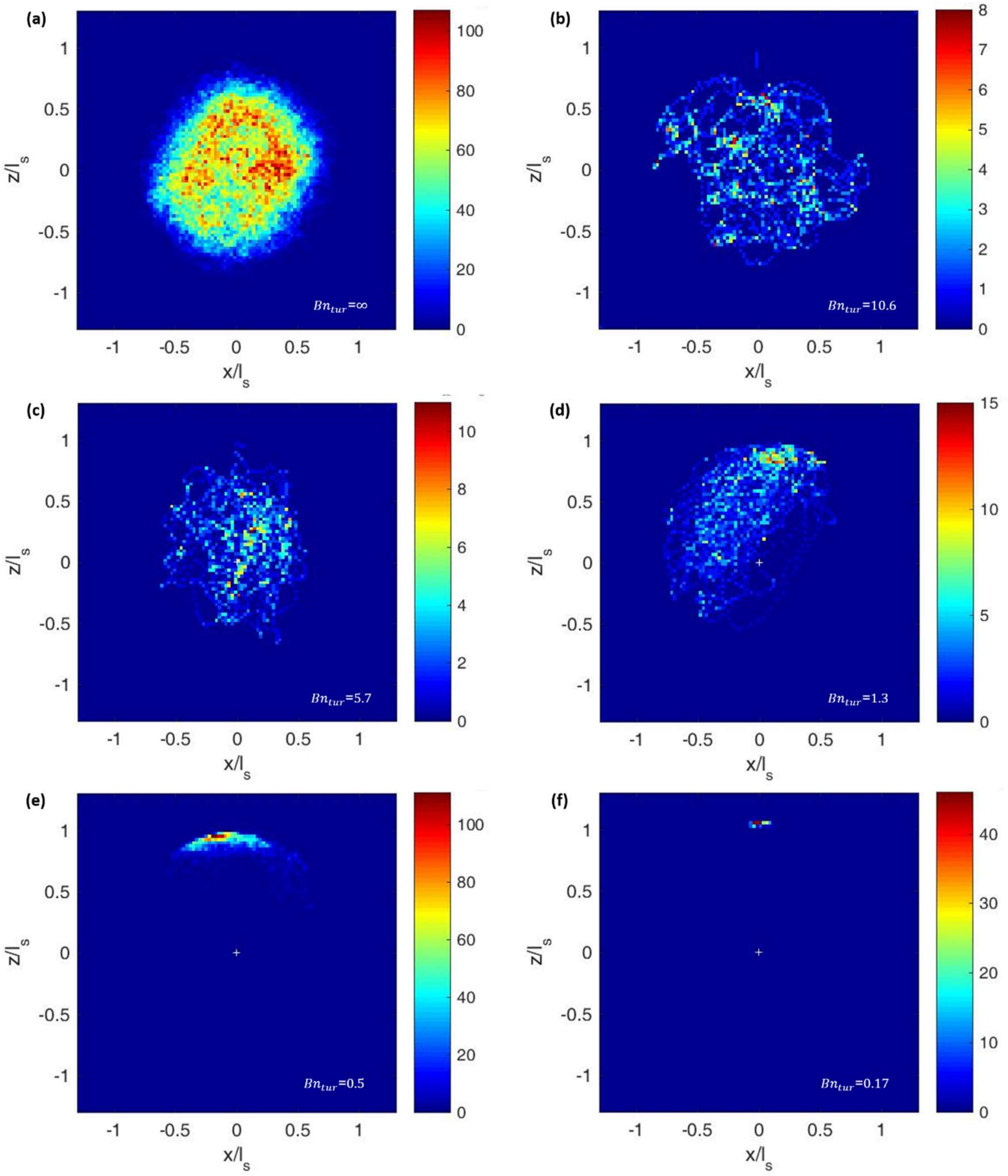


Figure S6. Projected spatial frequency of the end point of the filament. (the number of beads: 30, length: 4.45 m, viscosity: 0.2 kg/ms) (a) $Bn_{tur} = \infty$ (mass: $0.61 \times 10^{-4}\ kg$, $\sigma = 0.578$, $u' = 0.381$, $v' = 0.799$, $w' = 0.471$), (b) $Bn_{tur} = 10.6$ (mass: $0.05\ kg$, $\sigma = 0.713$, $u' = 0.633$, $v' = 0.772$, $w' = 0.725$), (c) $Bn_{tur} = 5.7$ (mass: $0.02\ kg$, $\sigma = 0.714$, $u' = 0.510$, $v' = 0.860$, $w' = 0.728$), (d) $Bn_{tur} = 1.3$ (mass: $0.1\ kg$, $\sigma = 0.831$, $u' = 0.735$, $v' = 0.686$, $w' = 1.030$), (e) $Bn_{tur} = 0.5$ (mass: $0.2\ kg$, $\sigma = 0.647$, $u' = 0.712$, $v' = 0.477$, $w' = 0.722$), (f) $Bn_{tur} = 0.17$ (mass: $1\ kg$, $\sigma = 1.012$, $u' = 1.118$, $v' = 1.251$, $w' = 0.509$) ; Unit of $\sigma, u', v', w'$: $[m/s]$

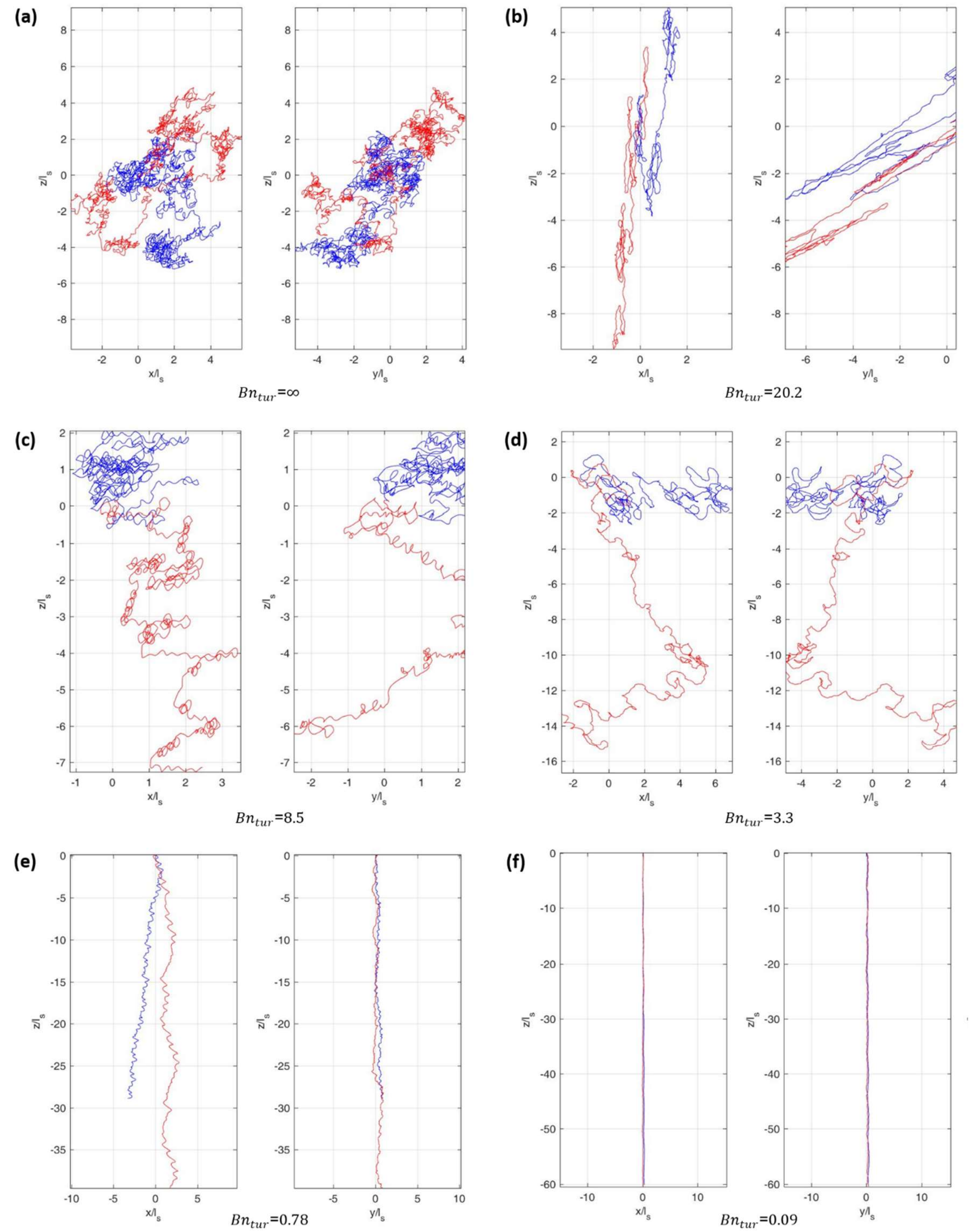


Figure S7. Trace plots of two different ballooners in an identical homogeneous turbulence. (blue line: a filament-like ballooner, red line: a point-like ballooner; the number of beads: 30, length: 4.45 m, viscosity: 0.2 kg/ms) (a) $Bn_{tur} = \infty$ (mass: $0.61 \times 10^{-4}\ kg$, $\sigma = 0.858$, $u' = 0.803$, $v' = 0.884$, $w' = 0.883$), (b) $Bn_{tur} = 20.2$ (mass: $0.01\ kg$, $\sigma = 1.229$, $u' = 0.475$, $v' = 1.764$, $w' = 1.093$), (c) $Bn_{tur} = 8.5$ (mass: $0.02\ kg$, $\sigma = 1.072$, $u' = 0.984$, $v' = 1.085$, $w' = 1.140$), (d) $Bn_{tur} = 3.3$ (mass: $0.05\ kg$, $\sigma = 1.063$, $u' = 1.106$, $v' = 1.130$, $w' = 0.944$), (e) $Bn_{tur} = 0.78$ (mass: $0.1\ kg$, $\sigma = 0.503$, $u' = 0.593$, $v' = 0.548$, $w' = 0.330$), (f) $Bn_{tur} = 0.09$ (mass: $1\ kg$, $\sigma = 0.554$, $u' = 0.364$, $v' = 0.524$, $w' = 0.716$) ; Unit of $\sigma, u', v', w'$: $[m/s]$

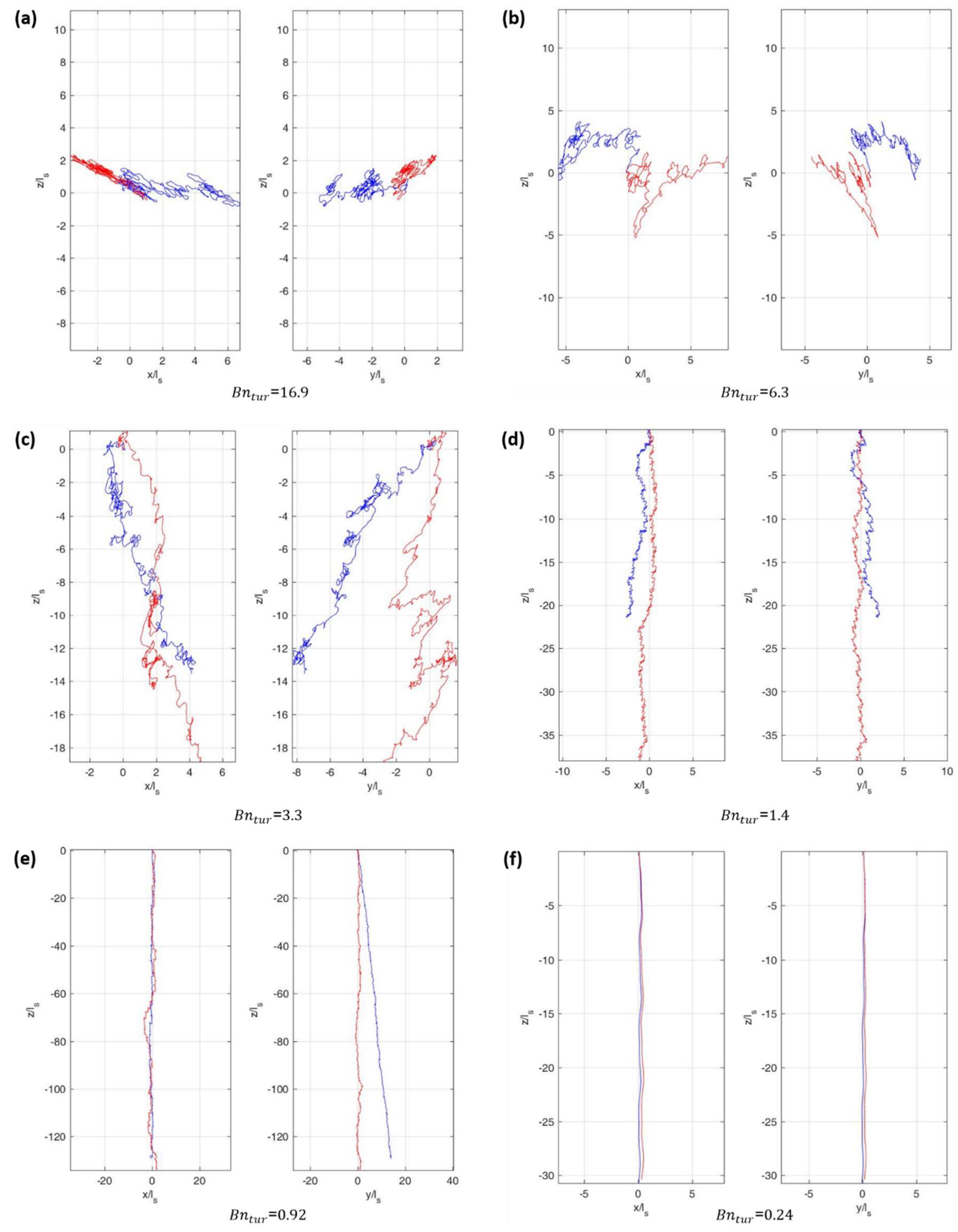


Figure S8. Trace plots of two different ballooners in an identical homogeneous turbulence. (blue line: a filament-like ballooner, red line: a point-like ballooner; the number of beads: 30, length: 4.45 m, viscosity: 0.2 kg/ms) (a) $Bn_{tur} = 16.9$ (mass: $0.01\,kg$, $\sigma = 1.034$, $u' = 1.365$, $v' = 0.757$, $w' = 0.879$), (b) $Bn_{tur} = 6.3$ (mass: $0.02\,kg$, $\sigma = 0.793$, $u' = 0.695$, $v' = 0.605$, $w' = 1.019$), (c) $Bn_{tur} = 3.3$ (mass: $0.05\,kg$, $\sigma = 1.060$, $u' = 0.849$, $v' = 0.947$, $w' = 1.325$), (d) $Bn_{tur} = 1.4$ (mass: $0.1\,kg$, $\sigma = 0.889$, $u' = 0.889$, $v' = 0.804$, $w' = 0.967$), (e) $Bn_{tur} = 0.92$ (mass: $0.2\,kg$, $\sigma = 1.177$, $u' = 1.130$, $v' = 0.981$, $w' = 1.384$), (f) $Bn_{tur} = 0.24$ (mass: $0.5\,kg$, $\sigma = 0.761$, $u' = 0.860$, $v' = 0.615$, $w' = 0.786$) ; Unit of $\sigma, u', v', w'$: $[m/s]$

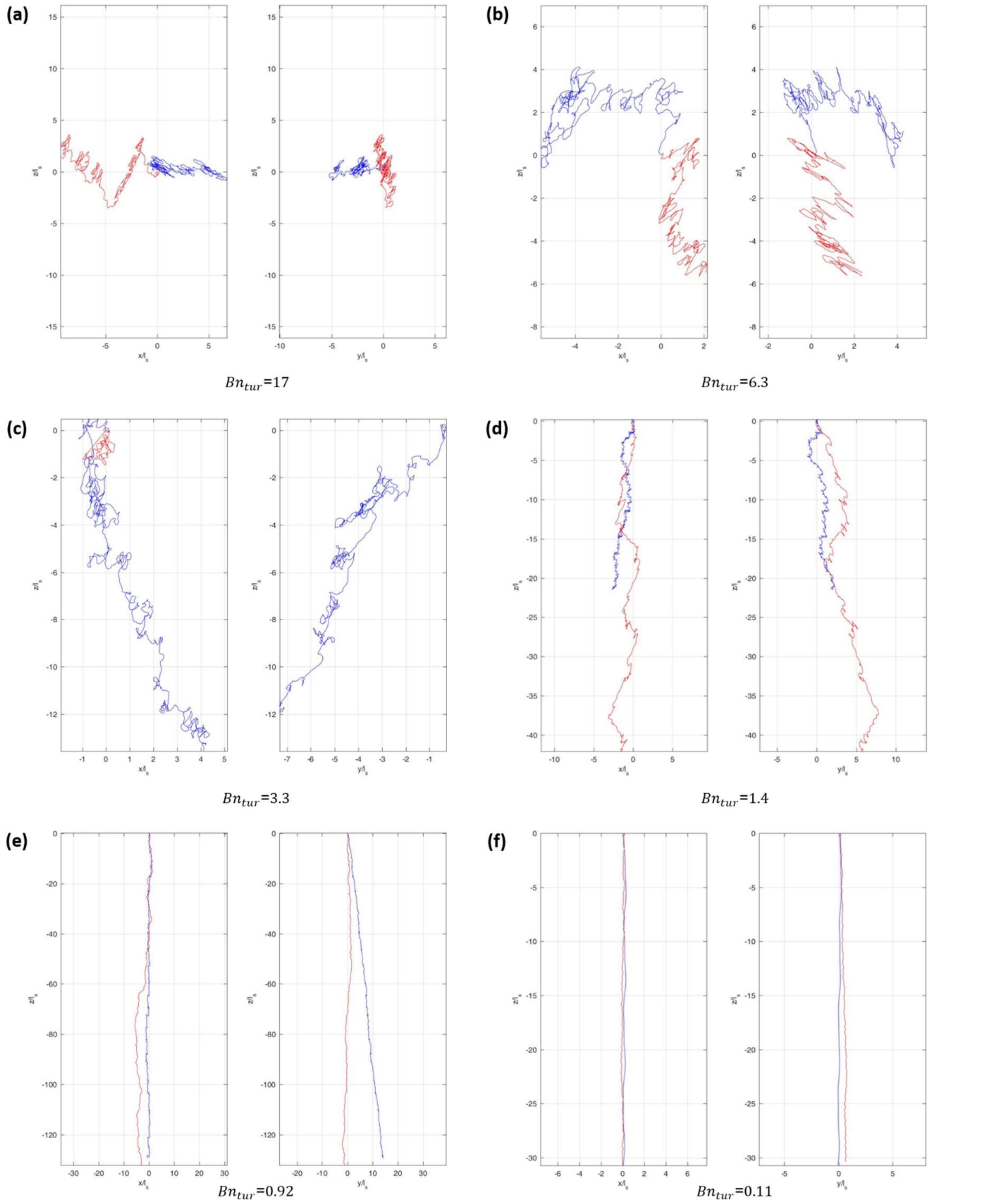


Figure S9. Trace plots of two different ballooners in an identical homogeneous turbulence. (blue line: a filament-like ballooner, red line: a point-like ballooner; the number of beads: 30, length: 4.45 m, viscosity: 0.2 kg/ms) (a) $Bn_{tur} = 17$ (mass: $0.01\,kg$, $\sigma = 1.034$, $u' = 1.365$, $v' = 0.757$, $w' = 0.879$), (b) $Bn_{tur} = 6.29$ (mass: $0.02\,kg$, $\sigma = 0.793$, $u' = 0.695$, $v' = 0.605$, $w' = 1.019$), (c) $Bn_{tur} = 3.3$ (mass: $0.05\,kg$, $\sigma = 1.060$, $u' = 0.849$, $v' = 0.947$, $w' = 1.325$), (d) $Bn_{tur} = 1.4$ (mass: $0.1\,kg$, $\sigma = 0.889$, $u' = 0.889$, $v' = 0.804$, $w' = 0.967$), (e) $Bn_{tur} = 0.92$ (mass: $0.2\,kg$, $\sigma = 1.177$, $u' = 1.130$, $v' = 0.981$, $w' = 1.384$), (f) $Bn_{tur} = 0.11$ (mass: $1\,kg$, $\sigma = 0.637$, $u' = 0.479$, $v' = 0.844$, $w' = 0.524$) ; Unit of $\sigma, u', v', w'$: $[m/s]$

#### S.4.2 Dynamic Motions in a Shear Flow

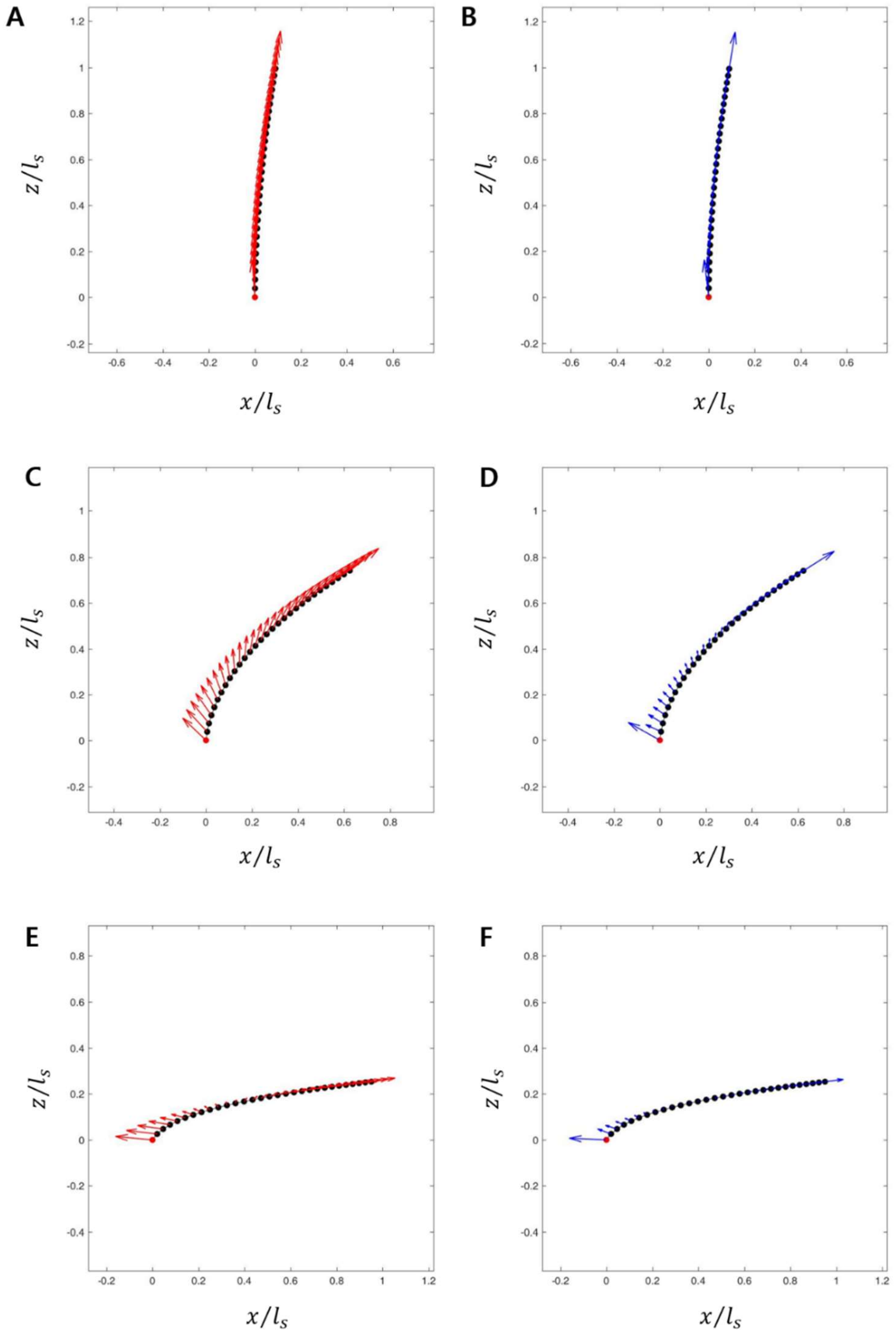


Figure S10. Relative flow velocity and resistance force vector distributions on the ballooning structure in a shear flow. (A) Velocity vector distribution when $Bn_{shear}$= 0.104; (B) Force vector distribution when $Bn_{shear}$= 0.104; (C) Velocity vector distribution when $Bn_{shear}$= 1.4; (D) Force vector distribution when $Bn_{shear}$= 1.4. (E) Velocity vector distribution when $Bn_{shear}$= 20.7; (F) Force vector distribution when $Bn_{shea}$ =20.7. (Red dots indicate weight, while black dots describe a flexible filament structure.)

#### S.4.3 Dynamic Motions in a Periodic Cellular Flow

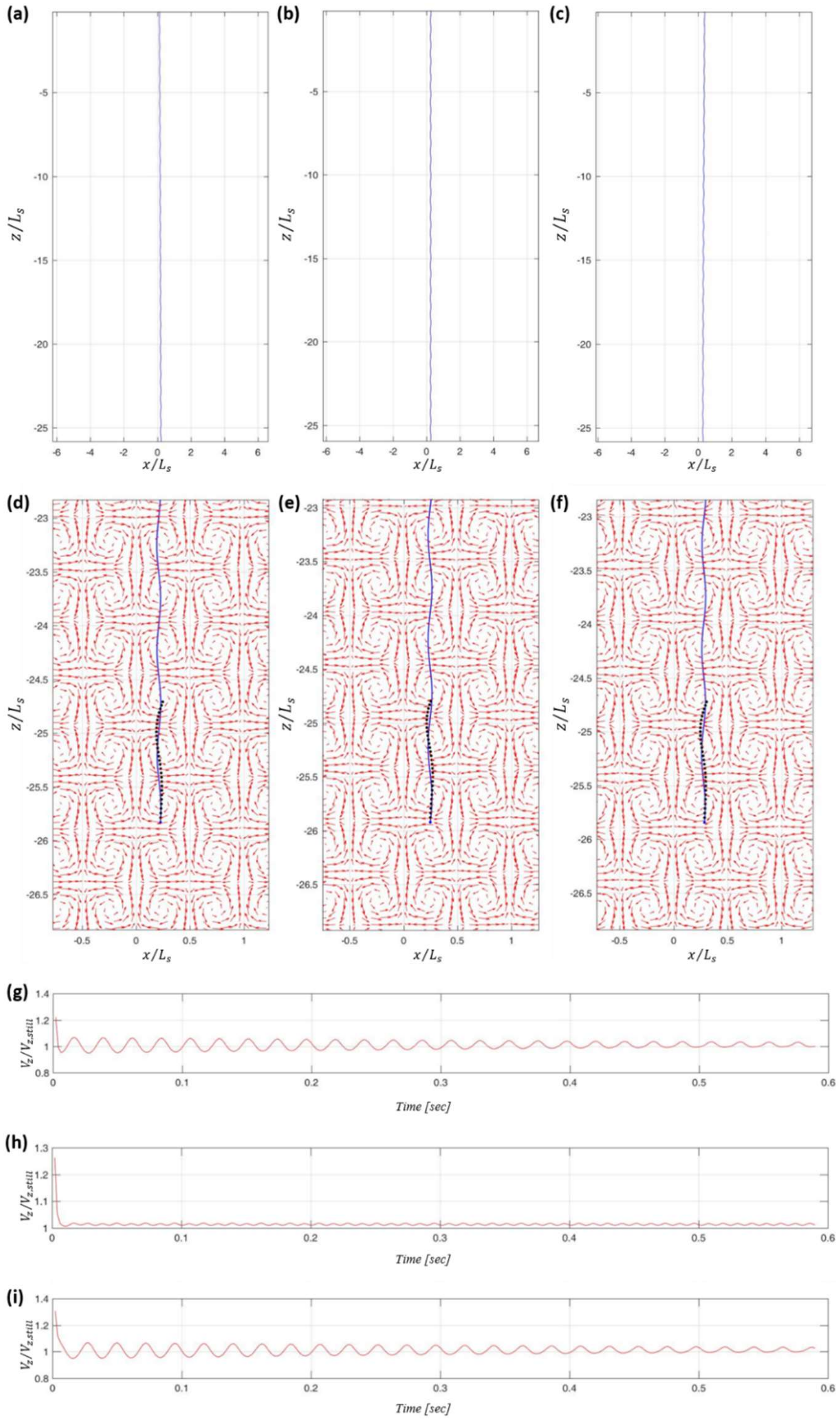

Figure S11. Eddy crossing $\langle V_z \rangle / V_{z,still} = 1.014$; (a)-(c): Trace plots of a point mass (spider body). (d)-(f): Flow fields and the bead-spring model. (blue lines: trajectory plots). (g)-(i): Nondimensionalised settling speeds over time. $Bn_{vor} = 0.13$ (the number of beads: 30, length: 4.45 m, mass: 1 kg, viscosity: 0.5 kg/ms, vortex velocity: $U_0 = 0.5\ m/s$, size of the vortex cell: $L_c/L_s = 0.5$; (a),(d),(g): $x_0/L_c = 0.25$; (b),(e),(h): $x_0/L_c = 0.5$; (c),(f),(i): $x_0/L_c = 0.75$)

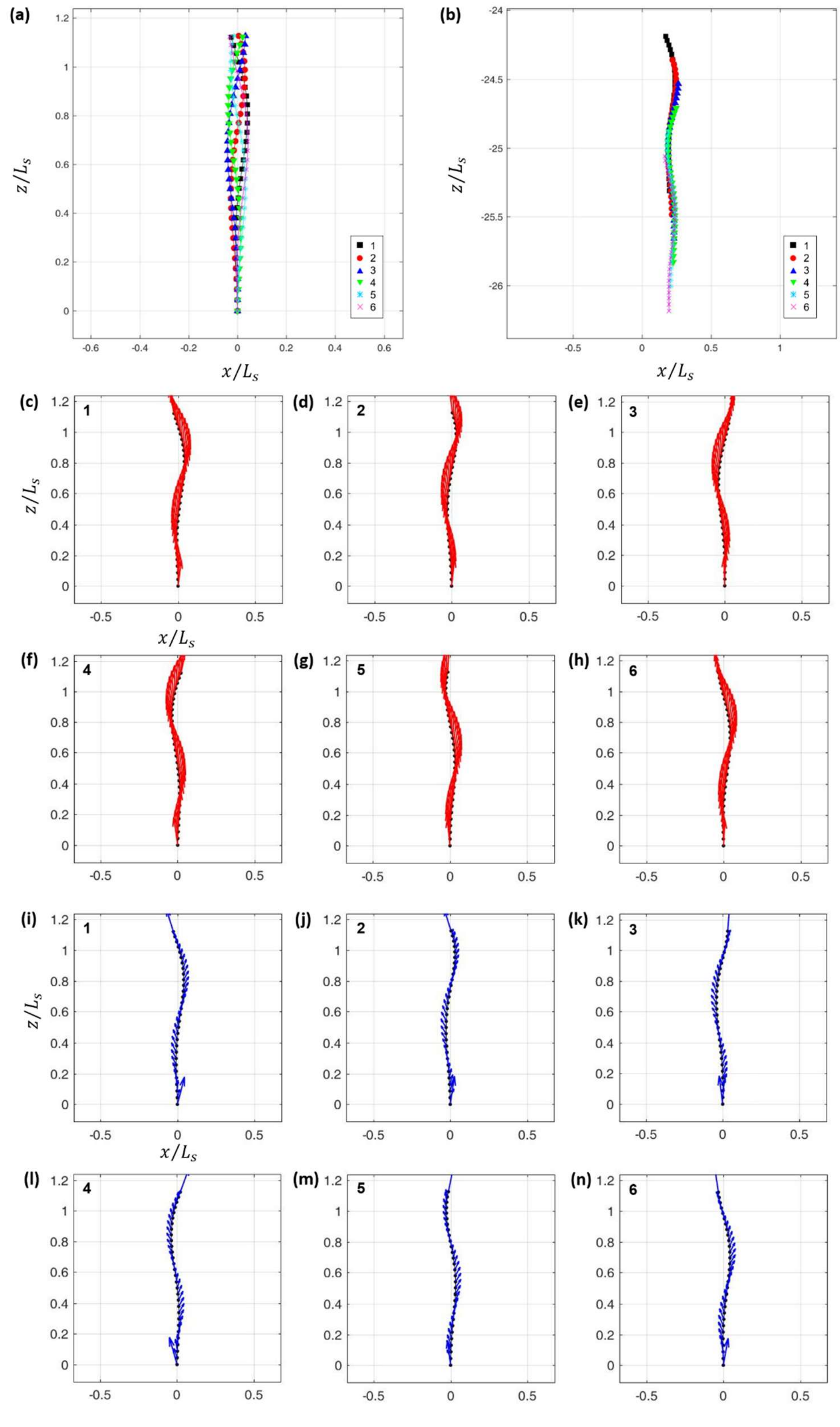


Figure S12. Eddy crossing $\langle V_z \rangle / V_{z,still} = 1.014$; (a) Shape of a filament during sedimentation in relative coordinate of the lower end. (b) Shape of a filament during sedimentation in absolute coordinate. (c)-(h): Distribution of the relative velocity vectors of the flow. (i)-(n): Distribution of the force vectors by the flow. $Bn_{vo} = 0.13$ (the number of beads: 30, length: 4.45 m, mass: 1 kg, viscosity: 0.5 kg/ms, vortex velocity: $U_0 = 0.5\ m/s$, size of the vortex cell: $L_c/L_s = 0.5$)

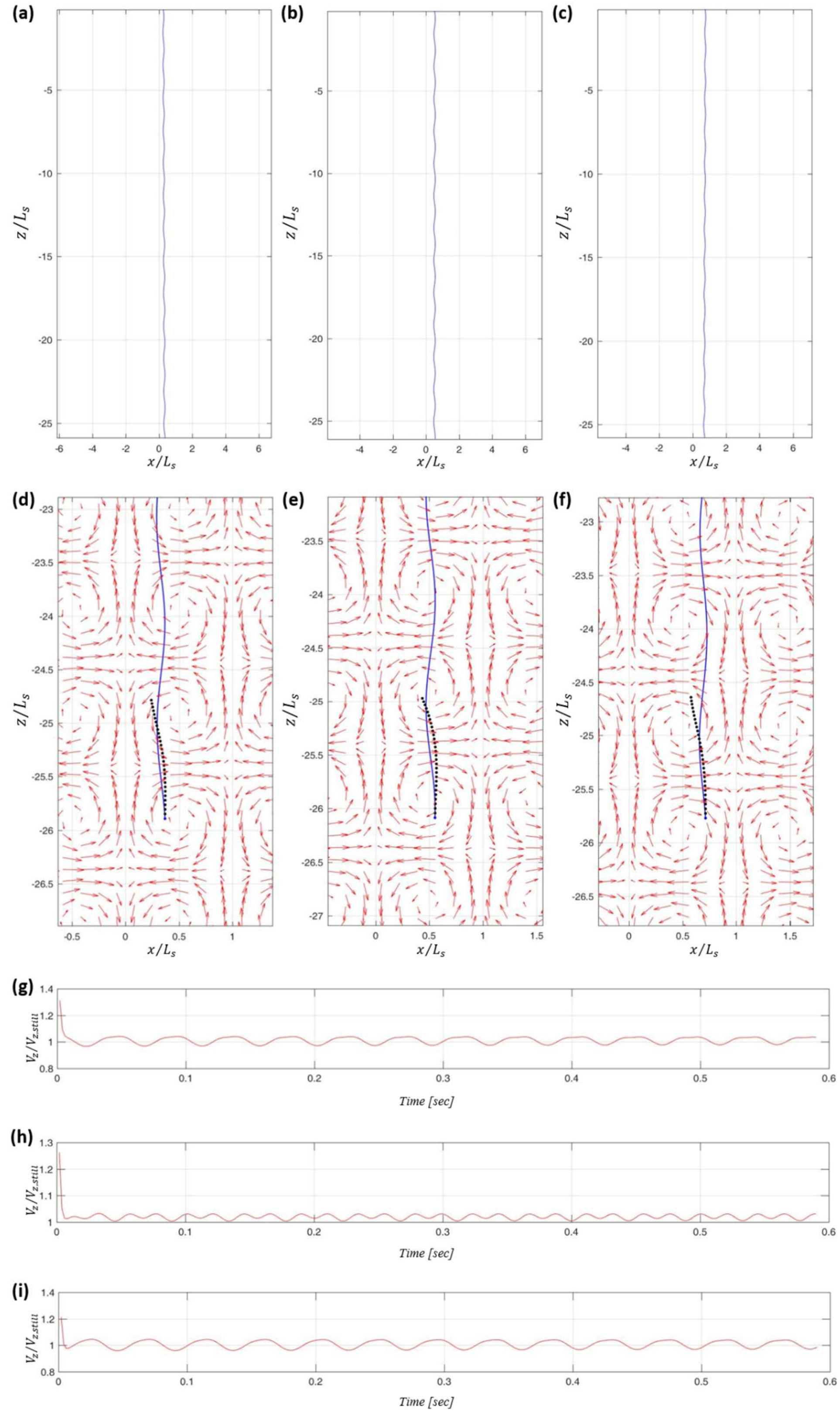


Figure S13. Eddy crossing $\langle V_z \rangle / V_{z,still} = 1.015$; (a)-(c): Trace plots of a point mass (spider body). (d)-(f): Flow fields and the bead-spring model. (blue lines: trajectory plots). (g)-(i): Nondimensionalised settling speeds over time. $Bn_{vor} = 0.13$ (the number of beads: 30, length: 4.45 m, mass: 1 kg, viscosity: 0.5 kg/ms, vortex velocity: $U_0 = 0.5\ m/s$, size of the vortex cell: $L_c/L_s = 1$; (a),(d),(g): $x_0/L_c = 0.25$; (b),(e),(h): $x_0/L_c = 0.5$; (c),(f),(i): $x_0/L_c = 0.75$)

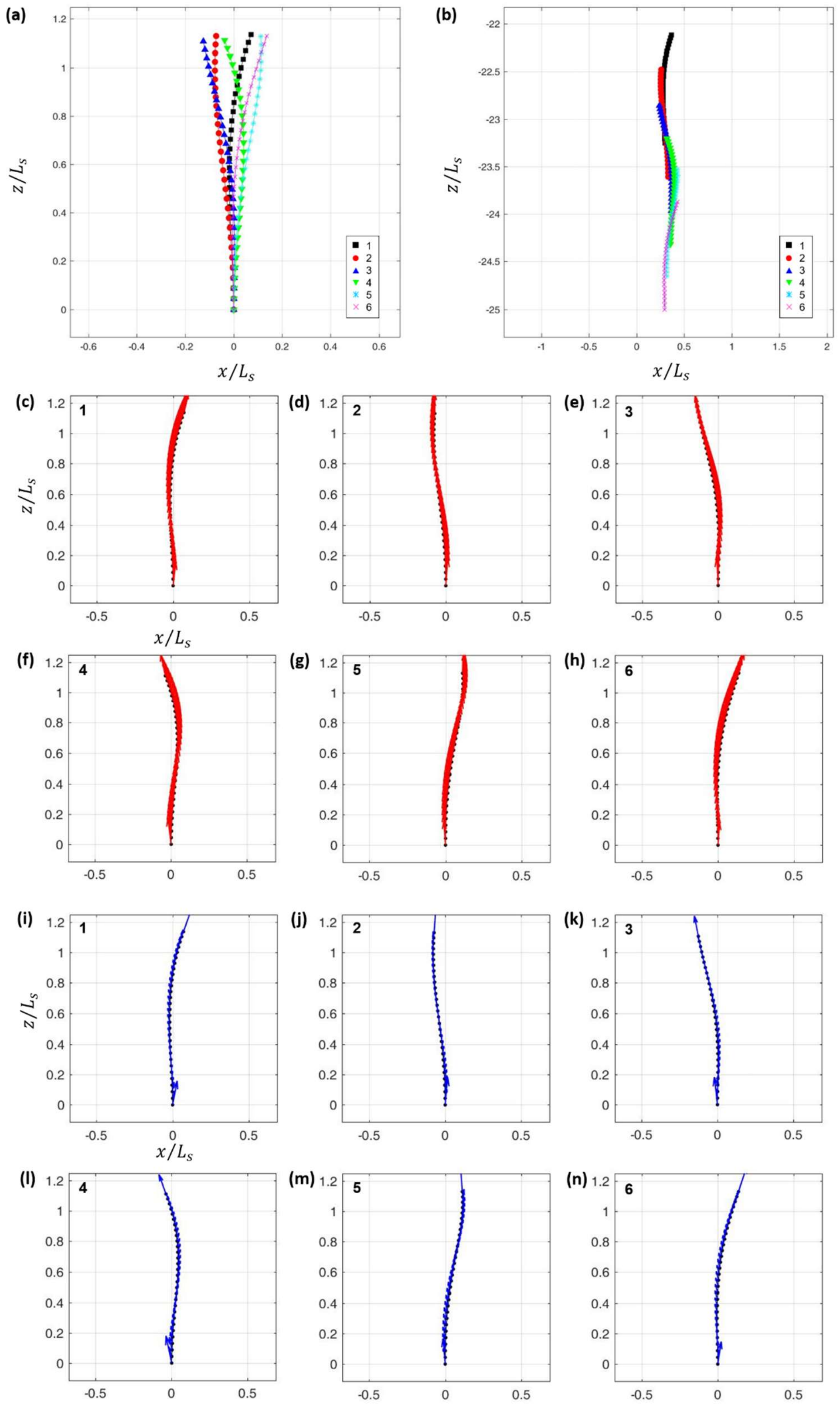

Figure S14. Eddy crossing $\langle V_z \rangle / V_{z,still} = 1.015$; (a) Shape of a filament during sedimentation in relative coordinate of the lower end. (b) Shape of a filament during sedimentation in absolute coordinate. (c)-(h): Distribution of the relative velocity vectors of the flow. (i)-(n): Distribution of the force vectors by the flow. $Bn_{vor} = 0.13$ (the number of beads: 30, length: 4.45 m, mass: 1 kg, viscosity: 0.5 kg/ms, vortex velocity: $U_0 = 0.5\ m/s$, size of the vortex cell: $L_c/L_s = 1$)

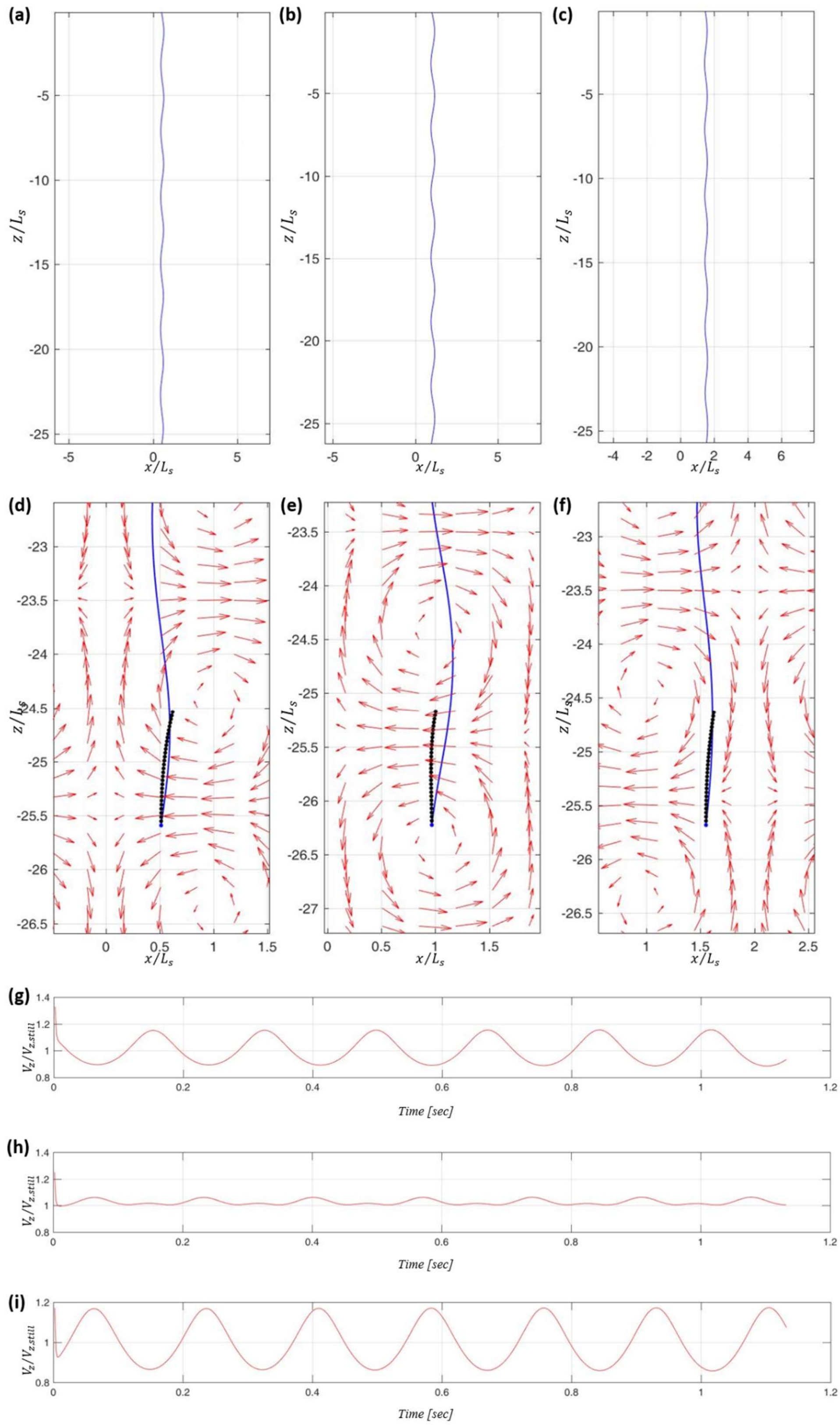


Figure S15. Eddy crossing $\langle V_z\rangle/V_{z,stil} = 1.001$; (a)-(c): Trace plots of a point mass (spider body). (d)-(f): Flow fields and the bead-spring model. (blue lines: trajectory plots). (g)-(i): Nondimensionalised settling speeds over time. $Bn_{vor} = 0.13$ (the number of beads: 30, length: 4.45 m, mass: 1 kg, viscosity: 0.5 kg/ms, vortex velocity: $U_0 = 0.5\ m/s$, size of the vortex cell: $L_c/L_s = 2$; (a),(d),(g): $x_0/L_c = 0.25$; (b),(e),(h): $x_0/L_c = 0.5$; (c),(f),(i): $x_0/L_c = 0.75$)

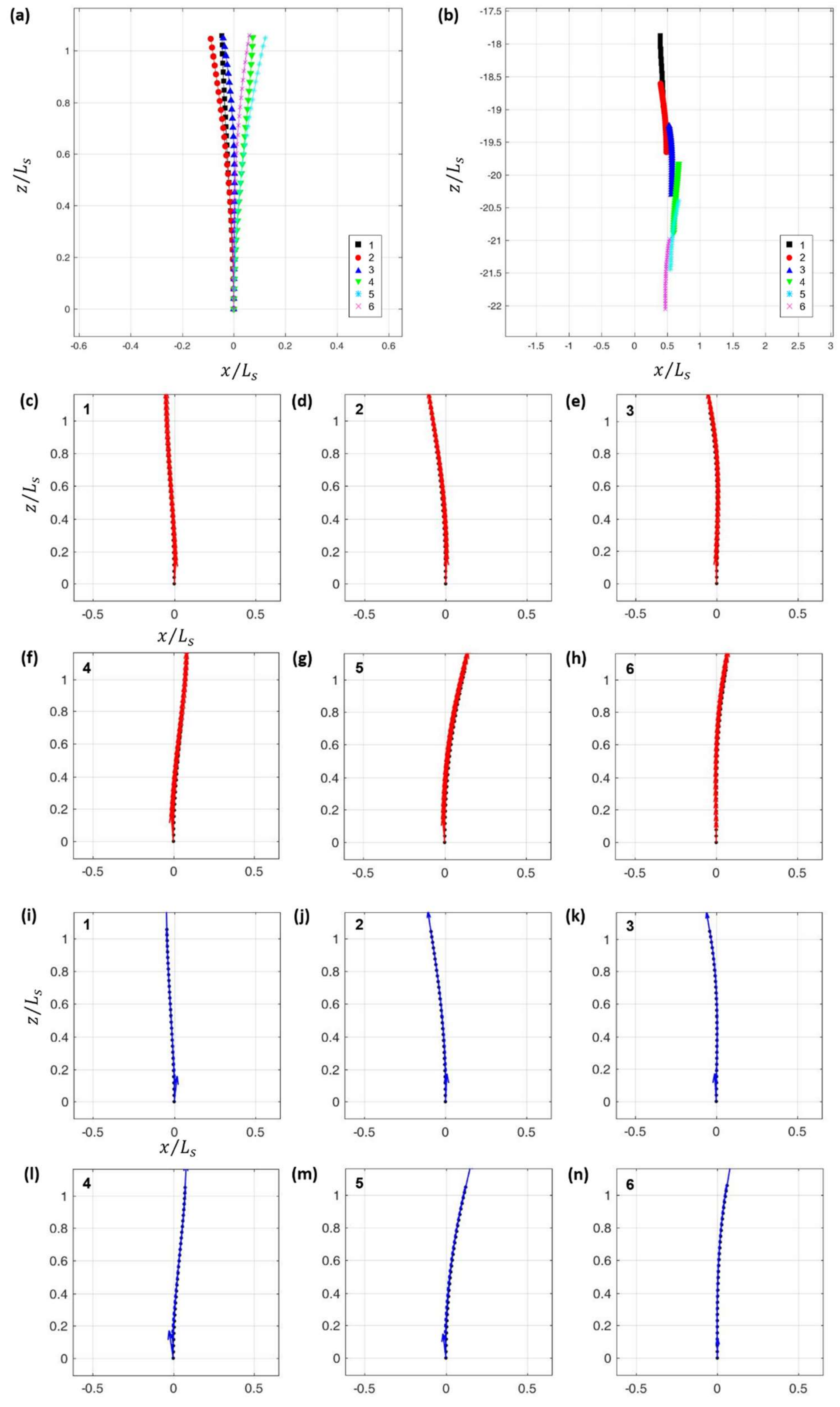


Figure S16. Eddy crossing $\langle V_z \rangle / V_{z,still} = 1.001$; (a) Shape of a filament during sedimentation in relative coordinate of the lower end. (b) Shape of a filament during sedimentation in absolute coordinate. (c)-(h): Distribution of the relative velocity vectors of the flow. (i)-(n): Distribution of the force vectors by the flow. $Bn_{vor} = 0.13$ (the number of beads: 30, length: 4.45 m, mass: 1 kg, viscosity: 0.5 kg/ms, vortex velocity: $U_0 = 0.5\ m/s$, size of the vortex cell: $L_c/L_s = 2$)

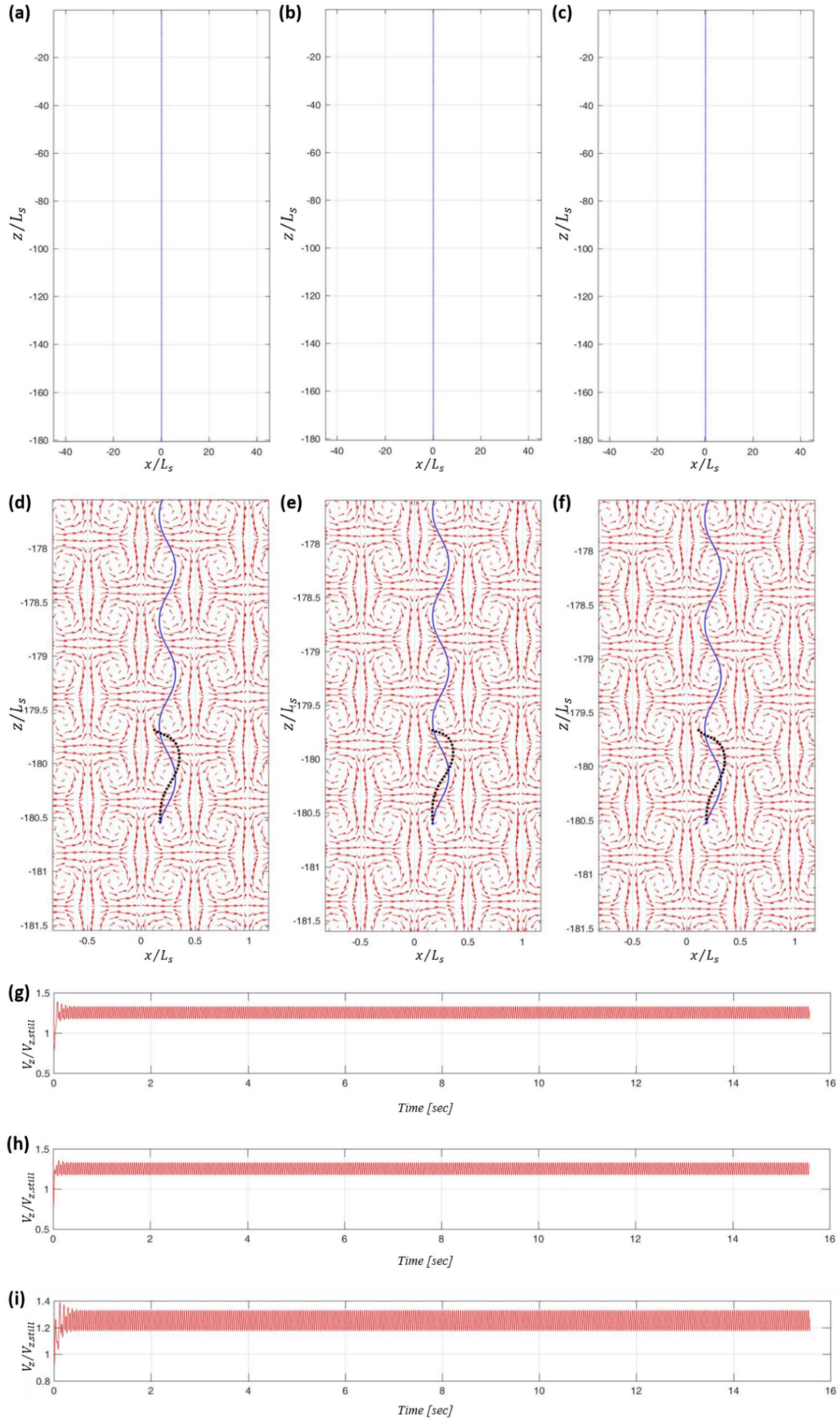

Figure S17. Fast settling $\langle V_z\rangle/V_{z,still} = 1.251$; (a)-(c): Trace plots of a point mass (spider body). (d)-(f): Flow fields and the bead-spring model. (blue lines: trajectory plots). (g)-(i): Nondimensionalised settling speeds over time. $Bn_{vor} = 0.58$ (the number of beads: 30, length: 4.45 m, mass: 0.2 kg, viscosity: 0.5 kg/ms, vortex velocity: $U_0 = 0.5\,m/s$, size of the vortex cell: $L_c/L_s = 0.5$; (a),(d),(g): $x_0/L_c = 0.25$; (b),(e),(h): $x_0/L_c = 0.5$; (c),(f),(i): $x_0/L_c = 0.75$)

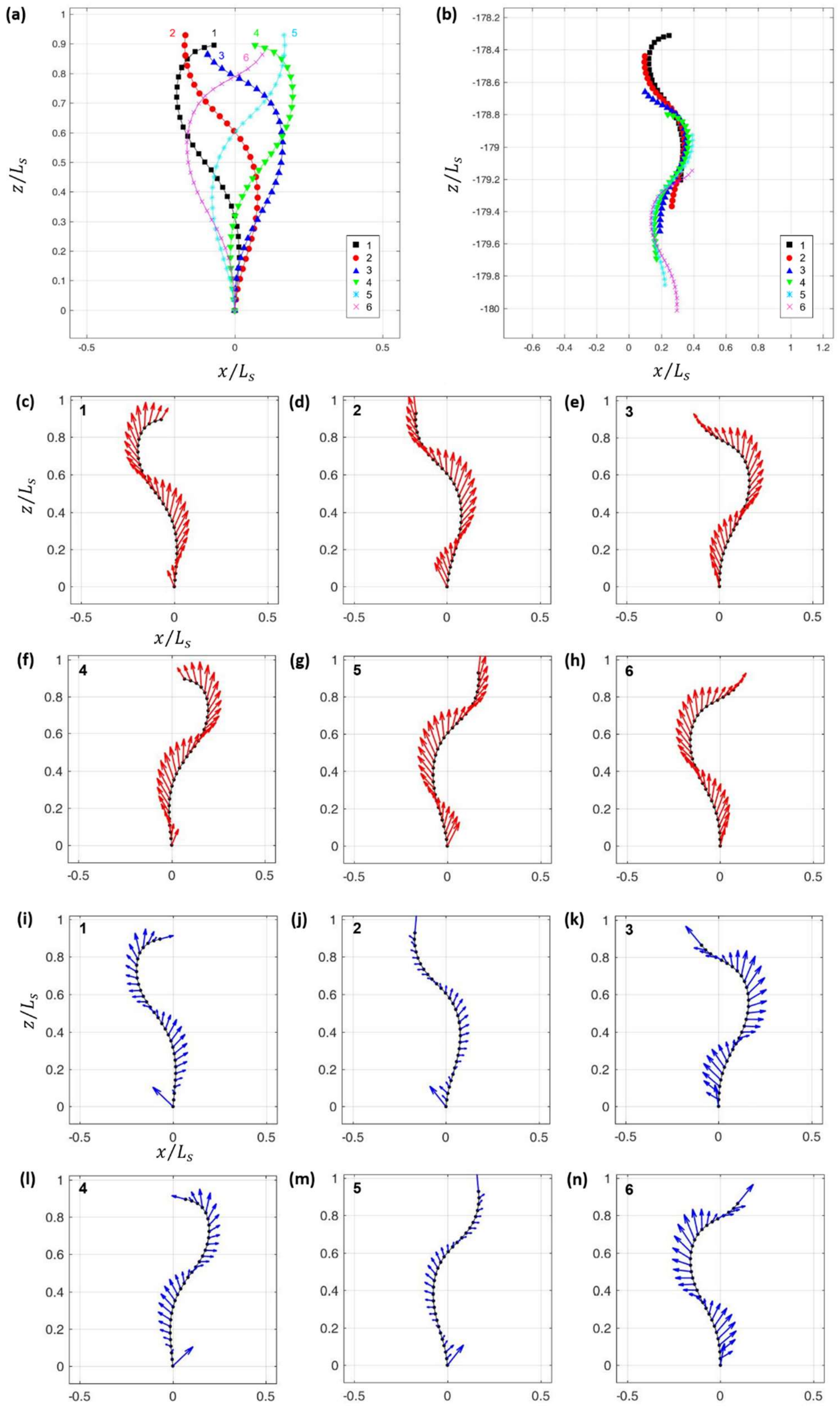


Figure S18. Fast settling $\langle V_z \rangle / V_{z,still} = 1.251$; (a) Shape of a filament during sedimentation in relative coordinate of the lower end. (b) Shape of a filament during sedimentation in absolute coordinate. (c)-(h): Distribution of the relative velocity vectors of the flow. (i)-(n): Distribution of the force vectors by the flow. $Bn_{vor} = 0.58$ (the number of beads: 30, length: 4.45 m, mass: 0.2 kg, viscosity: 0.5 kg/ms, vortex velocity: $U_0 = 0.5\ m/s$, size of the vortex cell: $L_c/L_s = 0.5$)

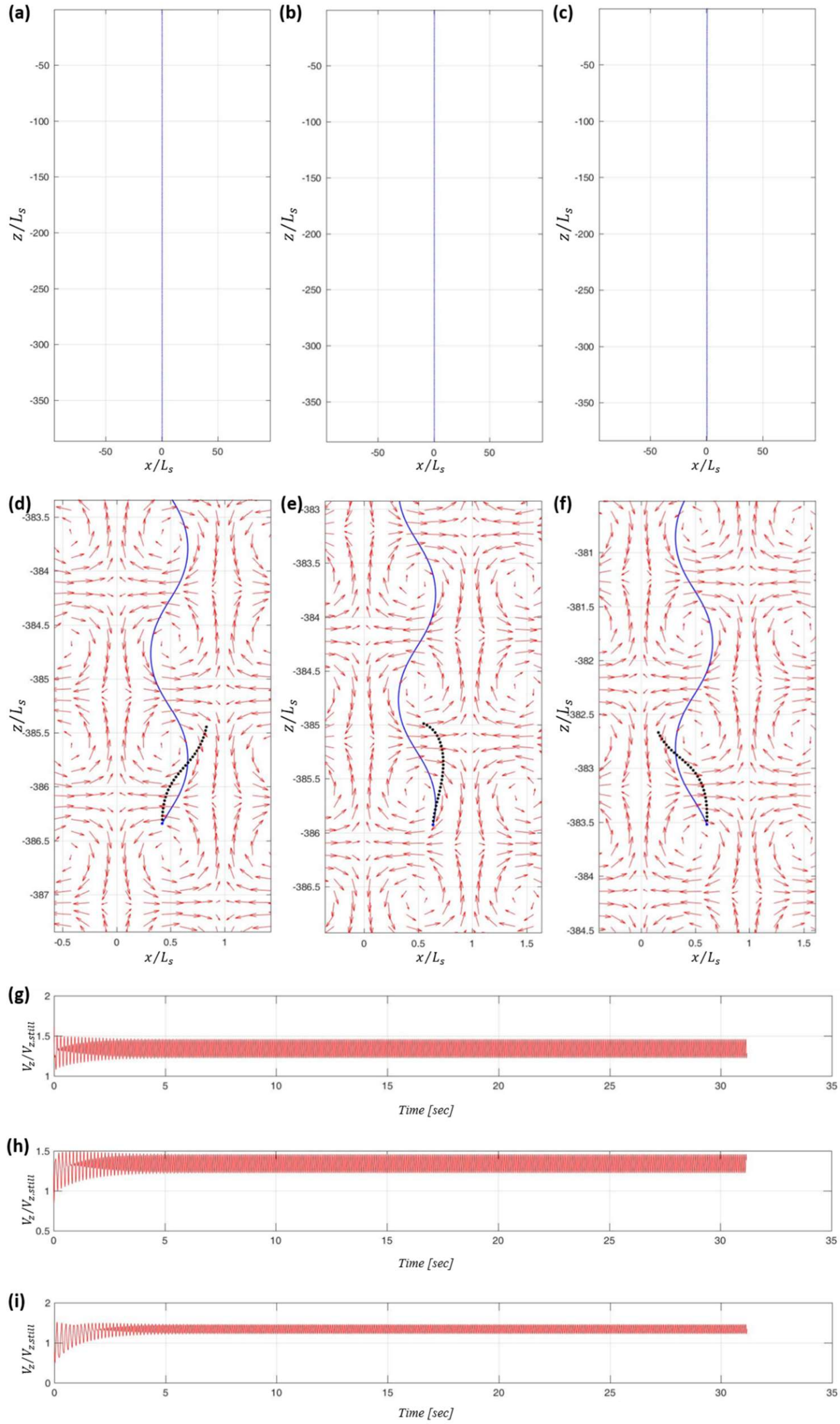


Figure S19. Fast settling $\langle V_z \rangle / V_{z,still} = 1.328$; (a)-(c): Trace plots of a point mass (spider body). (d)-(f): Flow fields and the bead-spring model. (blue lines: trajectory plots). (g)-(i): Nondimensionalised settling speeds over time. $Bn_{vor} = 0.58$ (the number of beads: 30, length: 4.45 m, mass: 0.2 kg, viscosity: 0.5 kg/ms, vortex velocity: $U_0 = 0.5\, m/s$, size of the vortex cell: $L_c/L_s = 1$; (a),(d),(g): $x_0/L_c = 0.25$; (b),(e),(h): $x_0/L_c = 0.5$; (c),(f),(i): $x_0/L_c = 0.75$)

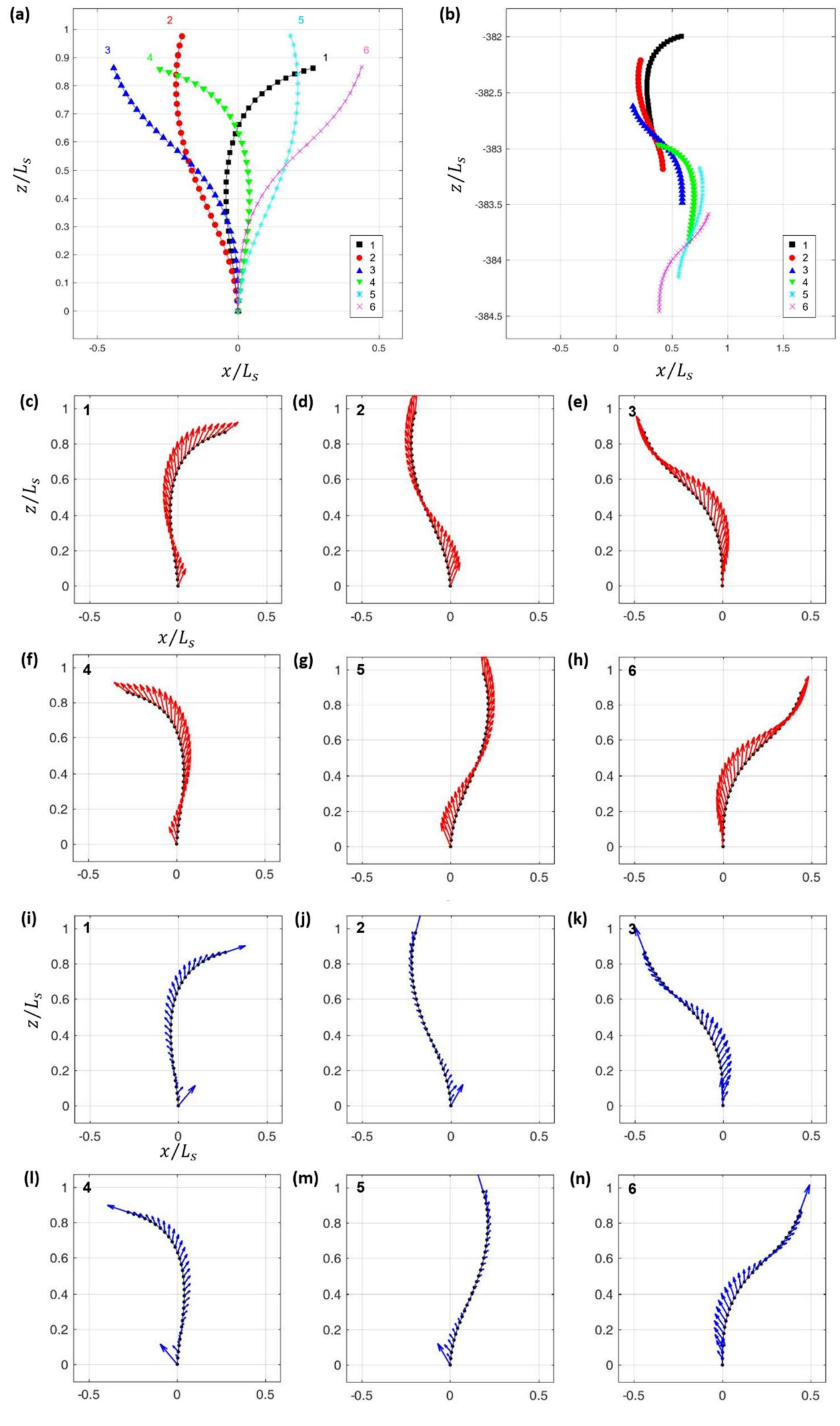


Figure S20. Fast settling $\langle V_z \rangle / V_{z,still} = 1.328$; (a) Shape of a filament during sedimentation in relative coordinate of the lower end. (b) Shape of a filament during sedimentation in absolute coordinate. (c)-(h): Distribution of the relative velocity vectors of the flow. (i)-(n): Distribution of the force vectors by the flow. $Bn_{vor} = 0.58$ (the number of beads: 30, length: 4.45 m, mass: 0.2 kg, viscosity: 0.5 kg/ms, vortex velocity: $U_0 = 0.5\ m/s$, size of the vortex cell: $L_c/L_s = 1$)

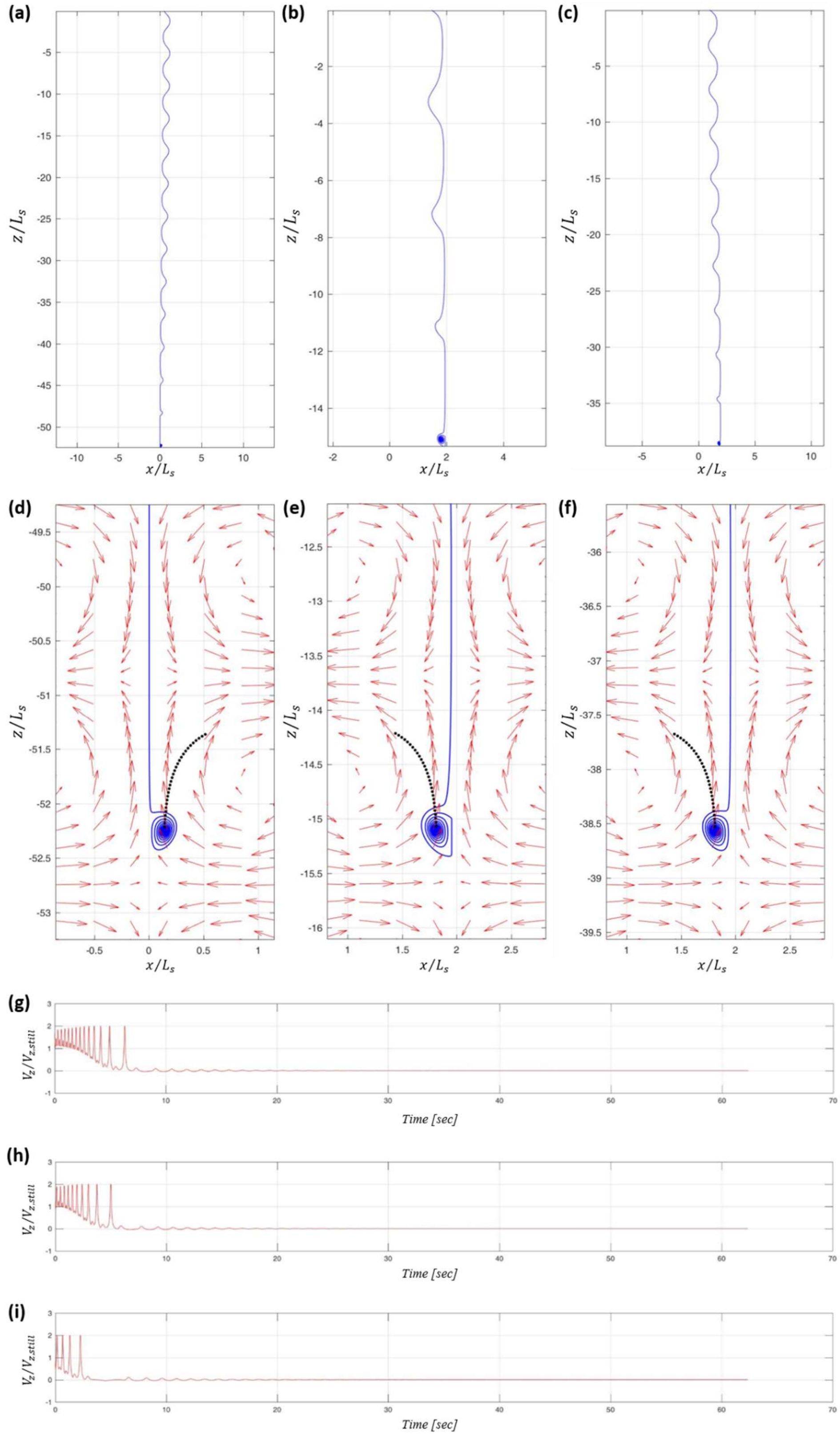


Figure S21. Trapping $\langle V_z \rangle / V_{z,still} = 0$; (a)-(c): Trace plots of a point mass (spider body). (d)-(f): Flow fields and the bead-spring model. (blue lines: trajectory plots). (g)-(i): Nondimensionalised settling speeds over time. $Bn_{vor} = 0.58$ (the number of beads: 30, length: 4.45 m, mass: 0.2 kg, viscosity: 0.5 kg/ms, vortex velocity: $U_0 = 0.5\ m/s$, size of the vortex cell: $L_c/L_s = 2$; (a),(d),(g): $x_0/L_c = 0.25$; (b),(e),(h): $x_0/L_c = 0.5$; (c),(f),(i): $x_0/L_c = 0.75$)

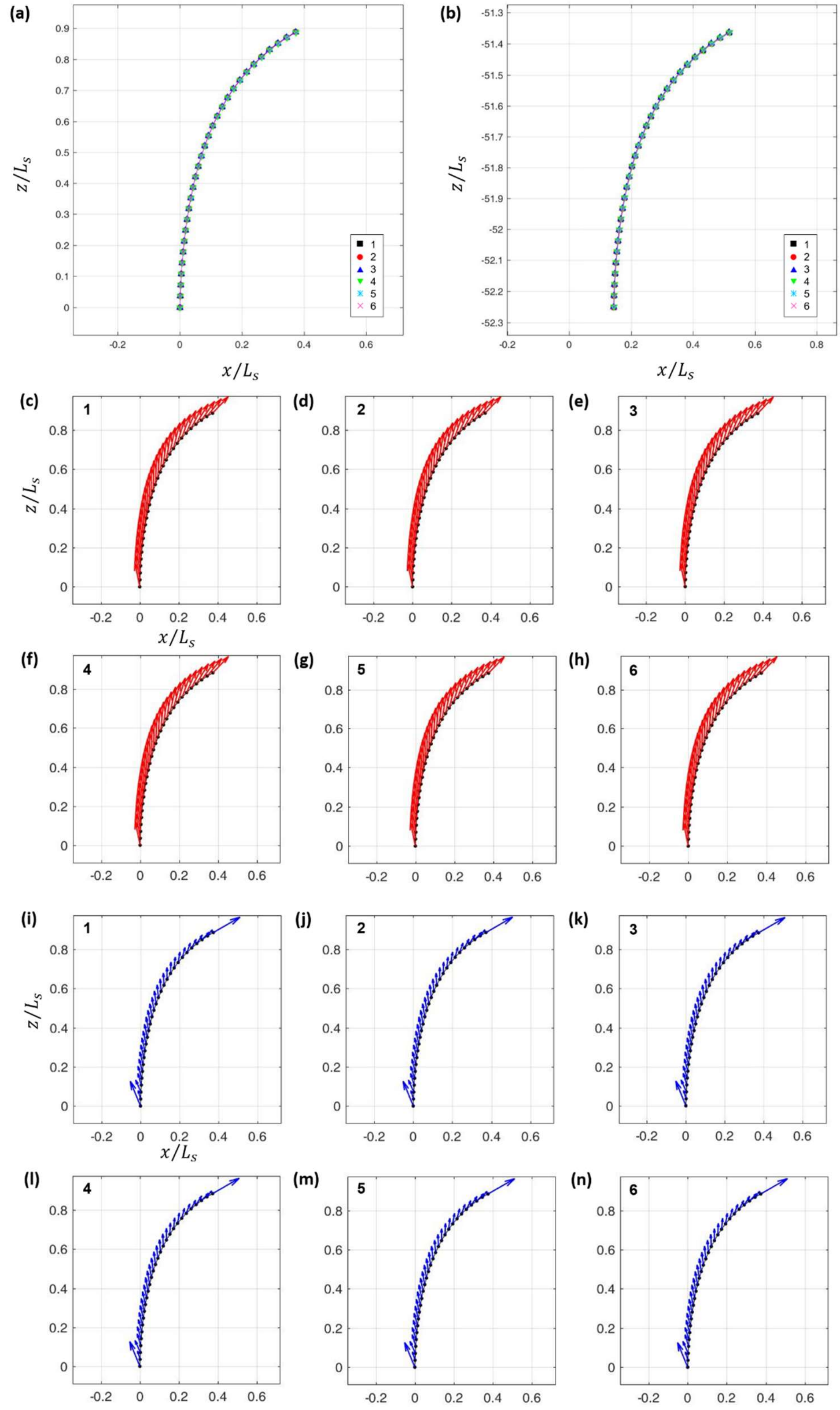


Figure S22. Trapping $\langle V_z \rangle / V_{z,still} = 0$; (a) Shape of a filament during sedimentation in relative coordinate of the lower end. (b) Shape of a filament during sedimentation in absolute coordinate. (c)-(h): Distribution of the relative velocity vectors of the flow. (i)-(n): Distribution of the force vectors by the flow. $Bn_{vor} = 0.58$ (the number of beads: 30, length: 4.45 m, mass: 0.2 kg, viscosity: 0.5 kg/ms, vortex velocity: $U_0 = 0.5\ m/s$, size of the vortex cell: $L_c/L_s = 2$)

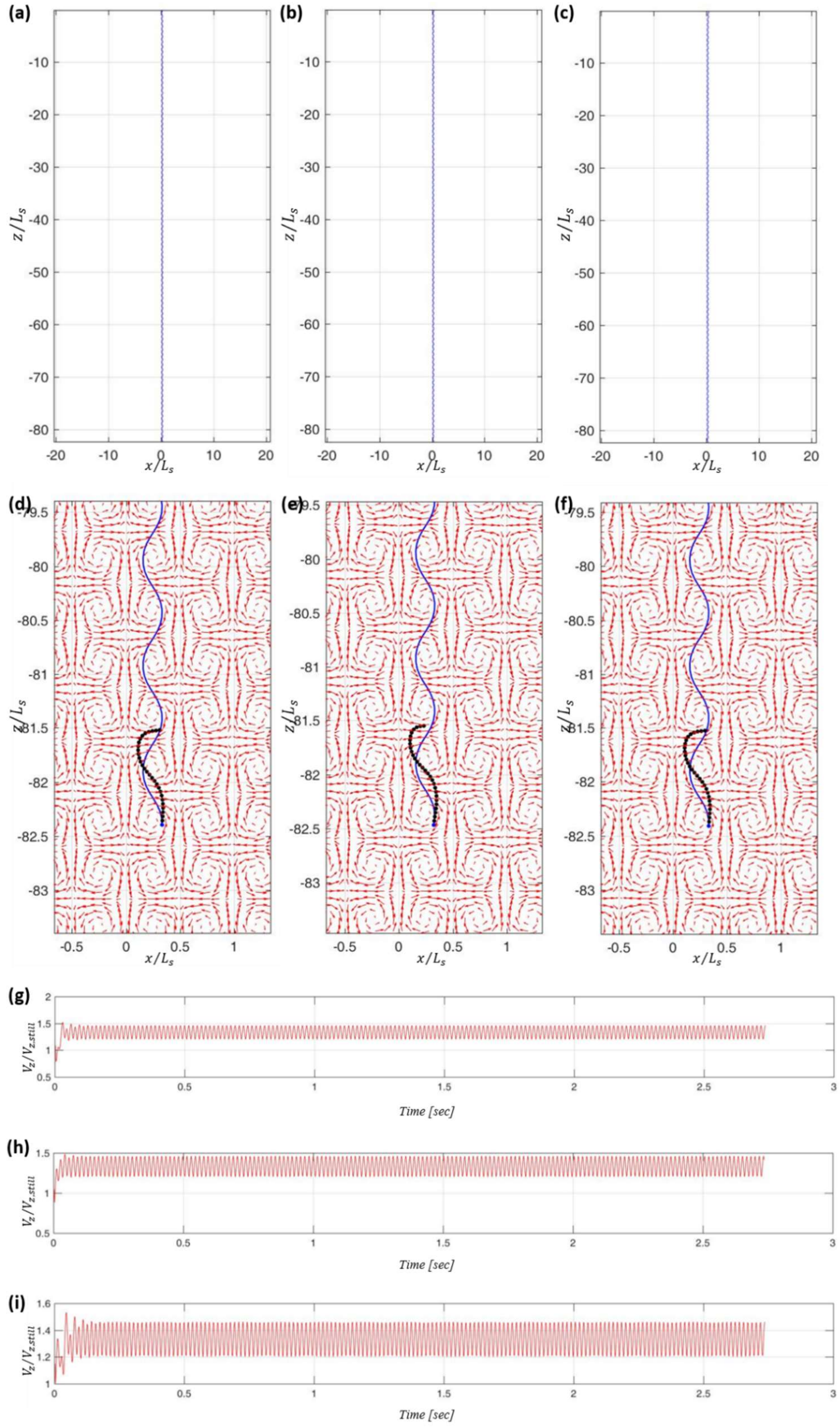

Figure S23. Fast settling $\langle V_z \rangle / V_{z,still} = 1.335$; (a)-(c): Trace plots of a point mass (spider body). (d)-(f): Flow fields and the bead-spring model. (blue lines: trajectory plots). (g)-(i): Nondimensionalised settling speeds over time. $Bn_{vor} = 0.72$ (the number of beads: 30, length: 4.45 m, mass: 0.5 kg, viscosity: 0.5 kg/ms, vortex velocity: $U_0 = 1.5\, m/s$, size of the vortex cell: $L_c/L_s = 0.5$; (a),(d),(g): $x_0/L_c = 0.25$; (b),(e),(h): $x_0/L_c = 0.5$; (c),(f),(i): $x_0/L_c = 0.75$)

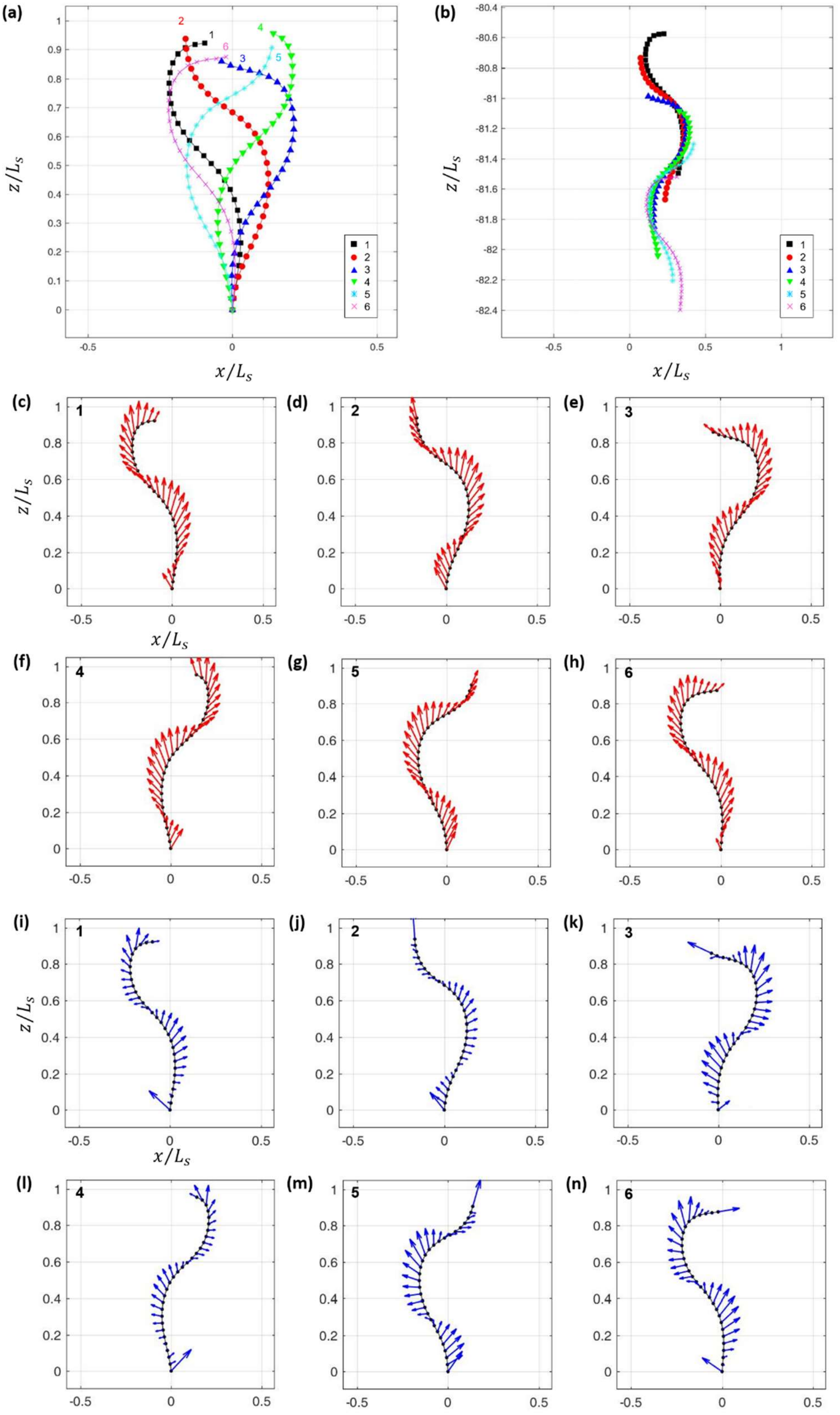


Figure S24. Fast settling $\langle V_z \rangle / V_{z,still} = 1.335$; (a) Shape of a filament during sedimentation in relative coordinate of the lower end. (b) Shape of a filament during sedimentation in absolute coordinate. (c)-(h): Distribution of the relative velocity vectors of the flow. (i)-(n): Distribution of the force vectors by the flow. $Bn_{vor} = 0.72$ (the number of beads: 30, length: 4.45 m, mass: 0.5 kg, viscosity: 0.5 kg/ms, vortex velocity: $U_0 = 1.5\ m/s$, size of the vortex cell: $L_c/L_s = 0.5$)

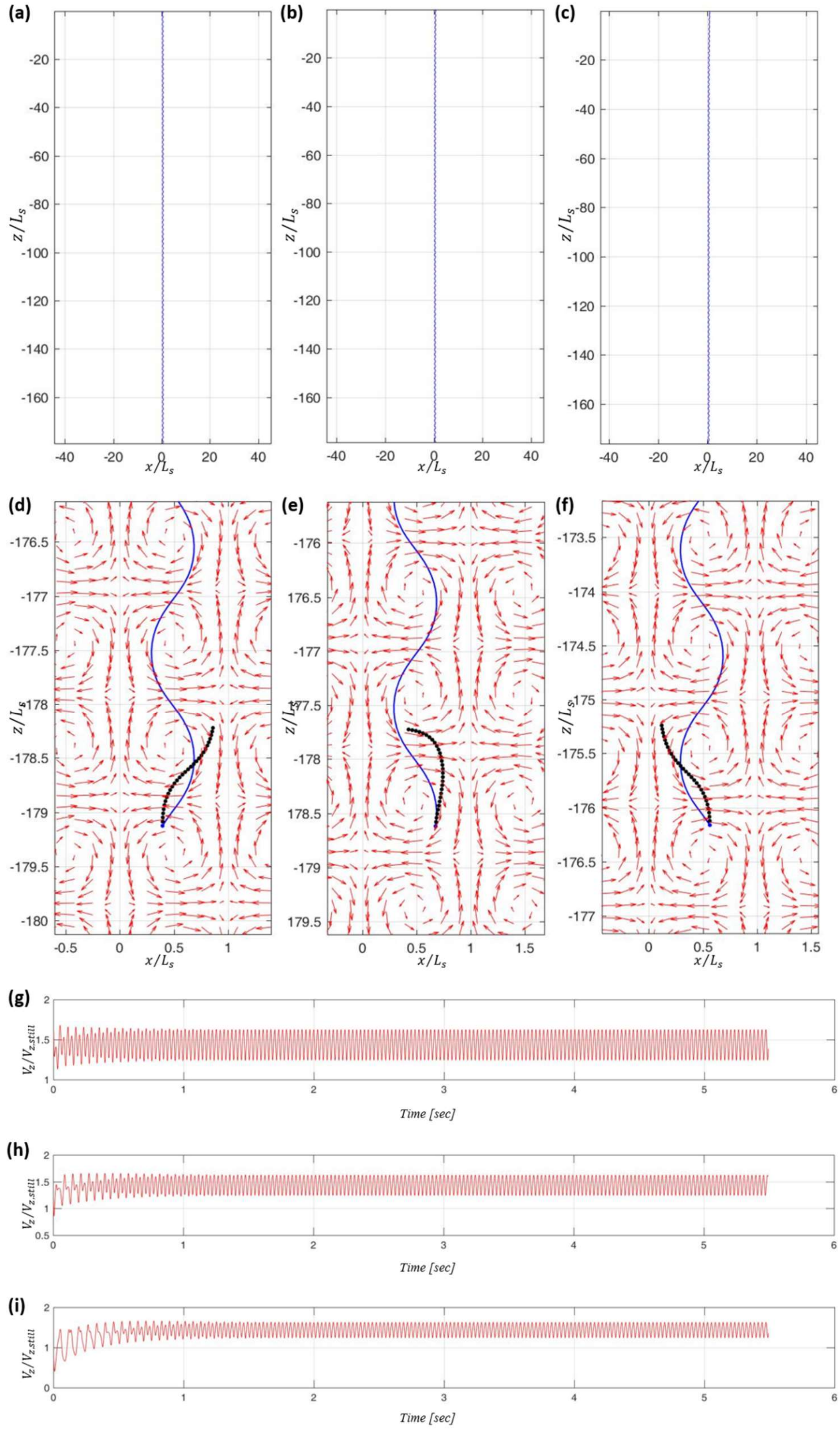


Figure S25. Fast settling $\langle V_z \rangle / V_{z,still} = 1.446$; (a)-(c): Trace plots of a point mass (spider body). (d)-(f): Flow fields and the bead-spring model. (blue lines: trajectory plots). (g)-(i): Nondimensionalised settling speeds over time. $Bn_{vor} = 0.72$ (the number of beads: 30, length: 4.45 m, mass: 0.5 kg, viscosity: 0.5 kg/ms, vortex velocity: $U_0 = 1.5\, m/s$, size of the vortex cell: $L_c/L_s = 1$; (a),(d),(g): $x_0/L_c = 0.25$; (b),(e),(h): $x_0/L_c = 0.5$; (c),(f),(i): $x_0/L_c = 0.75$)

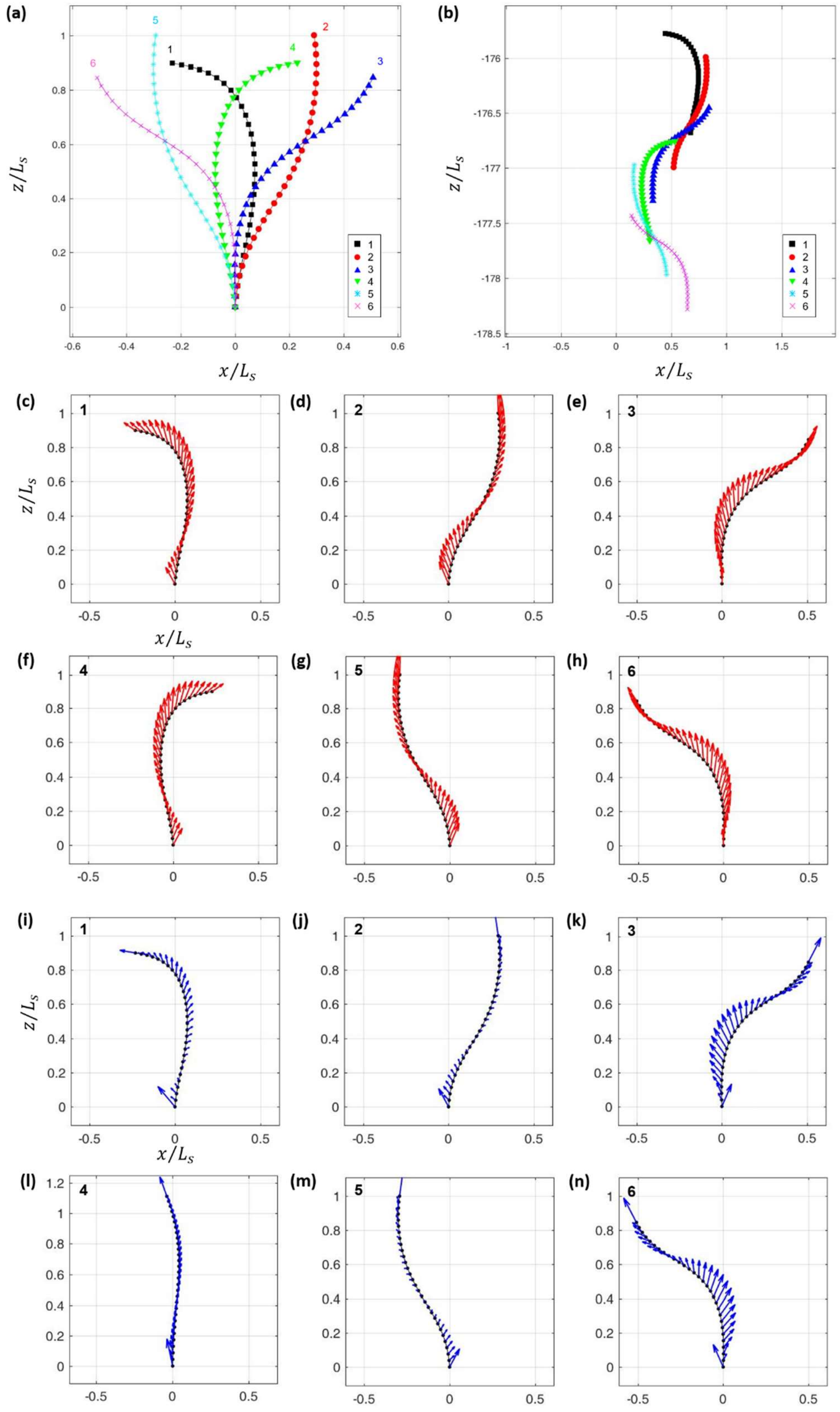

Figure S26. Fast settling $\langle V_z \rangle / V_{z,still} = 1.446$; (a) Shape of a filament during sedimentation in relative coordinate of the lower end. (b) Shape of a filament during sedimentation in absolute coordinate. (c)-(h): Distribution of the relative velocity vectors of the flow. (i)-(n): Distribution of the force vectors by the flow. $Bn_{vor} = 0.72$ (the number of beads: 30, length: 4.45 m, mass: 0.5 kg, viscosity: 0.5 kg/ms, vortex velocity: $U_0 = 1.5\ m/s$, size of the vortex cell: $L_c/L_s = 1$)

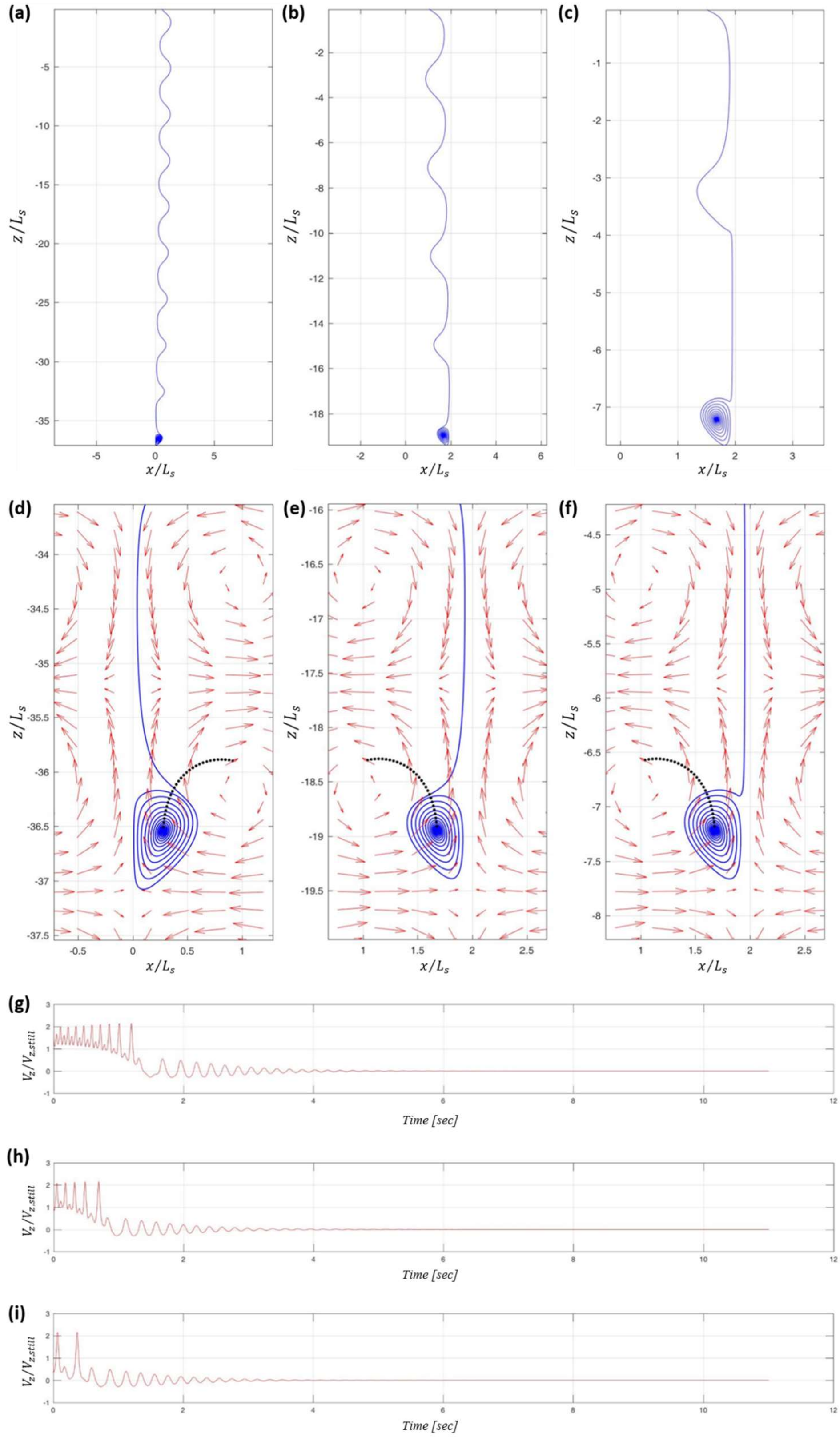


Figure S27. Trapping $\langle V_z \rangle / V_{z,still} = 0$; (a)-(c): Trace plots of a point mass (spider body). (d)-(f): Flow fields and the bead-spring model. (blue lines: trajectory plots). (g)-(i): Nondimensionalised settling speeds over time. $Bn_{vor} = 0.72$ (the number of beads: 30, length: 4.45 m, mass: 0.5 kg, viscosity: 0.5 kg/ms, vortex velocity: $U_0 = 1.5\ m/s$, size of the vortex cell: $L_c/L_s = 2$; (a),(d),(g): $x_0/L_c = 0.25$; (b),(e),(h): $x_0/L_c = 0.5$; (c),(f),(i): $x_0/L_c = 0.75$)

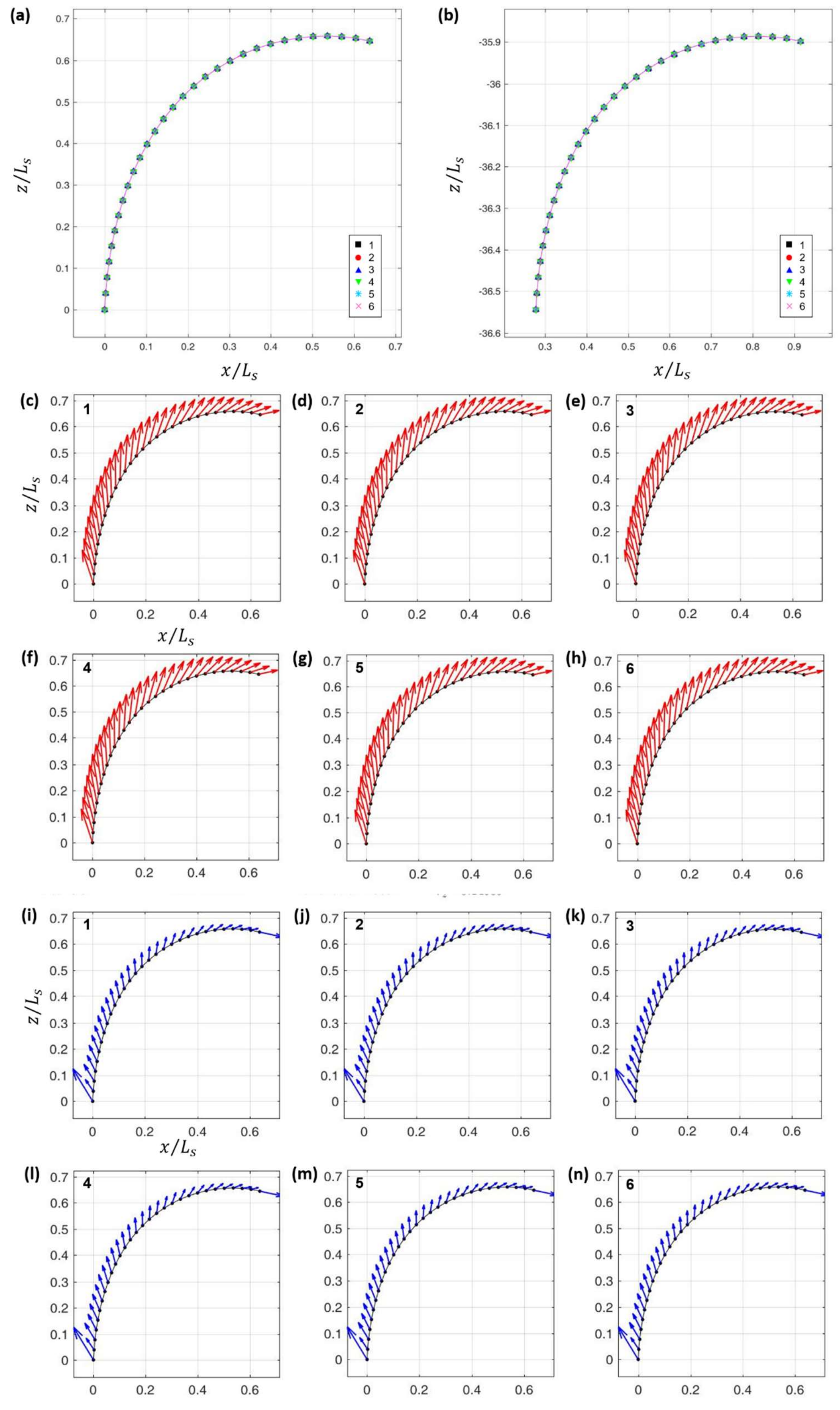


Figure S28. Trapping $\langle V_z \rangle / V_{z,still} = 0$; (a) Shape of a filament during sedimentation in relative coordinate of the lower end. (b) Shape of a filament during sedimentation in absolute coordinate. (c)-(h): Distribution of the relative velocity vectors of the flow. (i)-(n): Distribution of the force vectors by the flow. $Bn_{vor} = 0.72$ (the number of beads: 30, length: 4.45 m, mass: 0.5 kg, viscosity: 0.5 kg/ms, vortex velocity: $U_0 = 1.5\ m/s$, size of the vortex cell: $L_c/L_s = 2$)

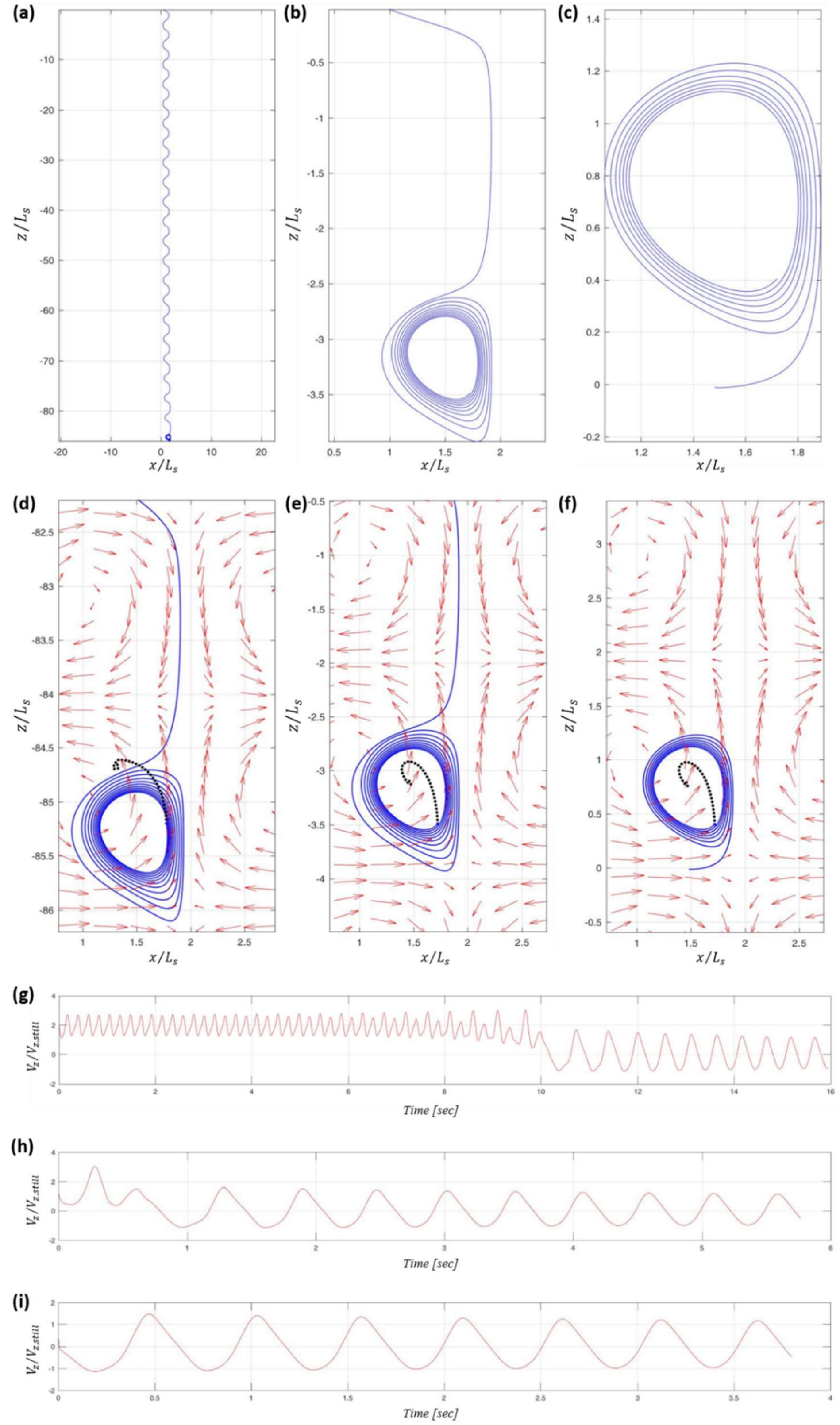


Figure S29. Trapping $\langle V_z \rangle / V_{z,still} = 0$; (a)-(c): Trace plots of a point mass (spider body). (d)-(f): Flow fields and the bead-spring model. (blue lines: trajectory plots). (g)-(i): Nondimensionalised settling speeds over time. $Bn_{vor} = 1.13$ (the number of beads: 30, length: 4.45 m, mass: 0.1 kg, viscosity: 0.5 kg/ms, vortex velocity: $U_0 = 0.5\, m/s$, size of the vortex cell: $L_c/L_s = 2$; (a),(d),(g): $x_0/L_c = 0.25$; (b),(e),(h): $x_0/L_c = 0.5$; (c),(f),(i): $x_0/L_c = 0.75$)

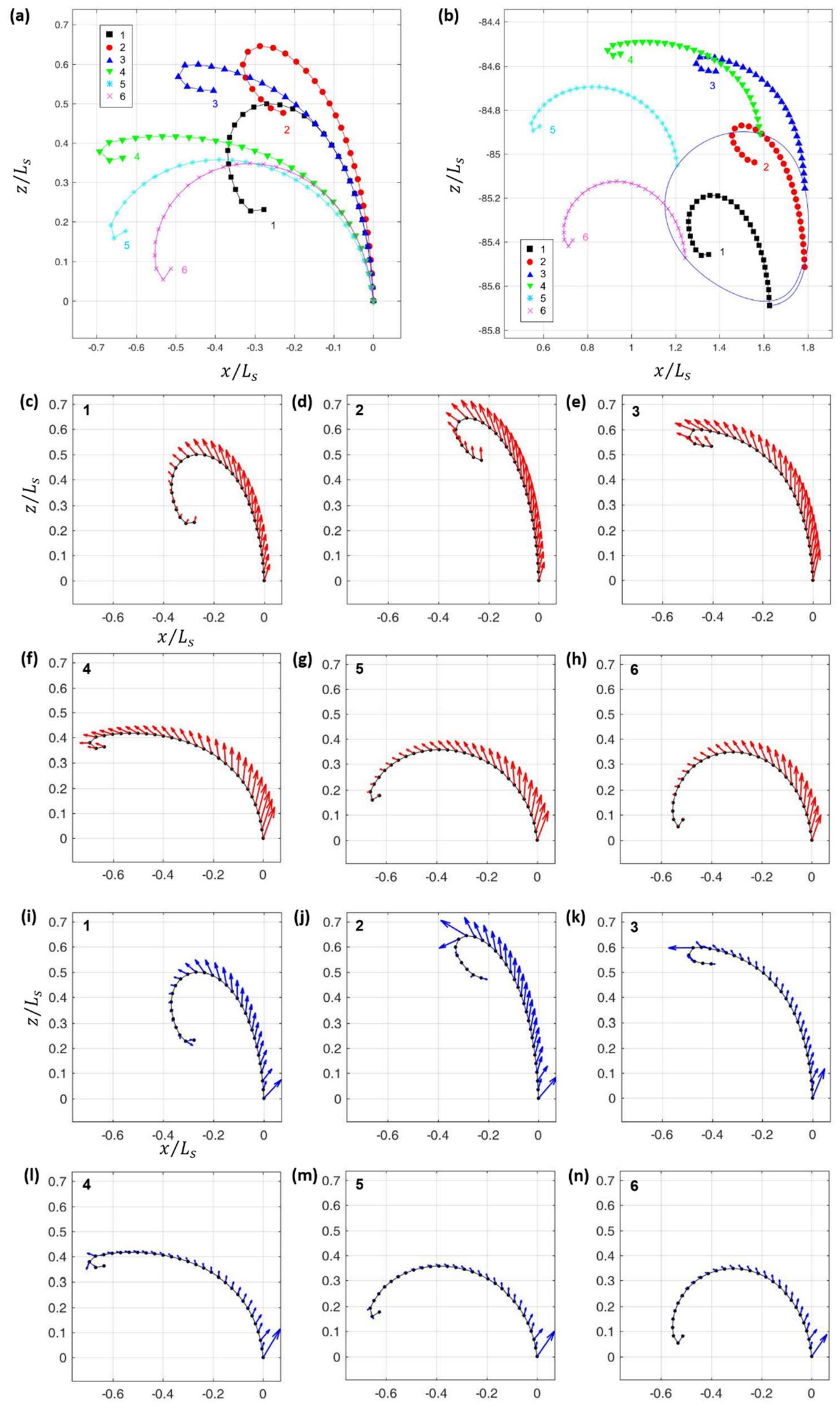


Figure S30. Trapping $\langle V_z \rangle / V_{z,still} = 0$; (a) Shape of a filament during sedimentation in relative coordinate of the lower end. (b) Shape of a filament during sedimentation in absolute coordinate. (c)-(h): Distribution of the relative velocity vectors of the flow. (i)-(n): Distribution of the force vectors by the flow. $Bn_{vor} = 1.13$ (the number of beads: 30, length: 4.45 m, mass: 0.1 kg, viscosity: 0.5 kg/ms, vortex velocity: $U_0 = 0.5\ m/s$, size of the vortex cell: $L_c/L_s = 2$)

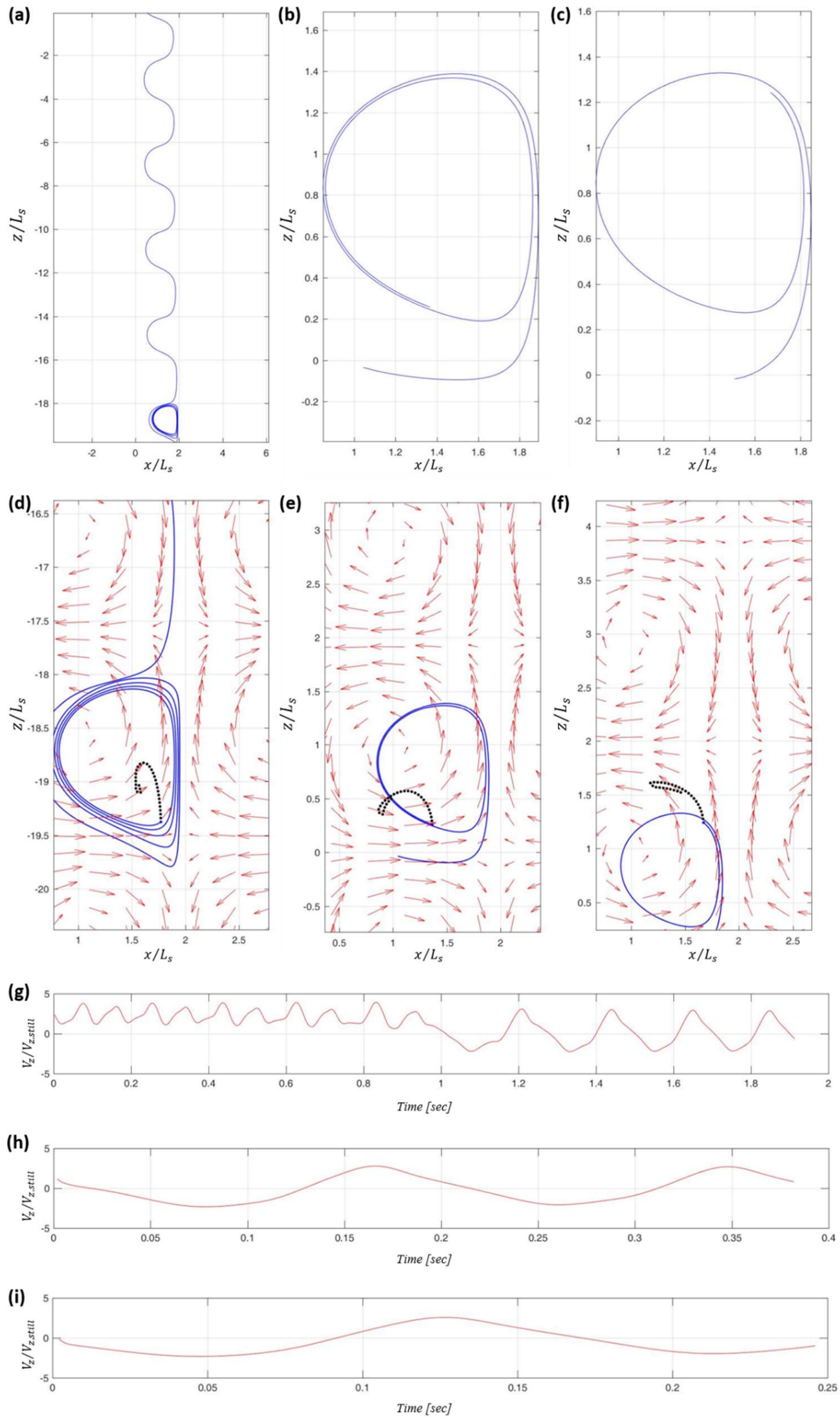


Figure S31. Trapping $\langle V_z \rangle / V_{z,still} = 0$; (a)-(c): Trace plots of a point mass (spider body). (d)-(f): Flow fields and the bead-spring model. (blue lines: trajectory plots). (g)-(i): Nondimensionalised settling speeds over time. $Bn_{vor} = 1.72$ (the number of beads: 30, length: 4.45 m, mass: 0.2 kg, viscosity: 0.5 kg/ms, vortex velocity: $U_0 = 1.5\, m/s$, size of the vortex cell: $L_c/L_s = 2$; (a),(d),(g): $x_0/L_c = 0.25$; (b),(e),(h): $x_0/L_c = 0.5$; (c),(f),(i): $x_0/L_c = 0.75$)

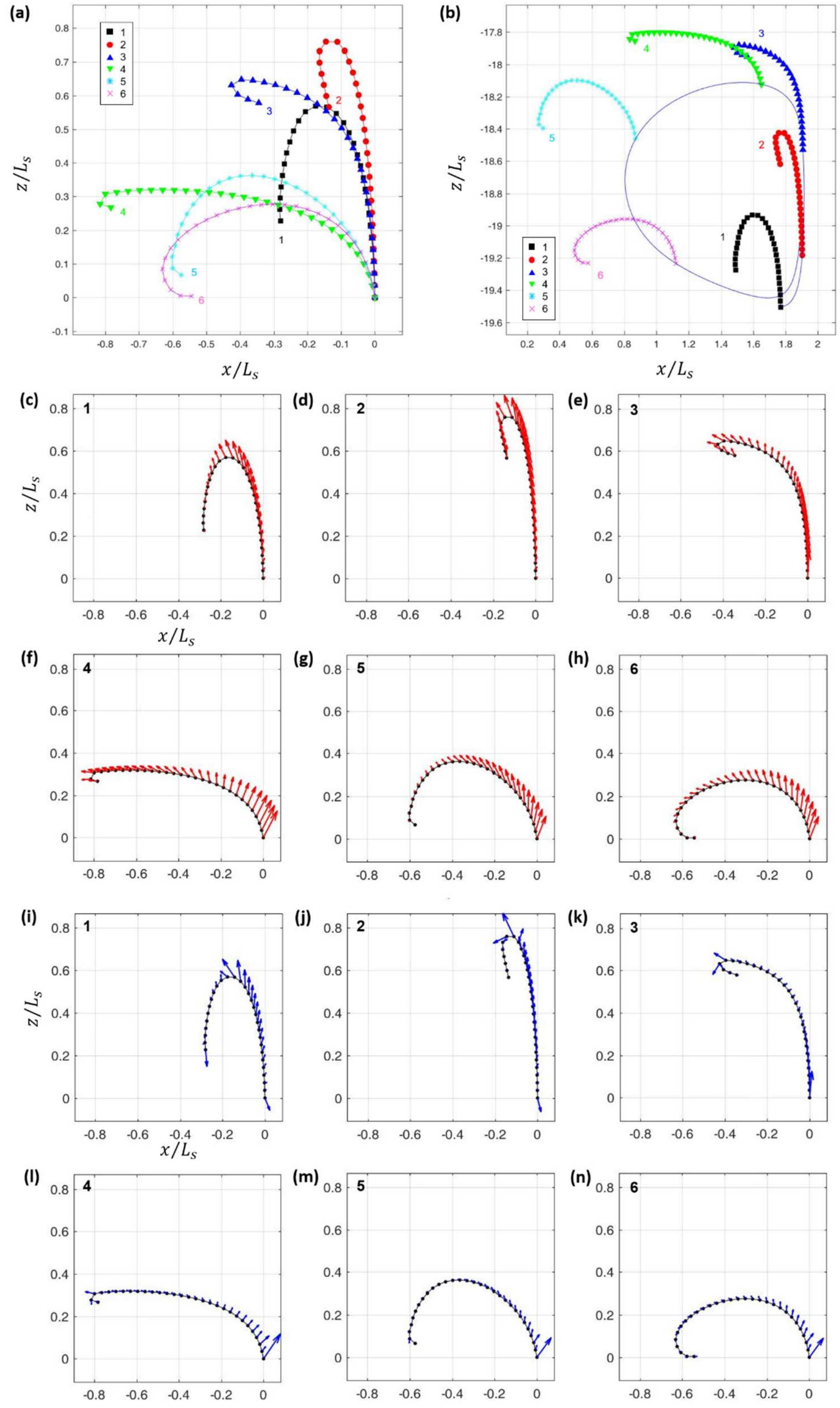


Figure S32. Trapping $\langle V_z \rangle / V_{z,still} = 0$; (a) Shape of a filament during sedimentation in relative coordinate of the lower end. (b) Shape of a filament during sedimentation in absolute coordinate. (c)-(h): Distribution of the relative velocity vectors of the flow. (i)-(n): Distribution of the force vectors by the flow. $Bn_{vor} = 1.72$ (the number of beads: 30, length: 4.45 m, mass: 0.2 kg, viscosity: 0.5 kg/ms, vortex velocity: $U_0 = 1.5\ m/s$, size of the vortex cell: $L_c/L_s = 2$)

#### List of symbols and abbreviation

| | |
|---|---|
| $A$ | bending stiffness |
| $a$ | radius of a bead |
| $a_i$ | radius of the *i*-th bead |
| $a_j$ | radius of the *j*-th bead |
| $\boldsymbol{a}_n$ | amplitude vector of the *n*-th mode |
| $\boldsymbol{b}_n$ | amplitude vector of the *n*-th mode |
| $Bn_{shear}$ | ballooning number for a shear flow |
| $Bn_{tur}$ | ballooning number for a turbulent flow |
| $Bn_{vor}$ | ballooning number for a vortex flow |
| $d$ | diameter of a fiber (the unit is meter in Suter's formula) |
| $E(k)$ | dimensionless form of energy spectrum function |
| $E^B$ | total bending free energy |
| $E^S$ | total stretching free energy |
| $F$ | fluid dynamic force |
| $f_i$ | factor for the consideration of free end condition of the beads |
| $F_N$ | drag force in the direction normal to the axis of the cylinder (or a straight fiber) |
| $F_T$ | drag force in the direction parallel to the axis of the cylinder (or a straight fiber) |
| $f_{tur}$ | factor of the total integration time for the simulation in a homogeneous turbulence |
| $f_{vor}$ | factor of the total integration time for the simulation in a periodic cellular flow |
| $F_z$ | resistance force of an object during sedimentation in a low Reynolds number flow |
| $\boldsymbol{F}_i^B$ | a bending force on the *i*-th bead in a bead-spring model |
| $\boldsymbol{F}_i^F$ | a fluid-dynamic force on the *i*-th bead in a bead-spring model |
| $\boldsymbol{F}_i^H$ | fluid-dynamic force on the *i*-th bead, which is induced by the horizontal component of the relative flow velocity (shear-induced fluid dynamic force on the *i*-th bead) |
| $\boldsymbol{F}_i^G$ | a gravity force on the *i*-th bead in a bead-spring model |
| $\boldsymbol{F}_i^S$ | a stretching force on the *i*-th bead in a bead-spring model |
| $\boldsymbol{F}_i^{total}$ | a net force on the *i*-th bead in a bead-spring model |
| $\boldsymbol{F}_i^V$ | fluid-dynamic force on the *i*-th bead, which is induced by the vertical component of the relative flow velocity (geometric fluid dynamic force on the *i*-th bead) |
| $F_z^H$ | vertical component of net shear-induced fluid-dynamic force |
| $F_{z,i}^H$ | vertical component of shear-induced fluid-dynamic force on the *i*-th bead |
| $F_z^V$ | vertical component of net geometric fluid-dynamic force |
| $F_{z,i}^V$ | vertical component of geometric fluid-dynamic force on the *i*-th bead |
| $g$ | acceleration of gravity |
| $g_1$ | numerical constant |
| $g_2$ | numerical constant |
| $k$ | wavenumber |
| $k_0$ | elastic constant of a spring |

| | |
|---|---|
| $k_n$ | length of the *n*-th wavenumber vector |
| $\boldsymbol{k}_n$ | wavenumber vector of the *n*-th mode |
| $L$ | length of a fiber (the unit is meter in Suter's formula) |
| $L_c$ | cell size of a periodic cellular flow |
| $l_0$ | length of the unstretched spring between the beads |
| $l_i$ | length of a spring between the i-th bead and the ($i$+$1$)-th bead |
| $n$ | number of fibers |
| $N_{tur}$ | total number of integration time-<br>steps for the simulation in a homogeneous turbulence |
| $N_{vor}$ | total number of integration time-<br>steps for the simulation in a periodic cellular flow |
| $\boldsymbol{r}_i$ | position vector of the *i*-th bead |
| $\boldsymbol{r}_j$ | position vector of the *j*-th bead |
| $\boldsymbol{r}_{ij}$ | position vector of the *j*-th bead from the local coordinate of the *i*-th bead |
| $\hat{\boldsymbol{r}}_{ij}$ | unit vector of the vector $\boldsymbol{r}_{ij}$ |
| $R_N$ | non-dimensional resistance coefficient in the direction normal to the axis of the cylinder (or a straight fiber) |
| $R_T$ | non-dimensional resistance coefficient in the direction parallel to the axis of the cylinder (or a straight fiber) |
| $\boldsymbol{t}_i$ | link vector between the i-th bead and the ($i$+$1$)-th bead |
| $\hat{\boldsymbol{t}}_i$ | unit vector of the vector $\boldsymbol{t}_i$ |
| $TKE$ | turbulence kinetic energy |
| $U$ | flow velocity (the unit is meter per second in Suter's formula); characteristic flow velocity |
| $\boldsymbol{u}^{\infty}$ | background flow velocity field |
| $U_0$ | maximum flow velocity in a periodic cellular flow |
| $v_i$ | volume of the *i*-th bead |
| $V_z$ | sediment velocity (z-direction) of the first bead (body) |
| $V_{z,still}$ | sediment velocity (z-direction) of the first bead (body) in the still air |
| $w_i$ | weight of the *i*-th bead |
| $x_0$ | horizontal coordinate of the release position of the body |
| $\Delta t$ | time step |
| $\lambda$ | shear rate |
| $\eta$ | dynamic viscosity of fluid |
| $\boldsymbol{\mu}_{ij}$ | bead mobility tensor |
| $\rho_{air}$ | density of the air |
| $\rho_f$ | density of a fluid |
| $\rho_i$ | density of the *i*-th bead |
| $\sigma$ | the root-mean-square fluctuation velocity of a homogeneous turbulence |

### References

Bonino, Mark J. (2003). Material Properties of Spider Silk. Degree of Master of Science. University of Rochester, Rochester, NY.

Burgers, J. M. (1938): in Second Report on Viscosity and Plasticity: Noord-Hollandsche Uitgevers-Maatschappij (Verhandelingen der Koninklijke Akademie van Wetenschappen, Afdeeling Natuurkunde: Wiskunde, natuurkunde, scheikunde, kristallenleer, sterrenkunde, weerkunde, en ingenieurswetenschappen).

Cho, Moonsung (2020): Suspension of a Point-Mass-Loaded Filament in Non-Uniform Flows: The Ballooning Flight of Spiders. PhD Thesis. Technical University of Berlin Repository

Doyle, Patrick S.; Underhill, Patrick T. (2005): Brownian Dynamics Simulations of Polymers and Soft Matter. In Sidney Yip (Ed.): Handbook of materials modeling, vol. 19. Dordrecht: Springer, pp. 2619–2630.

Drummond, I. T.; Duane, S.; Horgan, R. R. (1984): Scalar diffusion in simulated helical turbulence with molecular diffusivity. In J. Fluid Mech. 138 (-1), p. 75. DOI: 10.1017/S0022112084000045.

Fung, J. C. H.; Hunt, J. C. R.; Malik, N. A.; Perkins, R. J. (1992): Kinematic simulation of homogeneous turbulence by unsteady random Fourier modes. In J. Fluid Mech. 236 (-1), p. 281. DOI: 10.1017/S0022112092001423.

Gauger, Erik; Stark, Holger (2006): Numerical study of a microscopic artificial swimmer. In Physical review. E, Statistical, nonlinear, and soft matter physics 74 (2 Pt 1), p. 21907. DOI: 10.1103/PhysRevE.74.021907.

Happel, John; Brenner, Howard (1991): Low Reynolds number hydrodynamics. With special applications to particulate media. 5. printing. Dordrecht: Kluwer Acad. Publ (Mechanics of fluids and transport processes).

Hinze, Julius Oscar (1987): Turbulence. 2. ed., reissued. New York, NY: McGraw-Hill (McGraw-Hill series in mechanical engineering).

Hunt, J. C. R. (1973): A theory of turbulent flow round two-dimensional bluff bodies. In J. Fluid Mech. 61 (04), p. 625. DOI: 10.1017/S0022112073000893.

Jeffrey, D. J.; Onishi, Y. (1984): The forces and couples acting on two nearly touching spheres in low-Reynolds-number flow. In Z. angew. Math. Phys. 35 (5), pp. 634–641. DOI: 10.1007/BF00952109.

Ko, Frank K.; Kawabata, Sueo; Inoue, Mari; Niwa, Masako; Fossey, Stephen; Song, John W. (2001): Engineering Properties of Spider Silk. In MRS Proc. 702, p. 91. DOI: 10.1557/PROC-702-U1.4.1.

Kraichnan, Robert H. (1970): Diffusion by a Random Velocity Field. In Phys. Fluids 13 (1), p. 22. DOI: 10.1063/1.1692799.

Lauga, Eric; Powers, Thomas R. (2009): The hydrodynamics of swimming microorganisms. In Rep. Prog. Phys. 72 (9), p. 96601. DOI: 10.1088/0034-4885/72/9/096601.

Purcell, E. M. (1977): Life at low Reynolds number. In American Journal of Physics 45 (1), pp. 3–11. DOI: 10.1119/1.10903.

Turfus C. (1985): Stochastic Modelling of Turbulent Dispersion Near Surfaces: University of Cambridge.
